# Modification of DNA by a viral enzyme and charged tRNA

**DOI:** 10.1101/2023.03.24.534169

**Authors:** Rebekah M. B. Silva, Anton Slyvka, Yan-Jiun Lee, Chudi Guan, Sean R. Lund, Elisabeth A. Raleigh, Krzysztof Skowronek, Matthias Bochtler, Peter R. Weigele

**Affiliations:** Research Department, New England Biolabs, Inc.; Ipswich, MA, USA; International Institute of Molecular and Cell Biology in Warsaw; Warsaw, Poland; Institute of Biochemistry and Biophysics, Polish Academy of Sciences; Warsaw, Poland

## Abstract

Bacteriophage enzymes synthesize varied and complex DNA hypermodifications. The enzyme encoded by the phage Mu gene *mom* is necessary for post-replicative carbamoylmethyl addition to the exocyclic amine of deoxyadenosine in DNA during the lytic phase of the viral life-cycle. The molecular details of this modification reaction, including the molecular origins of the modification itself, have long eluded understanding. Here, we demonstrate that Mom co-opts the translational machinery of the host by harvesting activated glycine from charged tRNA^Gly^ to hypermodify adenine. Based on this insight, we report the first *in vitro* reconstitution of the Mu hypermodification from purified components. Using isotope labeling, we demonstrate that the carbamoyl nitrogen of the Mom modification is derived from the *N*6 of adenine, indicating an on-base rearrangement of the *N*6 aminoacylation product, possibly via a cyclic intermediate. Informed by the X-ray crystal structure of Mom, we have probed the location of the active site, identified a novel insertion, and established substrate specificities of the Mom enzyme.

## Introduction

Bacteriophages synthesize diverse modifications of their genomic DNA to evade host defenses (*1– 5*). Complex nucleobase modifications, termed hypermodifications (named by Ross H. Hall in 1971), stem from cellular metabolites, are introduced pre- and/or post-replicatively, and do not affect the coding capacity of the bases (*3, 4, 6*). One hypermodification of historic significance was found in the genome of phage Mu over 40 years ago and subsequently mapped to a loose SASNY consensus sequence (accounting for 15-25% of total dA) (Methods, fig. S1, and table S1) *(7–12)*. In 1983, McCloskey and coworkers determined the structure of Mu modified dA (Fig. 1A), reporting an unusual <u>c</u>arbamoyl<u>m</u>ethyl (glycinamide) group at the <u>N6</u>-position of adenine (abbreviated as 6-*N*cmdA) (*13*). These foundational studies on Mu also identified a gene, *mom* (<u>mo</u>dification of <u>M</u>u), necessary for the installation of 6-*N*cmdA during late stages of Mu infection (*7–11, 14–17*). More recently, it was established that *mom* is the only phage Mu gene required for hypermodification of dA (fig. S2-S3, referred to colloquially as ‘momylation’) (*16–20*). Protein sequence and structural comparisons showed that Mom belongs to the Gcn5-Related N-Acetyltransferase (GNAT) superfamily (*16, 18–20*), though it remained undetermined whether Mom is an active acyltransferase, and if so, which substrate it transfers (*13, 16, 17*). Here, we demonstrate that phage Mu enzyme Mom co-opts charged tRNA^Gly^ (Gly-tRNA^Gly^) in order to form 6-*N*cmdA. Moreover, the mechanism uncovered herein reveals that the Mom-catalyzed hypermodification is not formed via simple group transfer, helping to explain how this reaction evaded elucidation for so long.

**Figure 1:**
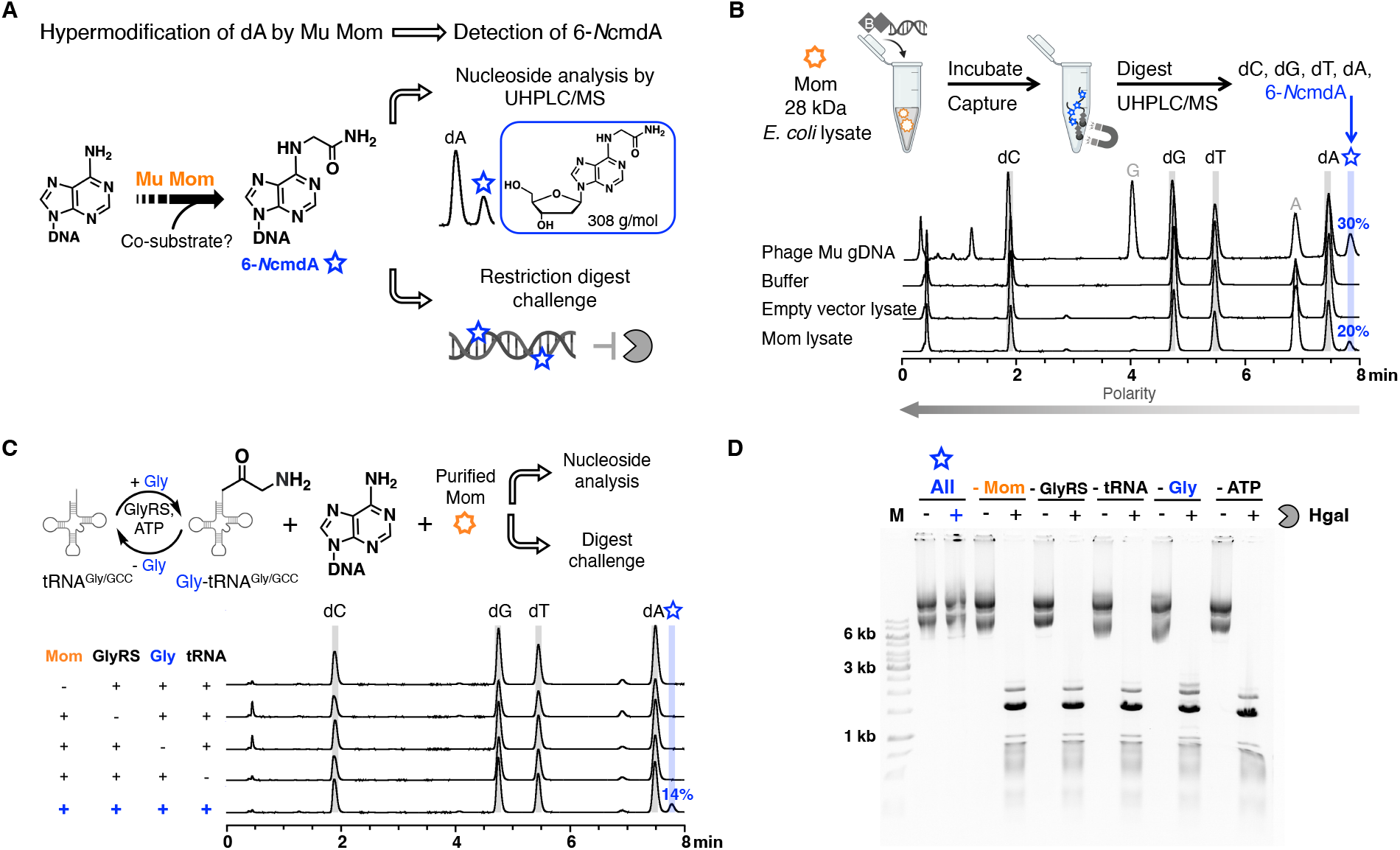
Mom uses Gly-tRNA^Gly^ as a co-substrate to hypermodify the *N*6 position of dA post-replicatively. (**A**) Detecting Mom-catalyzed hypermodification. Following nucleoside digestion of dsDNA hypermodified by Mom, 6-*N*cmdA can be resolved and detected by UHPLC/MS. Alternatively, hypermodified dsDNA can be visualized following challenge with restriction endonucleases. (**B**) Hypermodification of dA by Mom is post-replicative. In an *ex vivo* assay, clarified *E. coli* lysates (20 μg total protein) containing heterologously expressed Mom were incubated with biotinylated dsDNA substrates (250-1000 ng). Nucleoside analysis shows that 6-*N*cmdA is installed by Mom post-replicatively on dsDNA (bottom trace). (**C**) *In vitro* reconstitution of Mom activity demonstrated by nucleoside analysis of hypermodified DNA substrates. 6-*N*cmdA is detected only when purified Mom (2 μM, orange), dsDNA (1000 ng), and Gly-tRNA^Gly/GCC^ (generated *in situ* by GlyRS using tRNA^Gly/GCC^, glycine, and ATP) are included in a coupled activity assay (bottom trace). (**D**) *In vitro* reconstitution of Mom activity demonstrated by restriction challenge of hypermodified DNA substrates. Protection from HgaI digestion was only observed when purified Mom (0.6 μM), substrate plasmid (3.6 μg), and all components to generate charged Gly-tRNA^Gly/GCC^ *in situ* are included in a coupled activity assay (see lanes under All).

## Results

### Adenine hypermodification to 6-*N*cmdA can form post-replicatively on dsDNA

Given the possibility that nucleobases can be modified before or after DNA replication, we first sought to determine if the Mom enzyme accepts the DNA polymer as a substrate. We utilized an *ex vivo* activity assay where clarified lysates derived from *E. coli* cultures heterologously expressing WT Mom are incubated with exogenously added biotinylated DNA substrates that are recovered and analyzed by LC/MS (Fig. 1B and fig. S4) (*21, 22*). After incubation, the 6-*N*cmdA nucleoside was observed, replacing about 20% of the dA as compared to DNA incubated in control lysates. Furthermore, DNA recovered from cells heterologously expressing Mom is specifically modified at dA in the context of an SAS sequence motif as shown by PacBio SMRT sequencing and modification analysis, as well as by challenging DNA products to restriction endonuclease cleavage *in vitro* (fig. S1-S3 and table S1-S2) (*11*). These results demonstrate that Mom acts on double-stranded DNA (dsDNA).

### Mom uses Gly-tRNA^Gly^ as the co-substrate to hypermodify dA

Following the demonstration that Mom acts post-replicatively in lysates, we aimed to reconstitute the momylation reaction *in vitro* and found that Mom was active in the PURExpress® system when expressed through *in vitro* transcription/translation as well as when added exogenously in purified form (fig. S5-S7). The PURExpress® system contains purified enzymes and cofactors sufficient for transcription and translation, including tRNA, amino acids, and tRNA synthetases (*23, 24*). Although enzymes in the GNAT superfamily typically use acyl-CoA substrates to modify their targets, it is well-established that some enzymes of this superfamily use charged tRNA to aminoacylate substrates (*25, 26*). Because glycine and the carbamoylmethyl group of 6-*N*cmdA are structural isomers, we hypothesized that Mom might use Gly-tRNA^Gly^ as a co-substrate to form 6-*N*cmdA.

We tested the activity of purified Mom in a buffered, enzyme coupled assay containing *in vitro* transcribed tRNA^Gly/GCC^ (fig. S8), glycine, ATP, purified *E. coli* glycyl-tRNA synthetase (GlyRS), and linear dsDNA substrate. As can be seen in Fig. 1C and 1D, the modification of DNA by Mom is only detected in those reactions that include tRNA^Gly/GCC^, GlyRS, and ATP. These data unambiguously identify activated glycine as the co-substrate of Mom. *E. coli* tRNA^Gly/CCC^ and tRNA^Gly/TCC^ isoacceptors similarly support momylation (fig. S8), showing that Mom can exploit all tRNA^Gly^ isoacceptors from *E. coli*. Reactions containing pre-charged Gly-tRNA^Gly^, but lacking GlyRS, are sufficient to sustain momylation (fig. S9-S10). We also see that Mom is active on well-matched oligomer duplexes, less active on A:X mismatched oligomer duplexes, and inactive on single-stranded DNA (fig. S11-S12).

### 6-*N*cmdA forms following an on-base isomerization of the exocyclic glycyl group

As highlighted above, the carbamoylmethyl group of 6-*N*cmdA and a glycyl group are structural isomers. To explain the apparent discrepancy between the structures of a modification formed by simple group transfer of the glycyl group to *N*6 versus the mature hypermodification, we hypothesized that, following transfer to the *N*6 position, the glycyl group undergoes isomerization.

We utilized LC/MS-MS and stable isotope labeling to investigate our hypothesis, focusing on a daughter ion of 6-*N*cmdA produced during fragmentation by bond breakage (dashed red line, top reaction of Fig. 2a) at the exocyclic C-C bond (*m/z* 148, termed [6mA^•^]^+^ for clarity) (fig. S13-S14) (*17*). A coupled Mom activity assay including ^15^N,^13^C-glycine (labeled at the primary amine and the carboxy group) and unlabeled dsDNA supported conversion of dA to 6-*N*cmdA (*m/z* = 311) (middle reaction in Fig. 2A and fig. S15). Similarly, reactions containing unlabeled glycine and dsDNA labeled with ^15^N_5_-dA (all 5 nitrogens in dA labeled with ^15^N, see Methods for substrate preparation) supported conversion of dA to 6-^15^*N*cm-^15^N_4_-dA (*m/z* = 314, bottom reaction in Fig. 2A and fig. S16-S17). Fragmentation patterns for 6-*N*cmdA labeled with ^15^N,^13^C-glycine (*m/z* = 311) produced a daughter ion with an *m/z* = 149 that we assigned to [6m-^15^N-A^•^]^+^, one Dalton heavier than the product produced with unlabeled glycine. Given that the labeled nitrogen originated from the primary amine of glycine, the increase in mass here (relative to the unlabeled reaction) can be explained by a rearrangement that must have occurred during the modification reaction. Conversely, reactions utilizing a DNA substrate labeled with ^15^N_5_-dA (*m/z* = 314) produced a daughter ion corresponding to [6m-^15^N_4_-A^•^]^+^ (*m/z* = 152) that is 4 mass units (rather than 5) heavier than [6mA^•^]^+^, as is observed in reactions without labeled components (compare, top and bottom reactions in Fig. 2).

**Figure 2:**
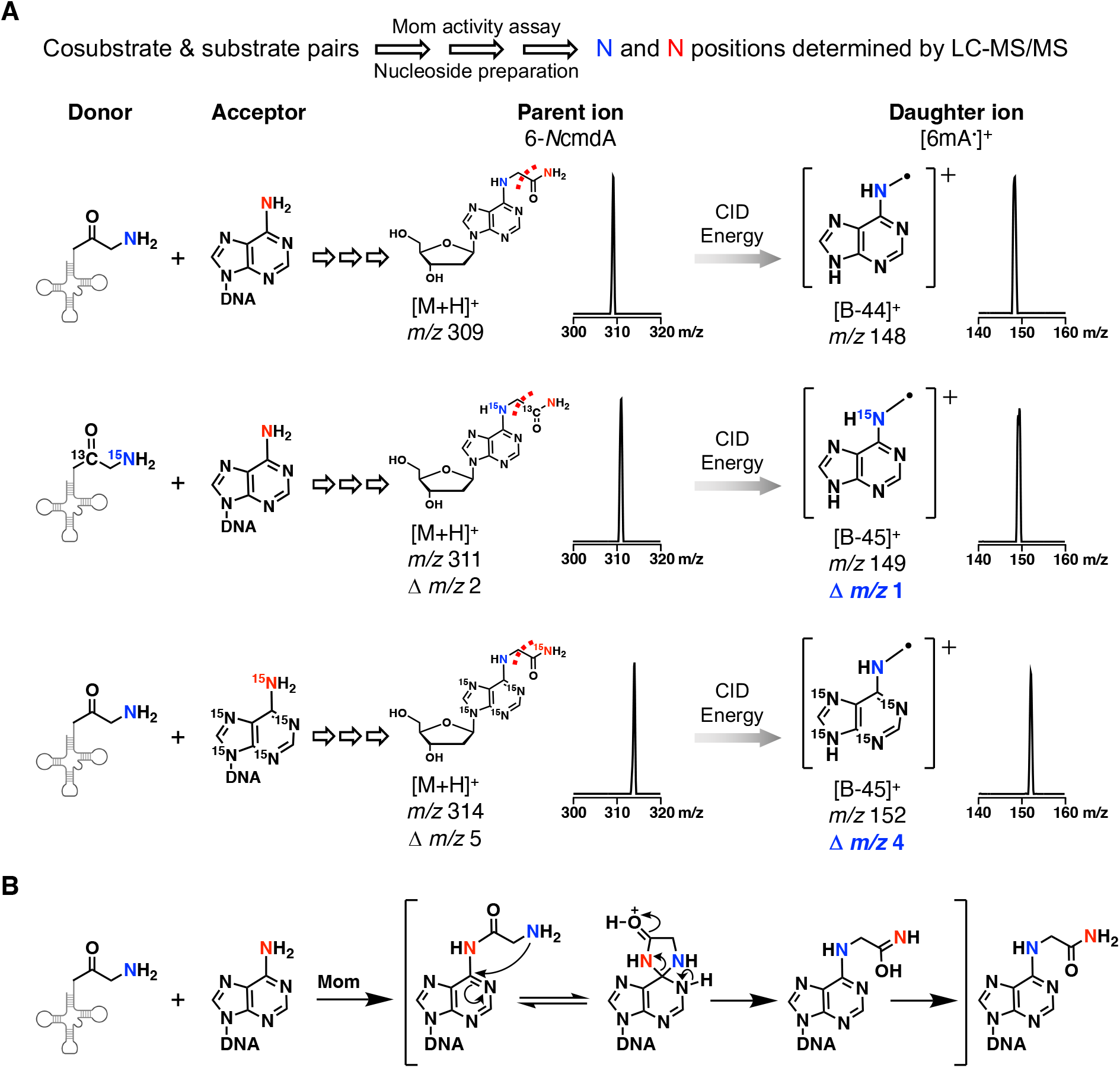
Mom installs 6-*N*cmdA through transfer and rearrangement of the glycyl group. (**A**) On-base rearrangement detected through fragmentation of 6-*N*cmdA and site-specific isotopes. Combinations of unlabeled and isotopically-labeled co-substrate and substrate pairs (left) were used in activity assays to track the positions of the α amine nitrogen from glycine (blue) and the *N*6 nitrogen from dA (red) in the mature hypermodification (center). One daughter ion of 6-*N*cmdA, [6mA^•^]^+^ (*m/z* 148, top), fragments at the exocyclic C-C bond, which separates the two exocyclic nitrogens (top right). In the middle reaction, [6m-^15^N-A^•^]^+^ is 1 Dalton heavier in mass (*m/z* 149), indicating that the ^15^N-labeled α amine nitrogen originating from glycine has rearranged to the *N*6 position. In the complementary experiment (bottom), [6m-^15^N_4_-A^•^]^+^ (*m/z* 152) is 4 Daltons heavier (rather than 5), demonstrating that an on-base rearrangement occurs following transfer of the glycyl group. (**B**) Proposed mechanism for formation of 6-*N*cmdA. Rearrangement of exocyclic groups covalently-bound to dA at *C*6, resulting in a mixture of products, have been reported in abiotic reactions. In a biological context, we propose that following momylation an analogous rearrangement could occur through a five-membered ring intermediate where 6-*N*cmdA is the favored product (transaminoacylation, bracketed steps).

Taken together, these data indicate an exchange between the *N*6 proximal nitrogen and the distal primary amine occurs during momylation (chemically, a transaminoacylation followed by a rearrangement). This “flipping” of the transferred group places the glycyl C_α_ one atom closer to the nucleobase and apparently displaces the carbonyl oxygen one atom further away from dA. To account for the nitrogen exchange, we propose a mechanism for isomerization that proceeds via a five-membered ring intermediate through ring closure upon attack of *C*6 by the exocyclic primary amine followed by ring opening through breakage of the initial *C*6-*N*6 bond to yield the native hypermodification (Fig. 2B). Controlled isomerization of exocyclic groups anchored at the *N*6 position have not previously been reported in a biological system, though rearrangements of exocyclic groups at *N*6 have been induced chemically in abiotic systems. Under harsh conditions, Ross H. Hall and Girish B. Chheda reported carboxymethyl products resulting from tricyclization, hydrolysis, and deamination of adenine aminoacylated at *N*6 (*27, 28*). A more recent study aiming to synthesize peptides conjugated to adenine at *N*6 similarly reported a mixture of products upon deprotection of the primary amine in strong organic acid (*28*). Distinct from these synthetic approaches, enzymatic momylation leads to formation of a single, detectable hypermodification via isomerization of the exocyclic glycyl group, suggesting that the Mom enzyme favors 6-*N*cmdA over any other chemically possible product.

### Crystal structure and mutants reveal putative active site and novel features of Mom

In order to gain a more detailed picture of momylation, we aimed to solve a crystal structure of Mom. Activity assays showed that the truncation of 10 N-terminal residues (Δ10) was well tolerated (fig. S18-S19) (*17*). We also tested variants of the conserved residues R111 and S124, previously identified as potentially important for activity (*17*). In our hands, S124A had much reduced activity, and R111A appeared to be completely inactive (fig. S18-S20). Crystallization screens identified conditions leading to the growth of orthorhombic crystals of Mom(Δ10) S124A in space group P2(1)2(1)2(1) that contained two Mom protomers in the asymmetric unit and that diffracted to ∼2.0 Å resolution (Fig. 3A). The structure was solved by molecular replacement using an AlphaFold generated search model of the Mom protomer, and refined with manual edits in Coot (table S3) (*29–31*).

**Figure 3:**
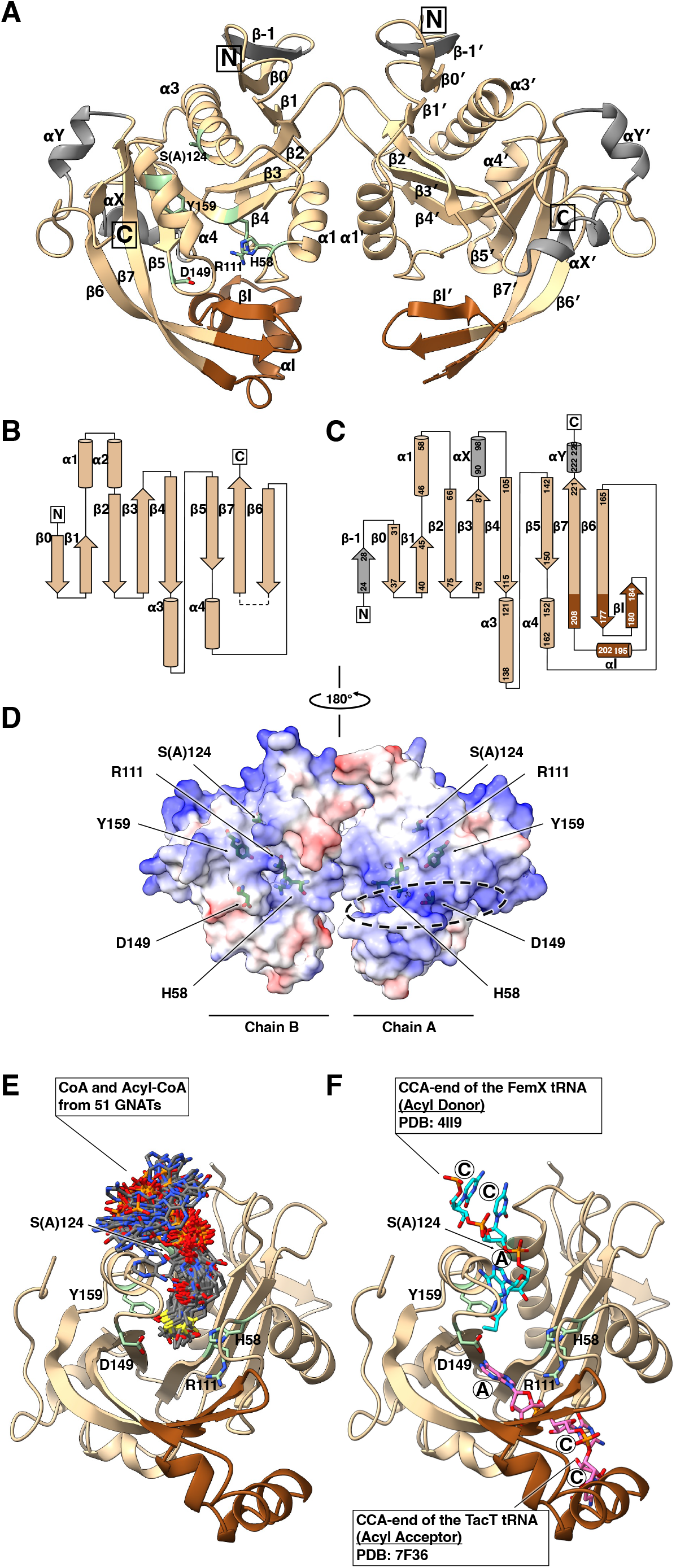
Crystal structure of S124A Mom. (**A**) Ribbon representation of the Mom dimer. Featured are the conserved GNAT core (pale brown), the insertion unique to Mom (brown), residues important for activity (sticks, pale green), and additional regions of low sequence conservation in the GNAT superfamily (gray). (**B**) General GNAT topology diagram. The line that connects β-6 and β-7 is dashed, because β-7 can be contributed by another protomer in some GNATs. (**C**) Mom topology diagram. (**D**) Electrostatic representation of the Mom dimer surface. A positively charged cleft is highlighted by the dashed oval. Residues important for catalysis are labeled. (**E**) Superposition of Mom with 51 diverse GNAT:CoA structures. A Mom protomer (Chain A) was superposed with 51 diverse GNATs co-crystallized with CoA or acyl-CoA (see Methods). Only the overlaid CoA or acyl-CoA molecules of these GNATs are shown. (**F**) Superposition of Mom with GNAT:tRNA structures. Chain A of Mom was superposed with FemX_Wv_ (PBD: 4II9, Ala-tRNA^Ala^ co-substrate) and TacT (PBD: 7F36, Gly-tRNA^Gly^ substrate). Only the 3’-CCA ends of tRNAs are shown (cyan and magenta, respectively). Mom could utilize a FemX-like strategy for binding tRNA and a TacT-like strategy for binding DNA, aided by the positively charged cleft.

Consistent with sizing chromatography results (fig. S6), the structure shows that Mom forms tight dimers with an interface area of ∼850 Å^2^, and a predicted free energy of binding of 4.7 kcal/mol according to the PISA server (*32*). As anticipated (*16, 17, 31*), Mu Mom protomers have the GNAT fold, but with an atypical insertion spanning residues ∼175-209 that visually resembles a helix-turn-helix (HTH) motif (Fig. 3A-3D) (*33–35*). The insertion can also be seen with superpositions and sequence alignments of Mu Mom with the predicted structures of active homologs mined from metagenome databases (fig. S21-S22). Superposition of 51 diverse GNAT crystal structures with Mom suggests residues M107-S124 (including R111 and S124) potentially encompass key active site residues (Fig. 3E and 3F). Superposition of Mom with two GNATS where tRNA is a co-substrate or substrate – FemX_Wv_ (from *Weissella viridescens*, PDB ID: 4II9) and TacT (from *Salmonella typhimurium*, PDB ID: 7F36), respectively –reveals two possible binding modes of the tRNA each that could place the CCA stem within the active site (fig. S23) (*36*).

## Discussion

At multiple points in the evolution of the GNAT superfamily, the fold has adapted to use tRNA as a co-substrate to aminoacylate peptides, proteins, lipids, and natural products (fig. S24) (*37*). As revealed here, the Mom enzyme represents another case where the GNAT fold has adapted to use charged tRNA, but for aminoacylation of a novel substrate: DNA (Fig. 4). Our data demonstrate that Mom co-opts the host translational machinery and uses a readily available source of activated glycine to hypermodify dA. The biochemical data for Mom are consistent with glycyl transfer from the Gly-tRNA^Gly^ ester to the adenine target in an initial first step followed by an intramolecular rearrangement reaction in the second step. Our data leave open whether the substrate base adenine carries out the nucleophilic attack directly at *N*6, or whether there are intermediate transesterification steps. We note that a carbamoylmethyl nucleobase modification (ncm^5^) is also found on the *C*5 carbon of the wobble uridine U_34_ in tRNA of nearly all eukaryotes. It is widely thought that the carbamoylmethyl group represents an amidated version of the carboxymethyl group introduced in a radical reaction catalyzed by the multi-subunit Elongator complex (*38, 39*). If so, then carbamoylmethyl modifications found in tRNA versus DNA differ not only in the anchoring group (*N* versus *C*) but are also introduced by chemically and evolutionarily unrelated mechanisms. As such, the discoveries presented herein highlight the diverse strategies used to install critical hypermodifications onto nucleic acids, provide a foundation for further investigation of the DNA hypermodification reactions made by Mom-like enzymes, and underscore the versatility of the GNAT fold in enzyme evolution.

**Figure 4:**
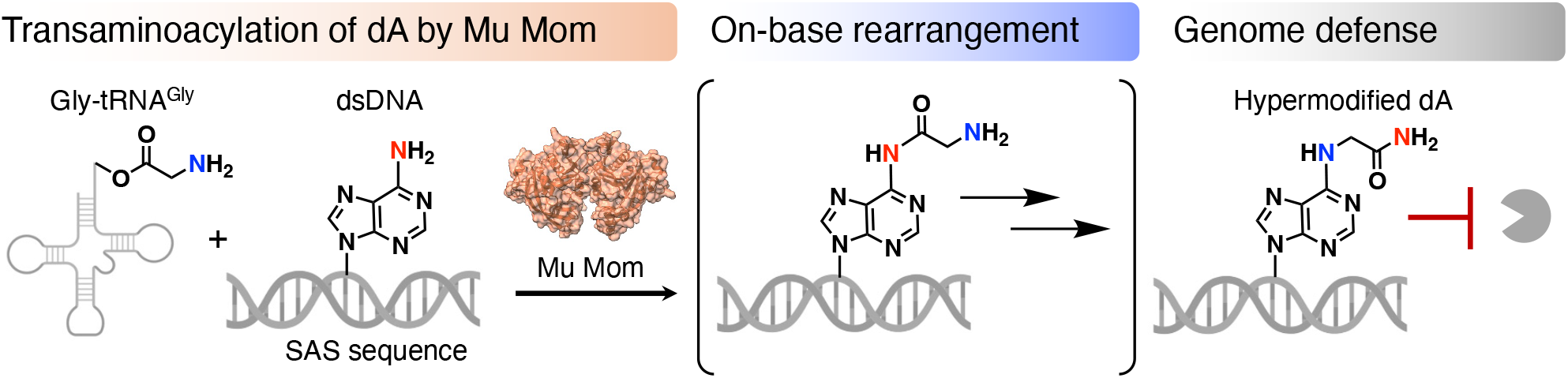
A novel mechanism for genome defense. Our work provides the mechanistic explanation behind a genetically simple but elegant strategy used by the Mom enzyme to protect DNA from endonucleases by hypermodification at *N*6 of adenine through transaminoacylation and isomerization of glycine co-opted from charged tRNA^Gly^.

## Acknowledgments

We thank Johanna Hakanpää and Guillaume Pompidor for access to the PETRA III P11 beamline at DESY, and Dr. Honorata Czapińska, Dr. Elzbieta Nowak and Michal Pastor for data collection. The authors are grateful for purified GlyRS from Corinna Tuckey and Dr. Ying Zhou, T7 RNA polymerase from Professor Janusz Bujnicki’s lab, a plasmid encoding tRNA^Gly/GCC^ from Dr. Haruichi Asahara, and plasmid pRY from Dr. Robert Yarrington. Critical readings of our manuscript were provided by Dr. Ivan R. Corrêa, Dr. Richard J. Roberts, and colleagues at NEB/IIMCB. We are thankful to Dr. Shweta Karambelkar for sharing her unpublished thesis from the Indian Institute of Science, Bangalore. M.B. and A.S. are grateful for funding from the Polish National Science Centre (NCN, 2018/30/Q/NZ2/00669) and the Foundation for Polish Science (FNP, POIR.04.04.00-00-5D81/17-00). R.M.B.S., Y.J.L., C.G., E.A.S., S.R.L., and P.R.W. are grateful for the generous support from the Comb Family and New England Biolabs, Inc. without whom this work would not have been possible. We dedicate this work to the memory of Professor Stanley Hattman.

## Data and materials availability

The final model of the structure solved in this work was deposited in Protein Data Bank under accession code 8BV8. All data are available in the manuscript or the supplementary materials.

## References

1. R. A. J. Warren, Modified Bases in Bacteriophage DNAs. Annu Rev Microbiol. 34, 137–158 (1980).

2. J. H. Gommers-Ampt, P. Borst, Hypermodified bases in DNA. Faseb J. 9, 1034–1042 (1995).

3. P. Weigele, E. A. Raleigh, Biosynthesis and Function of Modified Bases in Bacteria and Their Viruses. Chem Rev. 116, 12655–12687 (2016).

4. M. J. Parker, Y.-J. Lee, P. R. Weigele, L. Saleh, Comprehensive Natural Products III, 465–488 (2020).

5. G. Hutinet, Y.-J. Lee, V. de Crécy-Lagard, P. R. Weigele, EcoSal Plus, in press, doi:10.1128/ecosalplus.esp-0028-2019.

6. R. H. Hall, The Modified Nucleosides in Nucleic Acids (Columbia University Press, 1971).

7. B. Allet, A. I. Bukhari, Analysis of bacteriophage Mu and λ-Mu hybrid DNAs by specific endonucleases. J Mol Biol. 92, 529–540 (1975).

8. A. Toussaint, The DNA modification function of temperate phage Mu-1. Virology. 70, 17–27 (1976).

9. S. Hattman, Unusual Modification of Bacteriophage Mu DNA. J Virol. 32, 468–475 (1979).

10. H. Khatoon, A. I. Bukhari, Bacteriophage Mu-induced modification of DNA is dependent upon a host function. J Bacteriol. 136, 423–428 (1978).

11. S. Hattman, Specificity of the bacteriophage Mu mom+ -controlled DNA modification. J Virol. 34, 277–279 (1980).

12. A. Toussaint, My life with Mu. Bacteriophage. 5, e1034336 (2015).

13. D. Swinton, S. Hattman, P. F. Crain, C.-S. Cheng, D. L. Smith, J. A. McCloskey, Purification and characterization of the unusual deoxynucleoside, a-N-(9-fi-D-2’-deoxyribofuranosylpurin-6-yl)glycinamide, specified. Proceedings of the National Academy of Science (1983).

14. R. Kahmann, A. Seiler, F. G. Wulczyn, E. Pfaff, The mom gene of bacteriophage Mu: a unique regulatory scheme to control a lethal function. Gene. 39, 61–70 (1985).

15. D. H. Krüger, T. A. Bickle, Bacteriophage survival: multiple mechanisms for avoiding the deoxyribonucleic acid restriction systems of their hosts. Microbiol Rev. 47, 345–360 (1983).

16. K. H. Kaminska, J. M. Bujnicki, Bacteriophage Mu Mom protein responsible for DNA modification is a new member of the acyltransferase superfamily. Cell Cycle. 7, 120–121 (2008).

17. S. Karambelkar, S. Udupa, V. N. Gowthami, S. G. Ramachandra, G. Swapna, V. Nagaraja, Emergence of a novel immune-evasion strategy from an ancestral protein fold in bacteriophage Mu. Nucleic Acids Res. 48, 5294–5305 (2020).

18. L. M. Iyer, M. Tahiliani, A. Rao, L. Aravind, Prediction of novel families of enzymes involved in oxidative and other complex modifications of bases in nucleic acids. Cell Cycle. 8, 1698–1710 (2009).

19. L. M. Iyer, D. Zhang, L. Aravind, Adenine methylation in eukaryotes: Apprehending the complex evolutionary history and functional potential of an epigenetic modification. Bioessays. 38, 27–40 (2016).

20. L. M. Iyer, V. Anantharaman, A. Krishnan, A. M. Burroughs, L. Aravind, Jumbo Phages: A Comparative Genomic Overview of Core Functions and Adaptions for Biological Conflicts. Viruses. 13, 63 (2021).

21. Y.-J. Lee, N. Dai, S. E. Walsh, S. Müller, M. E. Fraser, K. M. Kauffman, C. Guan, I. R. Corrêa, P. R. Weigele, Identification and biosynthesis of thymidine hypermodifications in the genomic DNA of widespread bacterial viruses. P Natl Acad Sci Usa. 115, E3116–E3125 (2018).

22. Y.-J. Lee, N. Dai, S. I. Müller, C. Guan, M. J. Parker, M. E. Fraser, S. E. Walsh, J. Sridar, A. Mulholland, K. Nayak, Z. Sun, Y.-C. Lin, D. G. Comb, K. Marks, R. Gonzalez, D. P. Dowling, V. Bandarian, L. Saleh, I. R. Corrêa, P. R. Weigele, Pathways of thymidine hypermodification. Nucleic Acids Res. 50, 3001–3017 (2021).

23. Y. Shimizu, T. Ueda, Cell-Free Protein Production, Methods and Protocols. Methods Mol Biology. 607, 11–21 (2009).

24. C. Tuckey, H. Asahara, Y. Zhou, S. Chong, Curr Protoc Mol Biology, in press, doi:10.1002/0471142727.mb1631s108.

25. L. Favrot, J. S. Blanchard, O. Vergnolle, Bacterial GCN5-Related N-Acetyltransferases: From Resistance to Regulation. Biochemistry-us. 55, 989–1002 (2016).

26. R. M. Burckhardt, J. C. Escalante-Semerena, Small-Molecule Acetylation by GCN5-Related N -Acetyltransferases in Bacteria. Microbiol Mol Biol R. 84 (2020), doi:10.1128/mmbr.00090-19.

27. G. B. Chheda, R. H. Hall, P. M. Tanna, Aminoacyl Nucleosides. V. The Mechanism of the N6-(α-Aminoacyl)adenines into N-(6-purinyl)amino Acids. The Journal of Organic Chemistry (1969).

28. N. Masurier, F. Soualmia, P. Sanchez, V. Lefort, M. Roué, L. T. Maillard, G. Subra, A. Percot, C. E. Amri, Synthesis of Peptide-Adenine Conjugates as a New Tool for Monitoring Protease Activity. Eur J Org Chem. 2019, 176–183 (2018).

29. A. J. McCoy, R. W. Grosse-Kunstleve, P. D. Adams, M. D. Winn, L. C. Storoni, R. J. Read, Phaser crystallographic software. J Appl Crystallogr. 40, 658–674 (2007).

30. P. Emsley, B. Lohkamp, W. G. Scott, K. Cowtan, Features and development of Coot. Acta Crystallogr Sect D. 66, 486–501 (2010).

31. J. Jumper, R. Evans, A. Pritzel, T. Green, M. Figurnov, O. Ronneberger, K. Tunyasuvunakool, R. Bates, A. Žídek, A. Potapenko, A. Bridgland, C. Meyer, S. A. A. Kohl, A. J. Ballard, A. Cowie, B. Romera-Paredes, S. Nikolov, R. Jain, J. Adler, T. Back, S. Petersen, D. Reiman, E. Clancy, M. Zielinski, M. Steinegger, M. Pacholska, T. Berghammer, S. Bodenstein, D. Silver, O. Vinyals, A. W. Senior, K. Kavukcuoglu, P. Kohli, D. Hassabis, Highly accurate protein structure prediction with AlphaFold. Nature. 596, 583–589 (2021).

32. E. Krissinel, K. Henrick, Inference of Macromolecular Assemblies from Crystalline State. J Mol Biol. 372, 774–797 (2007).

33. L. Holm, Dali server: structural unification of protein families. Nucleic Acids Res. 50, W210– W215 (2022).

34. M. W. Vetting, L. P. S. de Carvalho, M. Yu, S. S. Hegde, S. Magnet, S. L. Roderick, J. S. Blanchard, Structure and functions of the GNAT superfamily of acetyltransferases. Arch Biochem Biophys. 433, 212–226 (2005).

35. R. A. Laskowski, J. Jabłońska, L. Pravda, R. S. Vařeková, J. M. Thornton, PDBsum: Structural summaries of PDB entries. Protein Sci Publ Protein Soc. 27, 129–134 (2018).

36. A. M. Cheverton, B. Gollan, M. Przydacz, C. T. Wong, A. Mylona, S. A. Hare, S. Helaine, A Salmonella Toxin Promotes Persister Formation through Acetylation of tRNA. Mol Cell. 63, 86– 96 (2016).

37. M. Moutiez, P. Belin, M. Gondry, Aminoacyl-tRNA-Utilizing Enzymes in Natural Product Biosynthesis. Chem Rev. 117, 5578–5618 (2017).

38. B. Huang, M. J. O. Johansson, A. S. Byström, An early step in wobble uridine tRNA modification requires the Elongator complex. Rna. 11, 424–436 (2005).

39. M. I. Dauden, J. Kosinski, O. Kolaj-Robin, A. Desfosses, A. Ori, C. Faux, N. A. Hoffmann, O. F. Onuma, K. D. Breunig, M. Beck, C. Sachse, B. Séraphin, S. Glatt, C. W. Müller, Architecture of the yeast Elongator complex. Embo Rep. 18, 264–279 (2017).

