## Supplementary Materials for "Modification of DNA by a viral enzyme and charged tRNA"

**Affiliations:**

**This PDF file includes:**

Materials and Methods
Figures S1 to S24
Tables S1 to S3

Fig. S15. Fragmentation of 6-*N*cmdA labeled with $^{15}N,^{13}C$-glycine ($H_3{}^{15}NCH_2{}^{13}CO_2H$) demonstrates that the glycyl group flips 180° during momylation, exchanging the nitrogen from the $\alpha$-amine originating from glycine and the nitrogen originating from the *N*6 position of dA. 34

### Materials and Methods

Reagents, media, enzymes, and general procedures.

All reagents were used as received and stored according to the manufacturer's instructions and used without additional purification unless otherwise specified. Stock solutions were made with autoclaved water purified on a Milli-Q Reference Ultrapure Water Purification System (≥18.2 MΩ cm) (MilliporeSigma, Burlington, MA, USA) or with Nuclease-Free Water from Invitrogen (Waltham, MA, USA) where specified. Media and media components were obtained from Beckon Dickinson-Difco (Franklin Lakes, NJ, USA) and Sigma-Aldrich (St. Louis, MO, USA). All commercially available enzymes used were manufactured by New England Biolab® (NEB, Ipswich, MA, USA) unless otherwise specified. Plasmids, primers, co-substrates, and substrates are listed in Table S2. Chemical structures were made using ChemDraw®. Several figures were created with BioRender.com.

Phage Mu preparation and genomic DNA isolation.

Two *E. coli* strains carrying phage Mu lysogen were obtained from Stan Hattman and were used to produce Mu phages. Of the two strains, one contains the Mu lysogen (denoted as ER3153; genotype: *sup mcrA thi hsdR mcrB zzz::Mu(Cts62)*) which produces the Mu phage with the native 6-*N*cmdA DNA modification phenotype, and the other has a mutant Mu lysogen which produces Mu phages without 6-*N*cmdA DNA modification but canonical 2′-deoxyadenosine phenotype (denoted as ER3154; genotype: *sup mcrA thi hsdR mcrB zzz::Mu(Cts62 mom)*).

A heat-shock procedure was performed to induce the production of Mu phage from the lysogen. LB medium was inoculated with the phage lysogen strain from an overnight culture and was incubated at 37 °C with agitation until an $OD_{600}$ reading of ~1. The culture was then incubated at 45 °C for 30 minutes and then again at 37 °C with agitation. Phage were harvested when the culture appeared translucent (cell lysed), a process that normally takes about 1.5 – 3 h after heat-shock. The phage lysate solution was then centrifuged at 10000 rcf x g at 4 °C for 10 min, and the supernatant was collected. The phage was then concentrated first by precipitation in a solution of 10% (w/v) poly(ethylene glycol) average MW 8 kDa and 1 M NaCl, followed by pelleting by centrifugation, and finally resuspension in a buffer composed of 50 mM Tris-HCl 7.5, 10 mM $MgCl_2$ and 75 mM NaCl. The phage solution was stored at 4 °C in the dark.

A previously published protocol was followed to extract and purify phage genomic DNA (*1*). Briefly, the phage solution from previous step was diluted in a solution of 100 mM Tris pH 8, 25 mM ethylenediaminetetraacetic acid (EDTA), 1 % (v/v) sodium dodecyl sulfate (SDS), and 200 µg/mL proteinase K and then incubated at 55 °C for 30 minutes until clear. Lysed bacteriophage solution was extracted with an equal volume of a mixture containing phenol, chloroform, and isoamyl alcohol at a ratio of 25:24:1 at neutral pH and agitated by hand for 3 minutes. Centrifugation was then used to separate phases, and the aqueous phase was re-extracted two times with equal volumes of chloroform. DNA was precipitated by addition of 1/10 volume of 3 M sodium acetate and 2.25 volumes of ice-cold ethanol and incubated on ice for 3 h to overnight. DNA was gently spooled onto a glass rod and successively immersed three times in 70 % ethanol to remove salts. DNA was dissolved in 10 mM Tris-HCl pH 8 buffer and stored at 4 °C.

SMRT Sequencing and SMRT Sequencing Data Analysis

Genomic DNA isolated from phage Mu was used to prepare the single-molecular real-time (SMRT) sequencing libraries. DNA was first sheared to around 2 kb fragments in TE buffer using Covaris Adaptive Focused Acoustics ultrasonication system ME220 in microTUBE-130 AFA Fiber sample tubes (Covaris, Woburn, MA). The sheared DNA was end-repaired and ligated to PacBio hairpin adapters to prepare the SMRTbell template libraries following the protocol "Procedure & Checklist – Preparing Multiplexed Microbial Libraries Using SMRTbell Express Template Prep Kit 2.0" published by Pacific Biosciences (Pacific Biosciences, Menlo Park, CA). Linear DNA fragments and library fragments containing a single adapter were digested with exonuclease III and exonuclease VIII. DNA qualification and quantification were performed using the Qubit fluorimeter (Thermo Fisher Scientific, Waltham, MA) and 4200 TapeStation System (Agilent Technology, Santa Clara, CA). SMRT sequencing was carried out on the PacBio Sequel system using standard protocols for small insert SMRTbell libraries with Sequel 3.0 chemistry.

SMRT sequencing reads were analyzed using SMRT Analysis software and pipelines from SMRT Link webtool (version 10.2.0) hosted by Pacific Biosciences. SMRT sequencing raw CCS reads were processed to generate HiFi reads in which the output sequences were used for the *de novo* assembly of the genome using MEGAHIT assembler software (version 1.2.9). (*2*). The longest assembled output sequence was used as the reference for subsequent SMRT Analyses steps. Base modification detection and motif analysis were performed with the same SMRT Link version. To detect the base modifications, the Base Modification Analysis application was used to first map the raw reads to the assembled genome and the mapped reads were analyzed further to extract the inter-pulse duration (IPD) kinetics information collected during sequencing course and processed for all pulses aligned to each position in the reference sequence using an in silico kinetic reference and a t-test based kinetic score detection of modified base positions. The per-base IPD ratio mapped outputs were further used to extract the modification motifs. The top three identified modification motif sequences were extracted separately and the sequences for each motif string were used to generate sequence logo by using WebLogo 3 webtool (version 3.7).

Restriction digest protection assay.

To assay for installation of 6-*N*cmdA by Mom, Mu gDNA or DNA substrates of Mom (see below) were subject to restriction digests with 6-*N*cmdA-sensitive restriction enzymes HgaI (NEB) or AasI FastDigest (Thermo Scientific™, Waltham, MA, USA) according to the manufacturer's instructions (*3*). The digestion products were resolved on an 0.8% agarose gel and imaged using GelRed (Biotium, Fremont, CA, USA) staining.

UHPLC-MS and UHPLC-MS/MS analysis of nucleosides.

Nucleoside samples of Mu gDNA or DNA substrates of Mom were prepared by using the Nucleoside Digest Mix Kit (NEB) in 30 µL reaction volumes according to the manufacturer's instructions (*1*, *4*, *5*). Generally, 0.5-1 µL of digest mix was used per 1 µg of DNA sample. Reactions were incubated for at least 1 hour and up to overnight at 37 °C. Nucleosides were filtered through Ultrafree-MC Centrifugal Filters (MilliporeSigma, 0.22 µm pore size, hydrophilic PVDF). Nucleosides were subjected to ultra-high-performance liquid chromatography (UHPLC) or UHPLC-mass spectrometry (UHPLC-MS) on an Agilent (Santa Clara, CA, USA) 1290 Infinity II UHPLC system equipped with a G7117 Diode Array Detector. Nucleosides were resolved on a Waters (Milford, MA, USA) XSelect HSS T3 C18 column (2.1 × 100 mm, 2.5 µm particle size) and operated at a flow rate of 0.6 mL/min with a linear gradient of aqueous buffer (10 mM ammonium acetate, pH 4.5) and methanol over 6 minutes. The course of chromatography was monitored at 260 nm. LC-MS was carried out on an Agilent 1290 Infinity II UHPLC-MS system equipped with a G7117 Diode Array Detector and a LC/MSD XT G6135 Single Quadrupole Mass Detector operated in both positive (+ESI) and negative (-ESI) electrospray ionization modes. MS was performed with a capillary voltage of 2500 V at both modes, a fragmentor voltage of 70 V, and a mass range of m/z 100 to 1000. The pre-MS liquid chromatography was performed with the same hardware and parameters as aforementioned LC method. Agilent ChemStation software was used for LC and LC-MS data processing. The peak area of each nucleoside species resolved in the UHPLC trace (recorded as absorbance at 260 nm) was measured using the integration function of the Agilent ChemStation software. Traces and integration values were exported as PDF files and further formatted and analyzed in Adobe Illustrator and Microsoft Excel respectively. The peak areas were divided by the corresponding nucleoside molar extinction coefficient (ε) at 260 nm. The extinction coefficients used were 7100 $cm^{-1}$ $M^{-1}$ for dC, 12180 $cm^{-1}$ $M^{-1}$ for dG, 8560 $cm^{-1}$ $M^{-1}$ for dT, and 15060 $cm^{-1}$ $M^{-1}$ for dA (*6–8*). The percent replacement of adenine with 6-*N*cmdA was calculated from the (ε)-adjusted quantity of nucleoside using the equation % 6-*N*cmdA formation (6-*N*cmdA/[dA+6-*N*cmdA]). The equation for % 6-*N*cmdA formation assumed that the extinction coefficients for dA and 6-NcmdA are equivalent.

Fragmentation of nucleosides by LC-MS/MS was performed on an Agilent 1290 Infinity II UHPLC system equipped with a G4212A diode array detector and a 6490A triple quadrupole mass detector operating at the positive electrospray ionization mode (+ESI). UHPLC was performed on a Waters XSelect HSS T3 C18 column (2.1 × 100 mm, 2.5 µm particle size) at a flow rate of 0.6 mL/min with a linear gradient of aqueous buffer (10 mM aqueous ammonium formate, pH 4.4) and methanol over 6 minutes. MS/MS fragmentation spectra were obtained by collision-induced dissociation in the positive product ion mode with the following parameters: gas temperature 200 °C, gas flow 14 L/min, nebulizer 45 psi, sheath gas temperature 350 °C, sheath gas flow 11 L/min, capillary voltage 2 kV, nozzle voltage 1.5 kV, fragmentor voltage 380 V and collision energy 5–65 V.

Constructs, cloning, and strains

Cloning was performed in *Escherichia coli* NEB® 5-alpha (NEB) or TOP10 strains (Invitrogen). Protein expression was performed in *E. coli* T7 Express (NEB) or BL21-CodonPlus (DE3)-RIL strains (Agilent). Plasmids were stored at -20 °C until use.

*Mom*. Primer pairs YJLo260/YJLo261 or YJLo260/YJLo263 were used to PCR amplify the *mom* genes using Q5 DNA polymerase (NEB) with the genomic DNAs of purified phage Mu from strains ER3153 or ER3154 as the PCR templates. NEBuilder HiFi DNA Assembly reagents (NEB) were used to assemble the PCR products of *mom* genes and the linearized pRY001 vector to generate the expression construct with the *mom* gene under the control of *T7lac* promoter. The pRY001 vector is derived from pET28c vector with a 387 bp insertion between *lacO* operator and RBS. The construct made from the PCR product of primer pair YJLo260/YJLo263 has the *Mom* gene containing open reading frame with no additional sequence; while the construct made from primer pair YJLo260/YJLo263 has the *Mom* gene containing open reading frame with additional 3′-end sequences that encodes 6xHis tag C-terminal of the expressed Mom protein. Expression cells harboring the pRY Mu Mom plasmid were used in downstream *in vivo* and *ex vivo* Mom activity assays. See below.

The synthetic open reading frame of Phage Mu full-length Mom protein (NP_050657.1; 241 amino acids), codon optimized for the expression in *E. coli*, was purchased from GeneArt (Thermo Scientific$^{TM}$). It was cloned into the pET28a vector under the T7 promoter by the restriction sites of NdeI and XhoI, adding an N-terminal 6xHis-Tag and thrombin cleavage site. Truncations of Mom N-terminal amino acids (Δ10 and Δ20), the deletion of Mom-specific insertion (Δ174-211, ΔI) and the site-directed mutagenesis (R111A and S124A) were performed by the quick-change method (PCR with subsequent degradation of the template plasmid with DpnI). In order to make a 6xHis-SUMO-Δ10 Mom version, Mom Δ10 ORF was ordered from GeneArt and cloned into the pET28a-6xHis-SUMO vector downstream of the SUMO tag by the restriction sites BamHI and XhoI. R111A and S124A variants of 6xHis-SUMO-Δ10 Mom were also created by the quick-change protocol. Expression cells harboring pET28a plasmids expression WT Mom or Mom variants were used for *in vivo* Mom activity assays or for purifying enzyme for crystallography.

*Glycyl-tRNA synthetase (GlyRS)*. The *E. coli* GlyRS expression construct, pET21a-GlyRS-$His_6$, was a gift from Sebastian Maerkl & Takuya Ueda (Addgene plasmid #124110; http://n2t.net/addgene:124110; RRID:Addgene_124110).

*E. coli tRNA$^{Gly}$ genes*.
The pUC19 plasmid containing the gene encoding the tRNA$^{Gly/GCC}$ isoacceptor from *E.coli* was kindly gifted by Haruichi Asahara. The genes encoding for the tRNA$^{Gly/CCC}$ and tRNA$^{Gly/TCC}$ isoacceptors were ordered from GenScript (Piscataway, NJ, USA) on a pUC19 plasmid. Sequences of the tRNA$^{Gly}$ isoacceptors from *E. coli* were accessed at the GtRNAdb Database (*9*, *10*).

Recombinant protein expression.
Mom was heterologously expressed in T7 Express or BL21-CodonPlus (DE3)-RIL strains. A single colony from freshly transformed T7 Express cells was used to inoculate LB/Kan (40 µg/mL) media (+ glucose, 0.1%). The starter culture was grown overnight at 37 °C at 240 rpm. A 1:100 to 1:1000 dilution of the overnight culture was made into fresh expression LB media (- glucose). Cells were grown to an $OD_{600}$ of 0.6 at 240 rpm and expression was induced with 100 µM IPTG. Cells were grown for 18 hours at 18 °C after which they were harvested by centrifugation and stored at -20 or -80 °C until use. Mom was also heterologously expressed in the BL21-CodonPlus (DE3)-RIL strain as was GlyRS in LB media under the selective pressure of kanamycin-chloramphenicol and ampicillin-chloramphenicol, respectively. Cells were scraped and used to inoculate starter cultures that were grown in the presence of 1% glucose up to a late log phase at 37 °C in a rotary shaker at 140 rpm. A 1:200 dilution of starter culture was used to inoculate expression media. After the $OD_{600}$ reached 0.6-0.7, the expression was induced with 250 µM IPTG. Cells were incubated with 140 rpm shaking for 18-20 h at 25 °C. Cells were then collected by centrifugation, washed with PBS (pH 7.4) and stored at -20 °C.

*In vivo* Mom activity assays.
To test the *in vivo* activity of Mom and its variants, expression plasmids from cells harvested after recombinant expression (as described in the "Protein Expression" section) were isolated by miniprep using the Monarch® Plasmid DNA Miniprep Kit (NEB) or by midiprep using the Midi-

Prep Kit (Syngen Biotech, Wrocław, Poland) kits. Plasmids were frozen until use. As a negative control, expression strains bearing the pET28a vector with no insert ("Empty") were used. Purified plasmids were challenged with the Restriction Digest Protection Assay and nucleoside digests of plasmids were analyzed by UHPLC-MS as described above.

<u>Preparation of biotinylated DNA substrates.</u>
Biotinylated DNA substrates were prepared digesting λ DNA (NEB) with MseI (NEB) in CutSmart Buffer (NEB) at 37 °C overnight (*1*, *4*, *5*). (25 units of MseI per 5 µg of DNA). The QIAquick Nucleotide Removal Kit (Qiagen, Hilden, Germany) was used to clean up digest reactions; 30 µL of elution buffer was used for each column. Digested λ DNA was backfilled with 50 µM Biotin-16-(5-aminoallyl)-dUTP (Jena Bioscience, Jena, Germany) using Klenow Fragment (3′→5′ exo-) (NEB) at a final concentration of 1 U/µL in NEBuffer™2 at 37 °C overnight. Ligation reactions were cleaned up with QIAquick Nucleotide Removal Kit and recovered DNA substrates were quantified by NanoDrop™ (Thermo Scientific™). Biotinylated DNA was stored at -20 °C until use.

<u>Preparation of clarified lysate.</u>
Clarified lysate was prepared by resuspending harvested T7 Express cells reserved from 10 mL of expression culture in prechilled lysis buffer (10 mM Tris·HCl, pH 8, 100 mM NaCl, 10 mM KCl, filtered) with components from VWR International (Radnor, PA, USA) and Sigma-Aldrich (*1*, *4*, *5*). In a chilled cuvette holder, cells were lysed using the Q500 Microtip 4-Prong Sonicator (Qsonica, Newtown, CT, USA) for 2 min with a 25% duty cycle at 30% power. Lysed cells were clarified by centrifugation at 21,000 × g at 4 °C for 15 min. Total protein concentration in the supernatant was quantified by the Bradford assay (Bio-Rad, Hercules, CA, USA) with BSA as a standard. Aliquots of clarified lysate were flash frozen and stored at −80 °C until use.

<u>*Ex vivo* Mom activity assays.</u>
All the following steps on completed ice. In a 15-40 µL reaction, *ex vivo* reaction buffer (25 mM Tris·HCl, pH 7.5, 5 mM $MgCl_2$, and 25 mM KCl, filtered and supplemented with 5 mM fresh DTT (NEB) the day of experiment, 1 µL of 20 mg/mL RNase A, 10-25 µg total protein from clarified lysate, and 250-1000 ng biotinylated DNA substrate was mixed and incubated at 37 °C for > 15 min (*1*, *4*, *5*). An equal volume of quench buffer (66 mM Tris, pH 7.5, 10 mM $MgCl_2$, 1 mM DTT, 7.5% PEG 6000) was added at room temperature. PEG 6000 was purchased from Sigma Aldrich. Some reactions were supplemented with $^{13}C_2$-acetyl-CoA (Sigma Aldrich) that had been stored in 1 M aliquots at -80 °C in water until use.

<u>Recovery and digestion of biotinylated DNA substrates.</u>
Streptavidin Magnetic Beads (NEB) were used to recover biotinylated DNA substrates from enzymatic reactions (*1*, *4*, *5*). The beads were prepared by washing in 5 volumes of Wash Buffer (20 mM Tris·HCl, pH 7.5, 0.5 M NaCl, and 1 mM EDTA) three times (buffer components from VWR, NEB, Invitrogen). Each quenched *ex vivo* reaction was incubated with 30 µL prepared slurry for > 10 min at ambient temperature. The beads were captured by a magnet, the supernatant was removed, and beads were washed 5 volumes of Wash Buffer three times. After the last wash, beads were incubated in a nucleoside digest mix reaction (30 µL) at 37 °C for 1 hr and up to overnight. Beads were removed with a magnet, filtered, and analyzed by UHPLC-MS as described above.

Protein purification.
All protein purification steps were carried out at the temperature range 0 – 12 °C. All buffers were filtered through 0.45 µm filter and degassed by vacuum pump. Equipment used for protein purification was ÄKTA Purifier FPLC system (GE Healthcare) and Minipuls 3 peristaltic pump (Gilson).

*Mom (N-ter 6xHis-Tag full-length MOM wt and R111A)*. Pellets from Mom-expressing cells were suspended in the Buffer 1 (50mM Sodium Phosphate, pH 8.0, 750 mM NaCl, 20% glycerol, 2 mM imidazole, 10 mM 2-mercaptoethanol (2-ME)) supplemented with 1mM PMSF and sonicated. The lysates were clarified by centrifugation at 15 000 g for 40 min. The soluble fraction was applied on 5mL HisTrap column (Cytiva) equilibrated with Buffer 1. The column was then washed with 100 mL of each Buffer 2 (50 mM Sodium Phosphate, pH 8.0, 2M NaCl, 20% glycerol, 2 mM imidazole, 10 mM 2-ME), Buffer 3 (50mM Sodium Phosphate, pH 8.0, 750 mM NaCl, 20% glycerol, 20 mM imidazole, 10 mM 2-ME), and Buffer 4 (20 mM Tris-HCl, pH 7.5, 700 mM NaCl, 10% glycerol, 10 mM 2-ME). The protein was eluted with 0 – 500 mM linear imidazole gradient in Buffer 4. Mom containing fractions were mixed with Buffer 5 (20 mM Tris-HCl, pH 7.5, 5 mM NaCl, 10% glycerol, 10 mM 2-ME) in 1:6 ratio and applied on 5 mL HiTrap Heparin column (Cytiva, Marlborough, MA) equilibrated with buffer 6 (20 mM Tris-HCl, pH 7.5, 100 mM NaCl, 10% glycerol, 10 mM 2-ME). Mom was eluted with 100 – 1000 mM NaCl gradient in Buffer 6. Mom containing fractions were buffer exchanged and concentrated to 1 mL using Amicon 10 MWCO centrifugal filter unit (Merck) and applied on Superdex 200 Increase 10/300 GL column (Cytiva) equilibrated with Buffer 7 (20 mM Tris-HCl, pH 7.5, 600 mM NaCl, 10% glycerol, 10 mM 2-ME). The best quality Mom fractions were either flash-frozen in liquid nitrogen and stored at -80 °C or mixed 1:1 with glycerol and stored at -20 °C.

*Mom (6xHis-SUMO-Δ10 Mom wt and S124A)*. Mom expressing bacterial pellets were suspended in Buffer 1 supplemented with 1mM PMSF and sonicated. The lysates were clarified by centrifugation at 15000g for 40 min. The soluble fraction was applied on 5 mL HisTrap column (Cytiva) equilibrated with Buffer 1. The column was then washed with 100 mL of each Buffer 2, Buffer 3, and Buffer 4H (20 mM HEPES-KOH, pH 7.5, 300 mM NaCl, 10% glycerol, 10 mM 2-ME). The protein was eluted with 0 – 500 mM linear imidazole gradient in Buffer 4H. Mom containing fractions were supplemented with SUMO protease (home-made) at final concentration of ~ 5 µg/mL and incubated at 8 °C for 5 hours. The protein solution was then mixed with Buffer 5H (20 mM HEPES-KOH, pH 7.5, 5 mM NaCl, 10% glycerol, 10 mM 2-ME) in 1:2 ratio and applied on 5 mL HiTrap Heparin column (Cytiva) equilibrated with buffer 6H (20 mM HEPES-KOH, pH 7.5, 100 mM NaCl, 10% glycerol, 10 mM 2-ME). Mom was eluted with 100 – 1000 mM NaCl gradient in Buffer 6H. Mom containing fractions were buffer exchanged and concentrated to 1 mL using Amicon 10 MWCO centrifugal filter unit (Merck) and applied on Superdex 200 Increase 10/300 GL column (Cytiva) equilibrated with Buffer 7H (20 mM HEPES-KOH, pH 7.5, 600 mM NaCl, 10% glycerol, 10 mM 2-ME). The best quality Mom fractions were either mixed 1:1 with glycerol and stored at -20°C or short-term stored on ice before proceeding to the crystallization.

*GlyRS*. Aliquots of *E.coli* GlyRS were kindly gifted by Corinna Tuckey and Dr. Ying Zhou. *E.coli* GlyRS was also purified according to the protocol from Yoshihiro Shimizu & Takuya Ueda with additional steps (*11*). Namely, after the elution from Ni-NTA column the stoichiometric ratio of GlyQ and GlyS appeared shifted towards GlyS. Therefore, the hetero-tetrameric GlyRS was enriched by two consecutive runs on Superdex 200 Increase 10/300 GL column equilibrated with Buffer 8 (2x PBS, pH 7.4, 10 mM 2-ME). Purified protein was mixed 1:1 with glycerol and stored at -20 °C.

Analytic size-exclusion chromatography.
The oligomeric state of Mom protein was tested using a Superdex 200 Increase 10/300 GL column and ÄKTA Purifier FPLC system. The sample containing 1 mg of N-ter-6His-Mom WT in 1 mL of Buffer 7 was applied on the column pre-equilibrated with the same buffer and resolved at 0.4 mL/min flow rate. Gel Filtration Standard (Bio-Rad) was used to generate a standard curve.

PURExpress® activity assays with Mom.
For cell free protein synthesis (also referred as *in vitro* transcription and translation or IVTT), the PURExpress® system (NEB) was used with 1000 ng of pRY WT Mom plasmid as the template in a reaction incubated at 25 °C for 24 hours according to the manufacturer's instructions (*11*, *12*). A control synthesis reaction was assembled as a control according to the manufacturer's instructions. WT Mom plasmid was recovered using the Monarch® PCR and DNA Cleanup Kit (NEB) and nucleoside samples were prepared and analyzed by UHPLC-MS as described above. The PURExpress® system was also utilized for activity assays (without plasmid template), which included PURExpress® kit components, purified WT Mom (2 µM), and 1000 ng of biotinylated DNA substrates. Reactions were incubated at 25 °C for 24 hours or 37 °C for ≥ 2 hours and biotinylated substrates were recovered, digested, and analyzed by UHPLC-MS as described above.

*In vitro* transcription of $tRNA^{Gly}$ genes.
*E. coli* $tRNA^{Gly/GCC}$, $tRNA^{Gly/CCC}$, $tRNA^{Gly/TCC}$ genes with an upstream T7 promoter were amplified from pUC19 plasmids (1 ng) with RS003/RS002, RS003/RS049, and RS003/RS047 primers, respectively, using the Q5® High-Fidelity 2X Master Mix (NEB) (*9*, *10*, *13*). Temperature cycling was as follows: 1. 30 seconds at 98 °C; 2. 25 cycles of 10 seconds at 98 °C, 30 seconds at 63 °C, and 30 seconds at 72 °C; 3. 120 seconds at 72 °C. Amplification was confirmed by gel electrophoresis using 1.2% Latitude™ Precast Agarose Gels (Lonza, Basel, Switzerland) run at 190 V for 45-90 minutes and stained with ethidium bromide. The ~250 bp amplicons were purified using the Monarch® PCR and DNA Cleanup Kit. $tRNA^{Gly}$ genes were transcribed using HiScribe™ T7 High Yield RNA Synthesis Kit (NEB) using 1000 ng of PCR template and GMP (20 mM, Sigma-Aldrich, St. Louis, MO, USA) with a 16-hour incubation at 37 °C. tRNA was purified using the Monarch® RNA Cleanup Kit (NEB), quantified using NanoDrop, and stored at -80 °C until further use. $tRNA^{Gly/GCC}$ was also transcribed using purified T7 polymerase (kindly gifted by Prof. Janusz Bujnicki's lab) in previously described conditions (*14*) and purified by size exclusion chromatography using a Superdex 200 Increase 10/300 GL column equilibrated with RNA Buffer (20 mM HEPES pH 7.5, 50 mM KCl, 50 mM NaCl, 1 mM $MgCl_2$, 1 mM DTT) and stored at -80 °C. Transcription products were mixed 1:1 in RNA Loading Dye (NEB), heated for 3-5 min at 95 °C, loaded (200 ng/lane) onto 6% or 15% Novex™ TBE-Urea Gels (Invitrogen™), and electrophoresed at 200 V in 1X TBE buffer for 30 minutes or 80 minutes, respectively, at ambient temperature with Low Range ssRNA Ladder (NEB) used as the standard. Gels were stained with SYBR™ Gold Nucleic Acid Gel Stain (Invitrogen™) in 1x TBE buffer for > 20

minutes and imaged on the Typhoon™ (Amersham) using the Cy2 channel, 400 PMT, and 50-100 µm pixel size. Images were analyzed using ImageLab (Bio-Rad).

<u>Oligo DNA substrates.</u>

Oligomers were ordered from Integrated DNA Technologies (IDT; Coralville, Iowa, USA) or Genomed (Warsaw, Poland) (Table S2). The sequence of the $WM_{Mu}$ 34-mer was taken from the phage Mu genome. Double-stranded DNA was obtained by heating a 1:1 mix of complementary oligonucleotides at 90 °C for 5 min and annealing by gradual cooling to the room temperature for 30 min. Duplexes were stored at -20 °C until use. $WM_{Mu}$ 34-mer was resuspended in Duplex Buffer (IDT), complementary strands were mixed in equimoloar ratios, and annealed by heating at 95 °C for 2 minutes and cooling to room temperature over a linear gradient. Annealing was confirmed using a 20% Novex™ TBE Gel (Invitrogen™) run at 180 V in 1X TBE buffer at 4 °C for 2 hours.

<u>3 kbp DNA substrates.</u>

The 3 kbp substrates were amplified by PCR from the pUC19 $tRNA^{Gly/GCC}$ plasmid (20 ng) with RS005/RS006 primers using the Taq Polymerase in Thermopol Buffer (NEB). The 3 kbp substrate was amplified using the supplied dNTP mix, and the $^{15}N_5$-3kbp substrate was amplified in the presence of $^{15}N_5$-dATP (Cambridge Isotope Laboratories), dGTP, dTTP, and dCTP. Temperature cycling is as follows: 1. 30 seconds at 98 °C; 2. 25 cycles of 10 seconds at 98 °C, 30 seconds at 62 °C, and 90 seconds at 72 °C; 3. 120 seconds at 72 °C. Amplification was confirmed by gel electrophoresis using 1.2% Latitude™ Precast Agarose Gels (Lonza, Basel, Switzerland) run at 190 V for 45-90 minutes and stained with ethidium bromide. The 3 kbp amplicons were purified using the Monarch® PCR and DNA Cleanup Kit and quantified by NanoDrop.

<u>Coupled Mom activity assay with biotinylated dsDNA or $WM_{Mu}$.</u>

Components for the coupled Mom activity assay were mixed on ice in the following order to a final volume of 30 µL and incubated 5 hours – overnight at 37 °C: activity buffer (50 mM HEPES pH 7.6, 100 mM KCl, 13 mM magnesium acetate, 1 mM DTT, and 2 mM ATP); Pyrophosphatase, Inorganic (0.025 units/mL, NEB); GlyRS (2-4 µM); glycine (2 mM); RNase Inhibitor, Murine (20 units, NEB); Mom (2 µM); DNA substrate (250 ng – 10 µg); $tRNA^{Gly}$ (~3 µg [4 µM]); and RNase-free water (*13*, *15*). Where specified, $^{13}C,^{15}N$-glycine (2 mM, Sigma) was used in place of glycine. For 34-mer dsDNA substrate, 1000 ng of duplex corresponds to concentration of 1.5 µM, and 10 µg of duplex corresponds to a concentration of 15 µM. Reaction components were omitted where specified in the text. After incubation, all reactions treated with 1-3 µL RNAse A (NEB) for > 1 hr. Biotinylated substrates were prepared for UHPLC-MS analysis as described above. The 3 kbp substrates and oligomer substrates were purified using the Monarch® PCR and DNA Cleanup Kit with the DNA Cleanup and Concentration and the Oligonucleotide Cleanup protocols, respectively. The 3 kbp substrates and oligomer substrates were prepared for UHPLC-MS analysis as described above.

<u>Coupled Mom activity assay with oligomer substrates.</u>

The *in vitro* activity of Mom on short DNA was tested in Reaction Buffer (16 mM HEPES, pH 7.5, 40 mM NaCl, 40 mM KCl, 20 mM $MgCl_2$, 0.8 mM DTT) in 100 µL reaction volume. The mixture contained 17 µg of 11 bp or 3 bp DNA, 0.6 µM Mom protein (monomer), 1.6 µM tRNA, 0.09 µM GlyRS (hetero-tetramer), 10 mM glycine and 2 mM ATP. The reaction was incubated at 37 °C for 5 h followed by the digestion (*16*). The resulting ribo- and deoxyribonucleotides were resolved on the ACQUITY UPLC HSS T3 Column, 100 Å, 1.8 µm, 2.1 mm X 150 mm at 0.3 ml/min. Mobile phase was a linear gradient of 10 mM ammonium formate pH 4.4 (A) and methanol (B) (1.5 – 11.5 min to 10% B, 19 – 24 min to 100% B). Absorbance was measured at 260 nm.

Mom activity assay with $^{32}$P labelled dsDNA

10 pmoles of 34-mer DNA oligonucleotide (top strand) was 5′ labelled with γ-$^{32}$P ATP (Hartmann Analytics) and polynucleotide kinase (PNK, Thermo), annealed to the unlabeled complementary strand (1:1.2) and purified using MiniQuick Spin DNA column (Roche). 2 pmoles of radiolabelled dsDNA were mixed with 0.6 µM Mom protein (monomer), 1.6 µM tRNA, 0.09 µM GlyRS (hetero-tetramer), 10 mM glycine and 2 mM ATP in 100 µL of the Reaction Buffer and incubated at 37 °C overnight. The DNA was then purified by phenol-chloroform and precipitated by cold ethanol. Purified DNA was treated by AasI FastDigest (Thermo), and the digestion product were resolved on 20% Urea-PAGE and visualized by autoradiography using Amersham Typhoon Biomolecular Imager.

*In vitro* plasmid protection assay.

The *in vitro* activity of Mom and its variants was tested in Reaction Buffer (16 mM HEPES, pH 7.5, 40 mM NaCl, 40 mM KCl, 20 mM $MgCl_2$, 0.8 mM DTT) in 100 µL reaction volume. The mixture contained 3.6 µg of substrate plasmid (pET28a-empty), 0.6 µM Mom protein (monomer), 1.6 µM tRNA, 0.09 µM GlyRS (hetero-tetramer), 10 mM glycine and 2 mM ATP. The reaction was incubated at 37 °C for 5 hours followed by plasmid purification on silica spin columns (DNA clean-up kit, A&A Biotechnology). Purified DNA was subjected to the digestion with momylation-sensitive restriction endonucleases HgaI (NEB) or AasI FastDigest (Thermo Scientific™). The digestion products were resolved on 0.8% agarose gel and imaged using GelRed (Biotium) staining.

Charging, labeling, and analysis of Gly-tRNA$^{Gly/GCC}$.

tRNA$^{Gly/GCC}$ (100-300 µg [20-60 µM]) was aminoacylated (charged) for 20 minutes – 1 hour at 37 °C in 100 mM HEPES pH 7.6, 2 mM ATP, 20 mM $MgCl_2$, 2 mM glycine, 1 mM DTT, 2-10 µM GlyRS, and Pyrophosphatase, Inorganic (5 units) in a total volume of 200-500 µL (*17*, *18*). Reactions were quenched with 1:10-1:4 volumes of 3 M NaOAc pH 5. Quenched reactions were mixed 1:1 with cold phenol (pH 4.5) and shaken for 10 min at 4 °C, and samples were spun at 13000 rpm at 4 °C for 5 minutes. The phenol phase was back extracted twice with equal volume of 300 mM NaOAc pH 5. An equal volume of chloroform was added to combined aqueous fractions, shaken for 10 min at 4 °C, and spun at 13000 rpm at 4 °C for 5 minutes. The aqueous phase was back extracted once with chloroform. 2.5 volumes of cold ethanol (-20 °C) was added to combined aqueous fractions and incubated on dry ice for 30 minutes. Samples were centrifuged at full speed for 30 min. Pellets were washed twice with 4 volumes of ice cold 70% ethanol and spun at full speed for 10 minutes. After removing the supernatant, pellets were dried on a vacuum concentrator briefly. Pellets were resuspended in 5 mM NaOAc (pH 5.3) and passed over Microspin™ G-25 Columns (Cytiva) that had been pre-equilibrated with 5 mM NaOAc pH 5. The Qubit™ Broad Range Assay Kit (Invitrogen™) was used to quantify the tRNA recovered, and samples were stored at -80 °C until use.

tRNA recovered from the charging reaction (~150 pmol [~3.6 µg, 2.5 µM]) was labeled with EZ-Link™ Sulfo-NHS-LC-Biotin (Thermo Scientific™) that was freshly prepared in RNase-free water (15 mM) by incubating at 4 °C for 1 hour in 60 mM HEPES pH 8 in a total volume of 60 µL. Labeling reactions were cleanup by ethanol precipitation twice. Pellets were air-dried, resuspended in 12 µL of RNase-free water, and quantified by Qubit. Biotin-labeled tRNA (~8 pmol) was further labeled with 8 µL of streptavidin (1 mg/mL stock in water, Invitrogen™) in a total volume of 10 µL for 20 minutes at room temperature. 200 ng of samples were mixed 1:1 RNA Loading Dye (NEB) without heating and assessed by Urea PAGE gels as described above. ImageLab (Bio-Rad) was used to quantify band intensities of uncharged and charged (shifted) bands. Percent charging was calculated diving the intensity of the shifted band by the sum of the intensities of shifted and unshifted bands. Generally, 20-30% charging of $tRNA^{Gly}$ was observed.

Mom activity assays with Gly-$tRNA^{Gly/GCC}$.

Components for the coupled Mom activity assay were mixed on ice in the following order to a final volume of 30 µL and incubated 5 hours – overnight at 37 °C: activity buffer (50 mM HEPES pH 7.6, 100 mM KCl, 13 mM magnesium acetate, 1 mM DTT, and ± 2 mM ATP); Pyrophosphatase, Inorganic (0.025 units/mL, NEB); ± GlyRS (2-4 µM); glycine (2 mM); RNase Inhibitor, Murine (20 units, NEB); ± Mom (2 µM); biotinylated DNA substrates (500 ng); Gly-$tRNA^{Gly/GCC}$ (4-16 µM total tRNA, 1-4 µM charged tRNA); and RNase-free water. Uncharged $tRNA^{Gly/GCC}$ (4 µM) was used in control reactions, and where specified, $^{13}C$,$^{15}N$-glycine (2 mM, Sigma) was used in place of glycine. After incubation, biotinylated DNA substrates were prepared for UHPLC-MS analysis as described above.

Centrifuge filtration activity assay with Gly-$tRNA^{Gly/GCC}$.

In the filtration assay, tRNA was charged by GlyRS in 1 mL Reaction Buffer that contained 0.72 µM $tRNA^{Gly/GCC}$, 0.018 µM GlyRS (hetero-tetramer), 10 mM glycine, 2 mM ATP. After a 15 min incubation at 37 °C, the solution was slowly (3000 g) filtered using 100 kDa MWCO Amicon Ultra-0.5 Centrifugal Filter Unit (Merck). During centrifugation, the flow-through containing charged $tRNA^{Gly/GCC}$ was immediately mixed with 150 µL of the Reaction Buffer containing Δ10 Mom (1 µM) and the substrate plasmid (5 µg pET28-empty) that added to the bottom reservoir of the filter unit before centrifugation. As a control to test for possible GlyRS leakage through the membrane, the flow-through of the solution containing only GlyRS was mixed with the rest of the components in the same setting. After filtering ~950 µL of tRNA charging solution, the resulting mix was incubated for another 2 h at 37 °C. The plasmid was then purified on silica spin columns (DNA clean-up kit, A&A Biotechnology) and subjected to HgaI (NEB) digestion. The digestion products were resolved on 0.8% agarose gel and imaged using GelRed (Biotium) staining.

Mom crystallization and structure determination.

Purified Δ10 Mom S124A protein was applied on a Superdex 200 Increase 10/300 GL column equilibrated with Buffer 9 (20 mM HEPES-KOH, pH 7.5, 550 mM NaCl, 50 mM KCl, 5% glycerol, 10 mM 2-ME). The best fractions were combined and concentrated to ~23 mg/mL using Amicon 10 MWCO centrifugal filter unit. Mom crystals were grown by mixing 2 µL of the protein with 2 µL of the condition H4 of the PACT premier crystal screen (Molecular Dimensions, Sheffield, England) (0.2 M Potassium Thiocyanate, 0.1 M Bis–Tris propane, pH 8.5, 20% PEG3350) using the vapor diffusion method in hanging drops. For cryo-protection, crystals were soaked in mother liquor containing 25% glycerol and flash-cooled in liquid nitrogen. Mom crystals diffracted up to approximately 2 Å on PETRAIII P11 beamline of the DESY synchrotron. The diffraction data was automatically processed by the XDSAPP program at the synchrotron (*19*). The structure of Mom was solved by molecular replacement using the Phaser program and the Alphafold 2 predicted model (*20*, *21*). The programs Phenix and COOT were used for the refinement (*22*, *23*). The data collection and refinement statistics are presented in Supplementary Table S3. The final model coordinates and the structure factors were deposited in Protein Data Bank under accession code: 8BV8.

3D structural analysis and graphic representation

Structure manipulation and image rendering was performed using UCSF ChimeraX program (*24*). The MatchMaker tool in ChimeraX was used for structure superposition. Secondary structure assessment (Fig. 3B and 3C) was guided by the PDBsum server (*25*). Vector Alignment Search Tool (VAST) was used to search for the structural similarity between Mom and the available GNAT structures on the medium redundancy setting (*26*). GNATs with productively bound CoA and acyl-CoA were used for superposition in Fig. 3E and were selected from the top 164 hits (the cut-off was manually determined). The list is provided below.

| **PDB ID/ chain** | **Description** | **Organism** |
|---|---|---|
| 4ri1/A | Pseudaminic acid biosynthesis N -acetyltransferase PseH | *Helicobacter pylori* |
| 4u9w/C | N-terminal acetyltransferase NatD | *Homo sapiens* |
| 1yre/D | Hypothetical protein PA3270 | *Pseudomonas aeruginosa* |
| 5t7e/C | Bialaphos Resistance (BAR) protein | *Streptomyces hygroscopicus* |
| 5us1/C | Aminoglycoside acetyltransferase AAC(2')-Ia | *Providencia stuartii* |
| 4pv6/I | Protein N-acetyltransferase Ard1 | *Thermoplasma volcanium* |
| 2psw/B | MAK3 homolog | *Homo sapiens* |
| 1s7n/B | Ribosomal L7/L12 alpha-N-protein acetyltransferase | *Salmonella* Typhimurium |
| 6ag5/A | N-terminal acetyltransferase Ard1 mutant E88H/H127E | *Sulfolobus solfataricus* |
| 2zw7/A | Bleomycin N-acetyltransferase | *Streptomyces verticillus* |
| 2zpa/A | $tRNA^{Met}$ cytidine acetyltransferase | *Escherichia coli* |
| 6edv/A | GNAT superfamily acetyltransferase PA3944 | *Pseudomonas aeruginosa* |
| 3fbu/A | GNAT family acetyltransferase | *Bacillus anthracis* |
| 3owc/A | GNAT superfamily protein PA2578 | *Pseudomonas aeruginosa* |
| 2vqy/A | Aminoglycoside N-acetyltransferase Aac(6')-Ib | *Escherichia coli* |
| 1wwz/B | PH1933 | *Pyrococcus horikoshii* |
| 2ge3/C | Probable acetyltransferase | *Agrobacterium tumefaciens* |
| 3r9g/A | Microcin C7 self-immunity acetyltransferase MccE | *Escherichia coli* |
| 1tiq/A | N1-spermidine/spermine acetyltransferase (PaiA) | *Bacillus subtilis* |
| 2jdd/A | Glyphosate N-acetyltransferase | *Bacillus licheniformis* |
| 2ref/A | GNATL domain of CurA | *Lyngbya majuscula* |
| 5ktd/A | dTDP-3-amino-3,6-dideoxy-d-galactose acyltransferase FdhC | *Acinetobacter nosocomialis* |
| 4ua3/A | N-terminal acetyltransferases NatD | *Schizosaccharomyces pombe* |
| 2cy2/A | TTHA1209 | *Thermus thermophilus* |
| 2i79/E | GNAT family acetyltransferase | *Streptococcus pneumoniae* |

| | | |
|---|---|---|
| 2fiw/A | GCN5-Related N-acetyltransferase: Aminotransferase | *Rhodopseudomonas palustris* |
| 6wn0/A | Acyl-homoserine lactone synthase RpaI | *Rhodopseudomonas palustris* |
| 1m4i/A | Aminoglycoside 2'-N-acetyltransferase | *Mycobacterium tuberculosis* |
| 5gi9/A | Dopamine N-Acetyltransferase | *Drosophila melanogaster* |
| 3zj0/A | O-GlcNAcase | *Oceanicola granulosus* |
| 4kvx/B | N-terminal acetyltransferase Naa10 (Ard1) | *Schizosaccharomyces pombe* |
| 3exn/A | Probable acetyltransferase | *Thermus thermophilus* |
| 3qb8/A | Polyamine acetyltransferase A654L | PBCV-1 |
| 2a4n/A | Aminoglycoside 6'-N-acetyltransferase type Ii | *Enterococcus faecium* |
| 1j4j/B | Tabtoxin Resistance Protein | *Pseudomonas amygdali* |
| 6add/B | l-glutamate/l-glutamine acetyltransferase ArgA | *Mycobacterium tuberculosis* |
| 5hmn/E | Aminoglycoside acetyltransferase HMB0005 | uncultured bacterium |
| 3pp9/B | Putative streptothricin acetyltransferase | *Bacillus anthracis* |
| 1qsm/C | Histone Acetyltransferase HPA2 | *Saccharomyces cerevisiae* |
| 3f8k/A | Protein acetyltransferase (PAT) | *Sulfolobus solfataricus* |
| 5k18/F | N-Terminal Acetyltransferase NatB | *Candida albicans* |
| 5wjd/A | N-terminal acetyltransferase NAA80 | *Drosophila melanogaster* |
| 3mgd/A | Predicted acetyltransferase | *Clostridium acetobutylicum* |
| 1p0h/A | Mycothiol synthase (Rv0819) | *Mycobacterium tuberculosis* |
| 6gtp/A | AtaT (tRNA acetylating toxin) | *Escherichia coli* |
| 5fvj/A | TacT (tRNA acetylating toxin) | *Salmonella* Typhimurium |
| 4zbg/A | GNAT family Acetyltransferase | *Brucella melitensis* |
| 6g96/A | TacT3 (tRNA acetylating toxin) | *Salmonella* Typhimurium |
| 6tdg/A | Glucosamine-6-phosphate N-acetyltransferase 1 | *Aspergillus fumigatus Af293* |
| 2q4v/A | Thialysine n-acetyltransferase (SSAT2) | *Homo sapiens* |
| 4h6u/A | Tubulin acetyltransferase (TAT) | *Danio rerio* |

Computational Methods.

Geneious Prime software (v 2020.1.1) was used to run a Basic Local Alignment Search Tool (BLAST) search of the Nucleotide collection (nr/nt) for homologs of Mu Mom, which retrieved 500 Mom homologs (*4*, *5*, *27*, *28*). Sequences were trimmed from hits that appeared to have a Mom-like domain fused to other domains. An HMM profile for Mom was built using the hmmbuild program from the HMMER software package (*29*). The hmm profile for Mom was used to search two metavirome sequence databases: the Joint Genome Institute's Integrated Microbial Genomes Viral Resource version 2 (IMG/VR v2) (*30*) and the Global Oceanic Viromes 2.0 (GOV2.0) (*31*). IMG/VR2 contains 715 672 contigs encoding 16 215 899 proteins computationally identified as viral and retrieved from metagenome sequences archived at IMG. Global Oceanic Viromes 2.0 (GOV2.0) is an environmental meta-metagenome dataset encompassing 145 viromes sampled from the world's oceans and containing 848,507 contigs encoding 12,486,732 proteins (sequence data available at https://datacommons.cyverse.org/browse/iplant/home/shared/iVirus/GOV2.0). Retrieved contigs (> 8000) were then obtained from the databases by text matching the protein ID to the contig ID. Open reading frames in the recovered contigs were predicted and annotated using Prokka (*32*) within JGI's KBase Knowledge Discovery Environment (*33*). A more thorough annotation process using the Pfam database (*34*) was completed using the hmmscan program. Contigs were visualized using Geneious Prime.

Sequence similarity network (SSN) for Mu Mom and predicted Mom homologs.
All-against-all BLAST comparisons of protein sequences including Mu Mom and >8000 predicted homologs were run using EFI-EST (https://efi.igb.illinois.edu/efi-est/) (Enzyme Function Initiative – Enzyme Similarity Tool) (*35*) with an alignment score threshold of 35 (*36*). To facilitate visualization, the final network generated was 95% representative, collapsing sequences of 95% identity into representative nodes (3852 nodes, 598,234 edges, representative of 9,723 sequences). The similarity network was visualized using Cytoscape (*37*). The alignment threshold and representative node percentage reported were selected based on empirical analysis. Lower thresholds yielded more edges but did not provide distinct clustering (not shown). Alternatively, increasing the percent identity used as the cutoff to generate the representative node network increased the total number of singletons (not shown).

*In vivo* activity screen of Mom homologs.
The SSN for Mom and homologs was used to guide initial screening *in vivo* for activity. Sequences were selected at random from larger clusters and ordered with codon optimization for expression in *E. coli* from GenScript inserted into a pET 28 expression vector with an N-terminal $His_6$ tag (*4*, *27*, *28*). Homologs were expressed in 24-well plates with 2 mL of culture (10 mL well capacity), in glass tubes with 5 mL cultures (29 mL tube capacity), or in flasks where the culture volume was ~20% of the flask volume (volumes varied, specified where appropriate). Growth and expression conditions were described as above. Expression plasmids were harvested and subjected to nucleoside analysis by UHPLC-MS as described above.

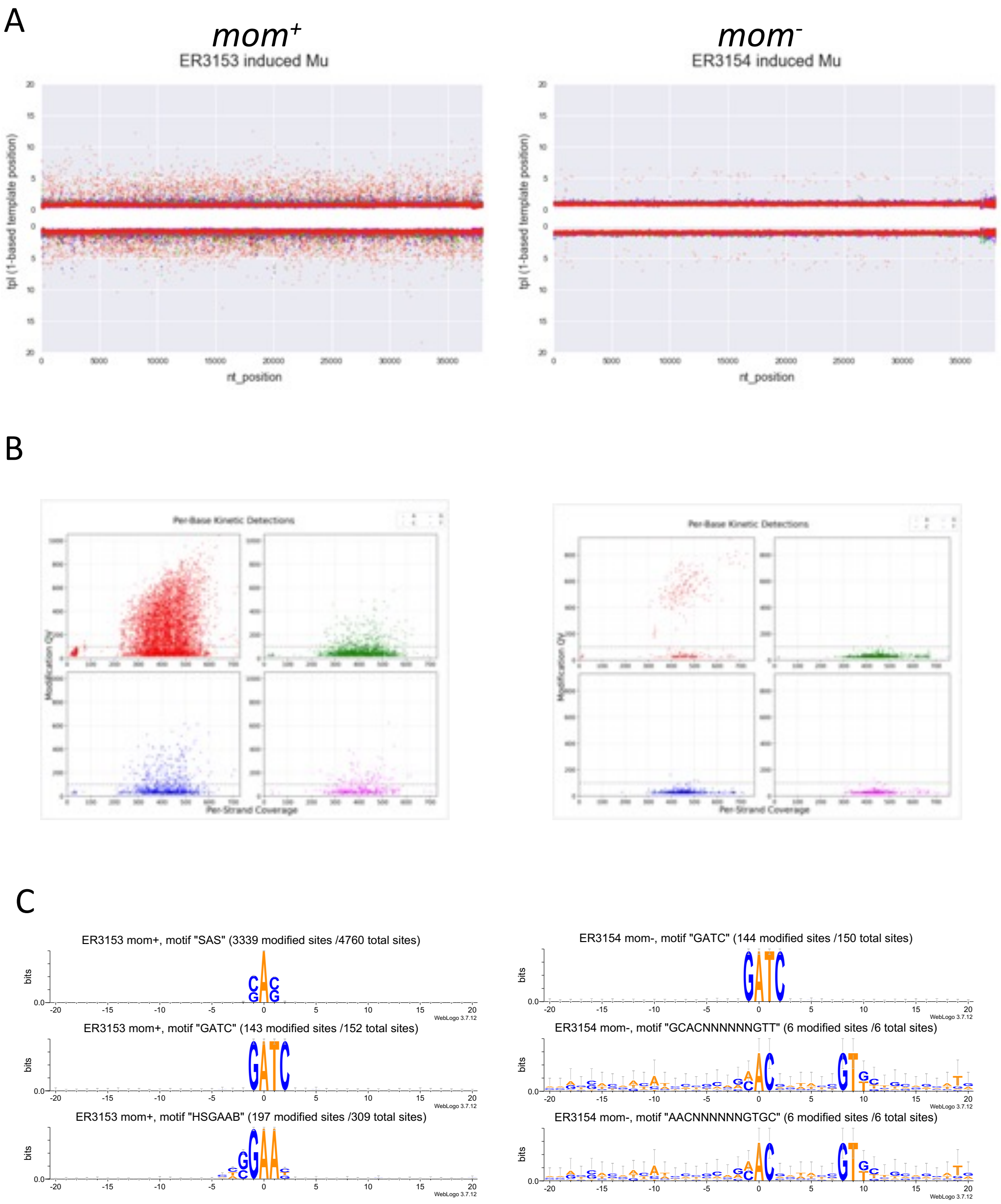

**Fig. S1. Phage Mu modifies dA in the context of an SAS sequence motif.**

DNAs extracted from phage Mu virions containing modified dA (ER3153) or from modification-defective virions (ER3154) were sequenced using PacBio SMRT technology followed by modification detection using SMRT Link (version 10.2.0) Base Modification and Motif Analysis tool. (**A**) The per-base IPD ratios for modified and unmodified sequencing templates were mapped to the reference sequence revealing extensive sites of increased IPD across the entire length of the modified DNA. (**B**) Per base kinetic detection plots show base A as the source of the majority of IPD signals. (**C**) Top three most abundant modification motifs detected in *mom*$^{+}$ and *mom*$^{-}$ Mu DNA represented as sequence logos generated using the WebLogo 3 webtool (version 3.7). See also Table S1.

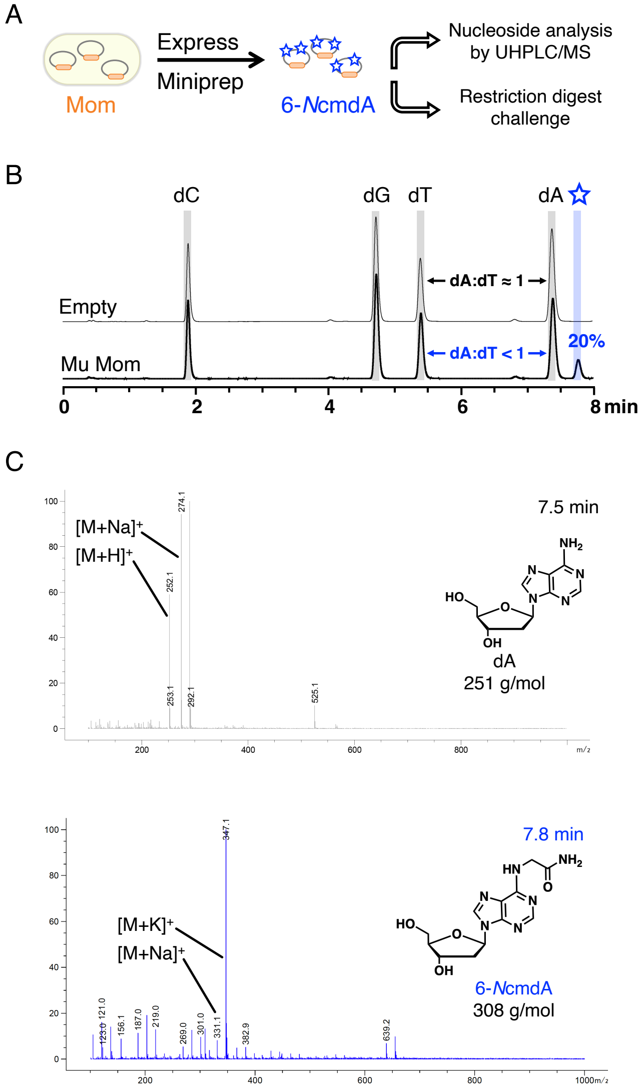

**Fig. S2. 6-*N*cmdA is detected following heterologous expression of Mu Mom in *E. coli*.**

(**A**) *In vivo* activity assay workflow for Mu Mom. Previously, Karambelkar and coworkers established that *mom* is the only Mu gene required to modify dsDNA in an *in vivo* system (*3*). Following expression of Mu Mom from a pRY vector in T7 Express (see Methods for expression conditions), hypermodified expression plasmids can be isolated and subjected to nucleoside analysis by HPLC/MS to detect 6-*N*cmdA. Hypermodified plasmids can alternatively be challenged with restriction endonucleases that are blocked by hypermodification at dA (ex. HgaI), and protected dsDNA can be visualized by gel electrophoresis (ex. see Fig. 1). (**B**) Resolution and (**C**) confirmation of 6-*N*cmdA by UHPLC-MS. Nucleoside digests of hypermodified plasmids were resolved by UHPLC. Using this method, a novel peak appears at ~7.8 minutes after heterologous expression of Mu Mom. This novel peak could be separated at baseline resolution from the canonical nucleosides and corresponds to greater than 20% depletion of dA. Mass spectrometry was used to assign the novel peak at ~7.8 minutes to 6-*N*cmdA (MW: 308 g/mol) (*3*, *38*).

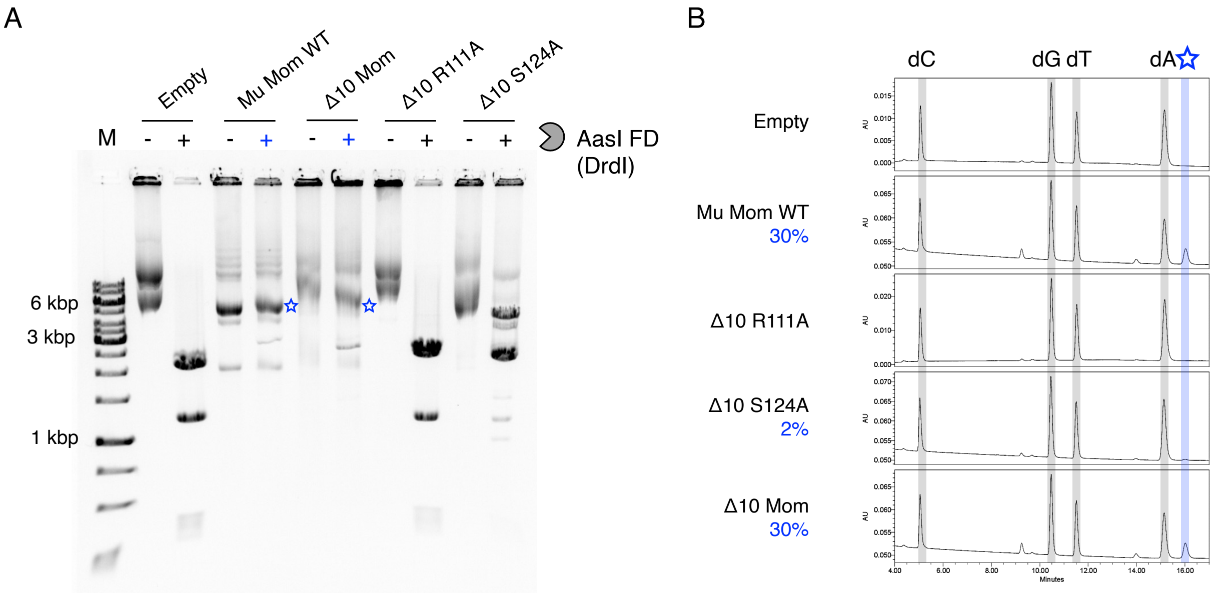

**Fig. S3.** ***In vivo* activity of Mu Mom and variants.**

(**A**) After IPTG induction and heterologous expression of Mom in *E. coli* strain BL21(DE3)-RIL, pET28a plasmids were isolated and subjected to a restriction digest challenge. “Empty” is a pET28a plasmid without a Mom ORF. (**B**) The same plasmids were digested to deoxyribonucleosides and analyzed by UHPLC. The percent conversion of dA to 6-*N*cmdA is indicated in blue. As with the “Empty” plasmid, the Mom mutant Δ10 R111A has no detectable activity, therefore, no conversion value is reported. The activity of Δ10 S124A was much reduced compared to WT Mu Mom (full-length) and Δ10 WT, as expected (*3*).

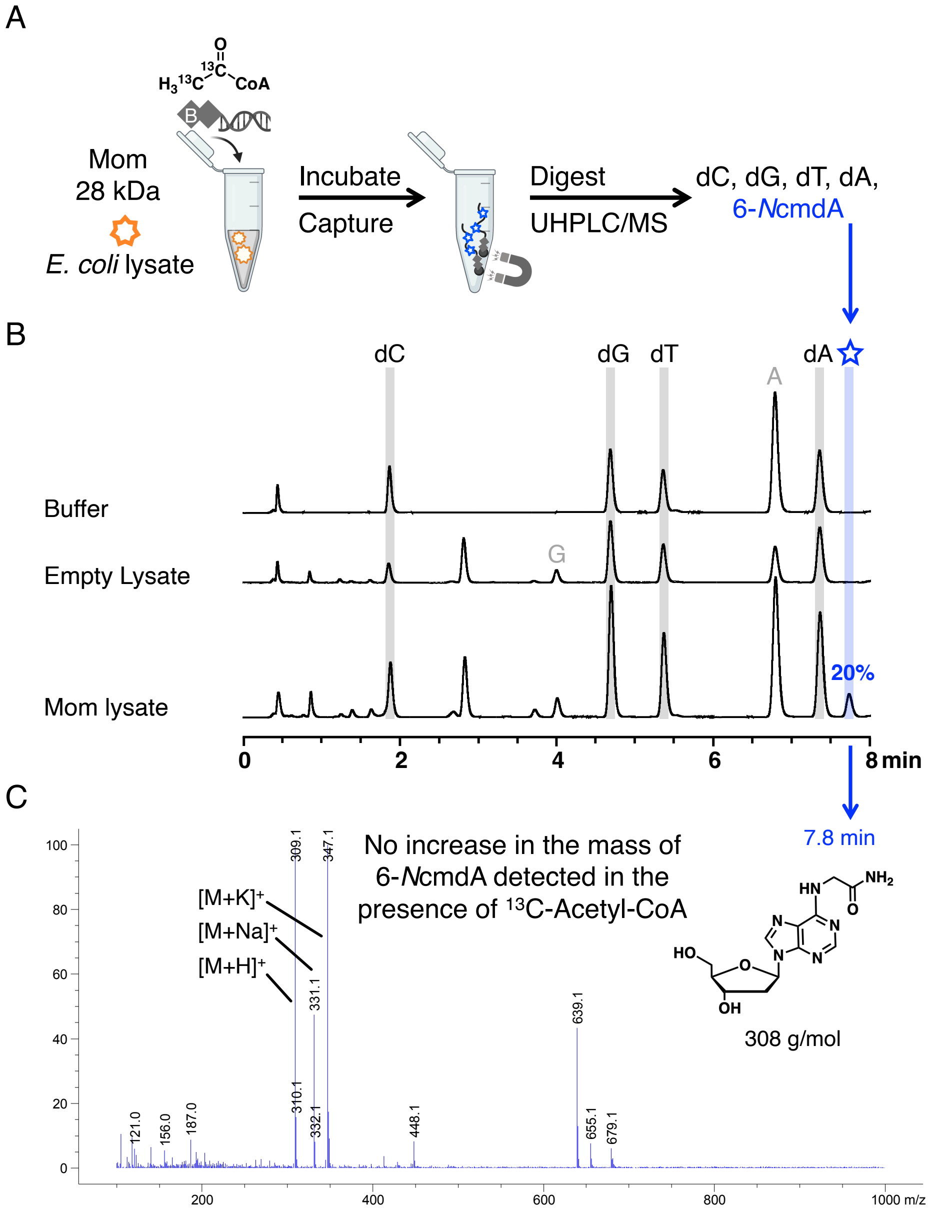

**Fig. S4. Labeled atoms were not transferred in an *ex vivo* Mom activity assay with $^{13}C_2$-acetyl-CoA**

GNAT-family enzymes commonly use acetyl coenzyme A (acetyl-CoA) as a co-substrate (*39–41*). While simple transfer of acetyl-CoA to dA would not result in 6-*N*cmdA, the possibility that acetyl-CoA could be the co-substrate of Mom was explored previously (*3*). A biosynthetic pathway for 6-*N*cmdA was proposed, where acetyl-CoA is transferred and oxidized by Mom through iron chemistry via an Elongator-like radical mechanism, followed by reduction and amidation steps accomplished by unidentified enzymes. In that study, a specific iron coordination site in the sequence of Mom eluded identification, and biosynthesis of 6-*N*cmdA was not observed *in vitro* with purified Mom. To the best of our knowledge, it is unprecedented for a GNAT family enzyme to utilize radical chemistry to acylate target substrates. This, it remained unclear if acetyl-CoA is not used to form 6-*N*cmdA or if the purified enzyme was inactive. As such, we utilized our *ex vivo* lysate assay (**A**), where Mom is active and converts ~20% of dA to 6-*N*cmdA (see Methods for conversion calculation), supplemented with isotopically-labeled $^{13}C_2$-acetyl-CoA to test if 6-*N*cmdA with an increased mass could be detected (**B**) (*4*, *5*). (**C**) 6-*N*cmdA produced in *ex vivo* reactions including dsDNA and $^{13}C_2$-acetyl-CoA contained no increase in mass. Nucleoside digests of biotinylated DNA substrate (500 ng) were analyzed by LC-MS following incubation with Mom lysate (25 µg total protein) and 1 mM $^{13}C$-acetyl-CoA (comparable to the intracellular concentration of acetyl-CoA, BNID 114622) (*42*). 6-*N*cmdA formed with the same mass as reactions without supplemented cofactor, suggesting the modifying atoms are not derived from acetyl-CoA.

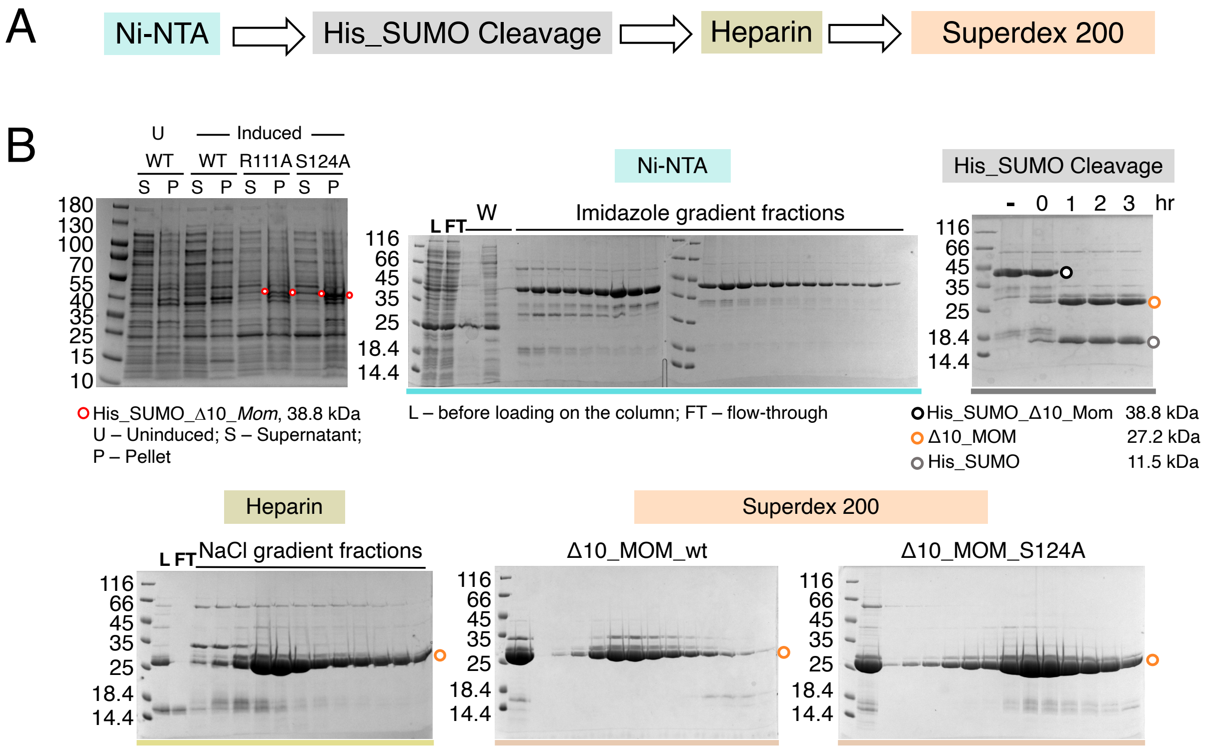

**Fig. S5. Expression and purification of wild-type Mom and variants.**
(**A**) Outline of the purification steps. WT Mom and S124A were expressed with an N-6His-SUMO tag. After affinity purification over an Ni-NTA column (5 mL His-Trap, Cytiva), the 6His-SUMO tag was cleaved by SUMO protease. Cleavage was followed by purification over Heparin Sepharose resin (5 mL Hi-Trap Heparin, Cytiva). In the final step, Mom was resolved using a size-exclusion column (Superdex 200 Increase 10/300 GL, Cytiva). (**B**) Purification progress monitored by SDS-PAGE.

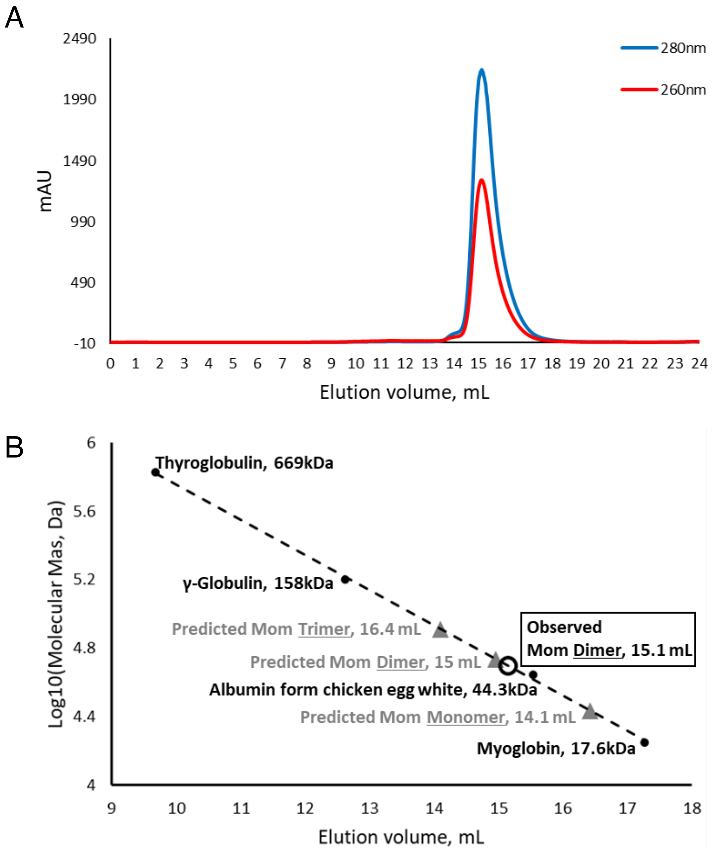

**Fig. S6. Estimation of Mom quaternary structure using size-exclusion chromatography.**

(**A**) The elution profile of Mom Δ10 from a Superdex 200 Increase 10/300 GL column was monitored by the UV absorption at 280 nm (blue trace) and 260 nm (red trace). (**B**) Determination of homo-dimeric structure of Mom in solution. Gel filtration standards (Bio-Rad) were used for calibration. Decadic logarithms of the molecular weights of the standards (y axis) were plotted against their elution volumes (x-axis). Predicted elution volumes for Mom monomer, dimer, and trimer (gray triangles) were calculated based on the molecular weight. The elution volume of Mom (black circle) strongly suggests a dimeric structure in solution as reported previously (*3*).

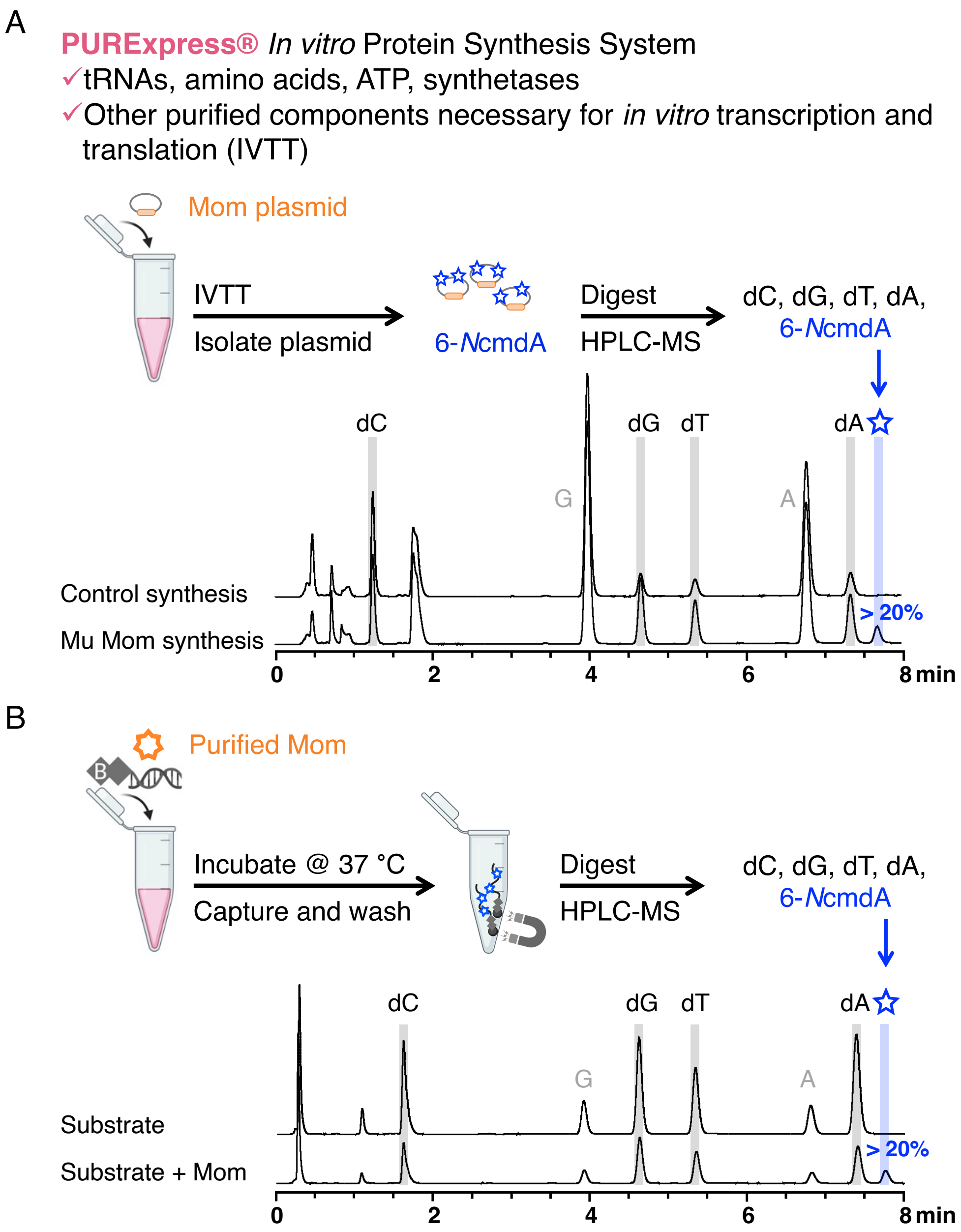

**Fig. S7. Adenine hypermodification to 6-*N*cmdA requires components of transcription and/or translation.**

(**A**) Reconstitution of 6-*N*cmdA by IVTT of Mom in the PURExpress® system. We aimed to reconstitute the momylation reaction *in vitro* using experimental approaches amenable to systematic dissection of reaction components, including potential group donors. We added 1 µg of pRY or pET28a expression plasmid encoding for Mom to the PURExpress® reconstituted protein synthesis system and incubated for 24 hours at 25 ºC. PURExpress® includes purified enzymes and cofactors sufficient for transcription and translation, including tRNA, amino acids, and tRNA synthetases. The control synthesis reaction was incubated according to the manufacturer's instructions and was used as a negative control for hypermodification. Expression plasmids were recovered after incubation and subjected to LC-MS as described. UHPLC traces and mass spectra (not shown) confirmed that *in vitro* transcription and translation of the *mom* gene leads to installation of 6-*N*cmdA, indicating that co-substrate used by Mom is a component of transcription and/or translation. We note that Mom was able to install 6-*N*cmdA without the addition of $Fe^{2+}$ or acetyl-CoA. The trace for Mu Mom Synthesis comes from an IVTT reaction using a pRY Mu Mom expression plasmid as the template. (**B**) Reconstitution of 6-*N*cmdA with purified Mom, dsDNA, and PURExpress® components *in vitro*. Purified Mom was mixed with biotinylated DNA substrates in the PURExpress® system in order to test if Mom was purified in an active state and could install 6-*N*cmdA in the absence of active protein synthesis. After incubation at 25 ºC overnight, the biotinylated substrate was recovered, digested to nucleosides, and analyzed. UHPLC traces and mass spectra (not shown) confirmed that the activity of purified Mom resulted in formation of 6-*N*cmdA. Previously, we and others noticed that the exocyclic group of 6-*N*cmdA and a glycyl group are structural isomers (*3*). Since there is considerable precedent for certain GNAT-family enzymes to aminoacylate target substrates using charged tRNAs (ex. Gly-$tRNA^{Gly}$) as group donors, we hypothesized that Gly-$tRNA^{Gly}$ could be the co-substrate used by Mom to hypermodify dA. This hypothesis was first supported experimentally with the PURExpress® activity assays (see main text for further discussion) because we observed Mom was active in the presence of Gly-$tRNA^{Gly}$, which is included in the PURExpress® kit as a necessary component for translation.

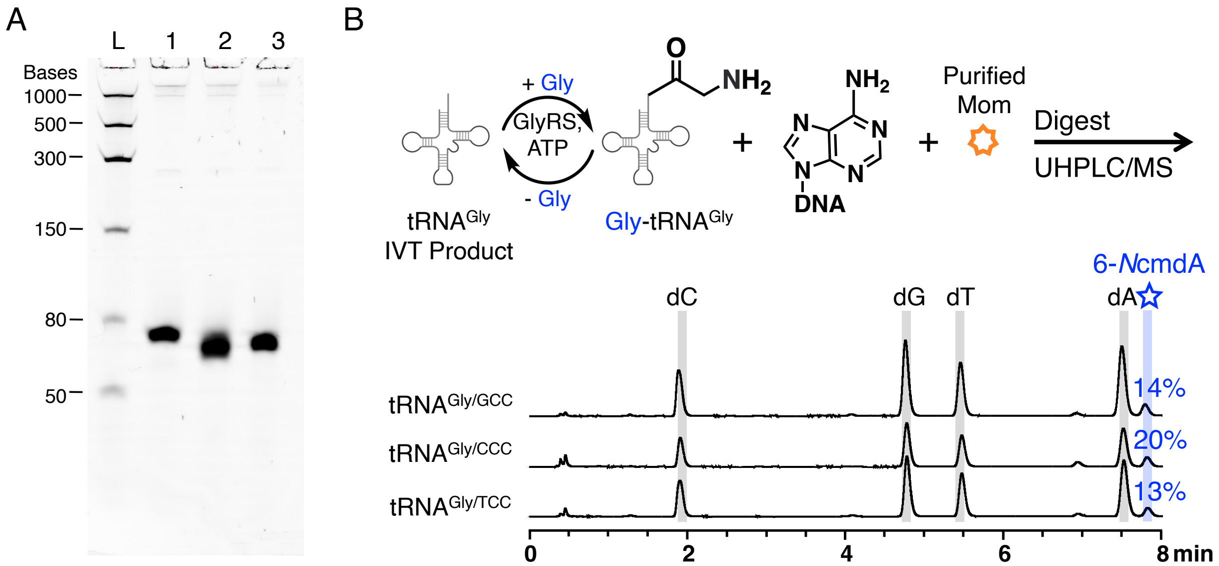

**Fig. S8. All tRNA$^{Gly}$ isoacceptors from *E. coli* support momylation**.

(**A**) Synthesis of tRNA$^{Gly}$ isoacceptors. *E. coli* encodes for three different isoacceptors of tRNA$^{Gly}$ (see table S2 for sequences) – tRNA$^{Gly/GCC}$ (76 bases, Lane 1), tRNA$^{Gly/CCC}$ (74 bases, Lane 2), and tRNA$^{Gly/TCC}$ (74 bases, Lane 3). tRNA$^{Gly/GCC}$ is reported to be the most frequently-used and most abundant isoacceptor in *E. coli* (*9*, *10*, *43*). Since there is precedence for a particular tRNA isoacceptor to be used in a non-canonical role outside of ribosome-dependent protein synthesis (*44*, *45*), we were interested in testing the co-substrate specificity of Mom. Each isoacceptor was synthesized by *in vitro* transcription, purified, and assessed on a denaturing 6% urea-TBE gel (~200 ng/lane, 200 V, 30 minutes). (**B**) Coupled activity assay with Mom and tRNA$^{Gly}$ isoacceptors. We developed a coupled activity assay containing purified Mu Mom, DNA, and components to generate Gly-tRNA$^{Gly}$ *in situ* – tRNA$^{Gly}$, glycine, ATP, and purified glycyl-tRNA synthetase (GlyRS) from *E. coli* (*15*). As can be seen from the HPLC traces of each reaction, all isoacceptors of tRNA$^{Gly}$ supported the momylation reaction. This finding is consistent with the strategy of the viral lifecycle, where available resources are scavenged rapidly in the process of hijacking the host cellular machinery. Because all tRNA$^{Gly}$ isoacceptors support momlyation, we speculate that Mom could be recognizing a small portion of charged tRNA, most likely at the 3′ acceptor end (5′-tRNA-CCA-Gly).

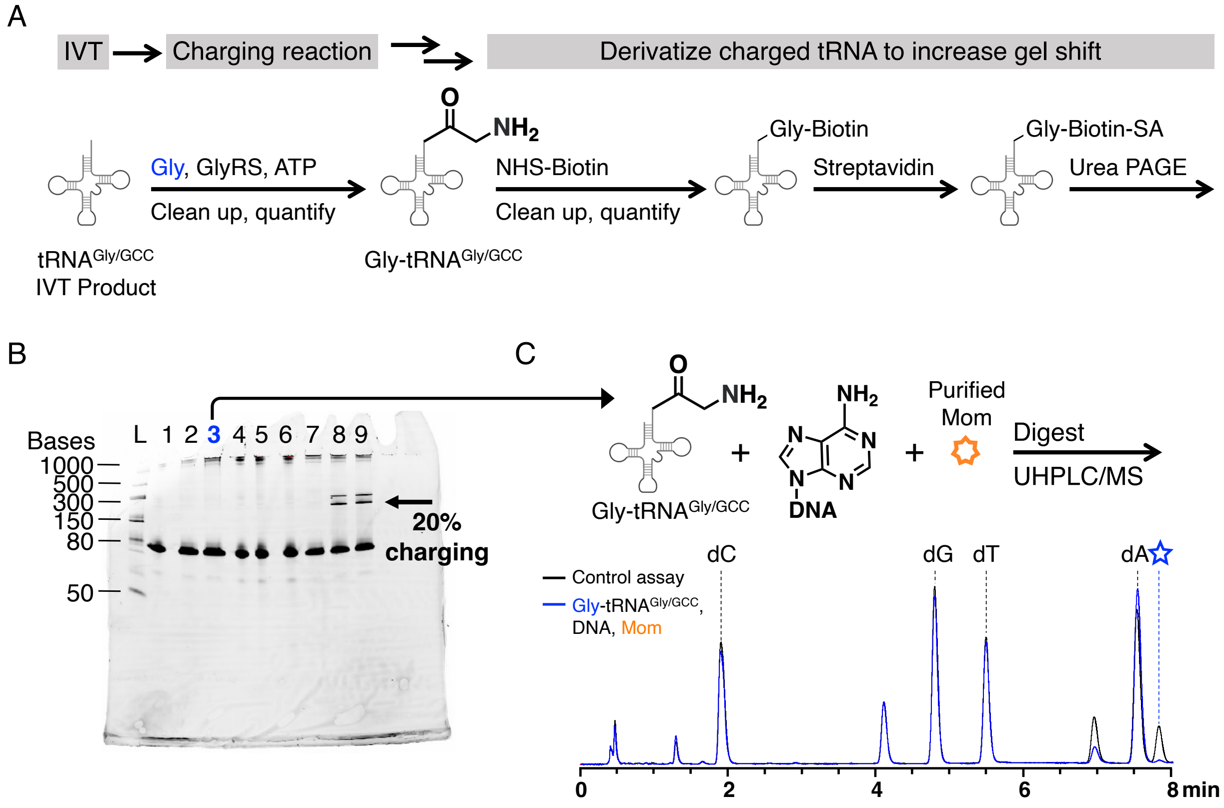

**Fig. S9. Mom can hypermodify dA using Gly-tRNA$^{Gly}$ in the absence of GlyRS.**

(**A**) Charging, purification, and derivatization of Gly-tRNA$^{Gly/GCC}$. Following reconstitution of 6-*N*cmdA in our coupled reaction, we sought to determine if GlyRS had any role, other charging Gly-tRNA$^{Gly}$, in momlyation. We started by testing if momlyation could proceed if GlyRS was excluded from the reaction. We purified Gly-tRNA$^{Gly/GCC}$ by phenol-chloroform extraction and ethanol precipitation following a charging reaction that included tRNA$^{Gly/GCC}$, glycine, GlyRS, and ATP (*13*, *17*, *46*) and used methods developed by Gamper and Hou to confirm generation of Gly-tRNA$^{Gly/GCC}$ (*18*). (**B**) Quantification of Gly-tRNA$^{Gly/GCC}$ yields by urea PAGE. Charging, purification, and derivatization of Gly-tRNA$^{Gly/GCC}$ was monitored by 15% TBE-Urea PAGE (200 ng/lane). L: Ladder, Lane 1: tRNA$^{Gly/GCC}$; Lane 2: Gly-tRNA$^{Gly/GCC}$ post-extraction and precipitation; Lane 3: Gly-tRNA$^{Gly/GCC}$ post-column purification; Lane 4: Lane 1 incubated with NHS-Biotin; Lane 5: Lane 2 incubated with NHS-Biotin; Lane 6: Lane 2 incubated with NHS-Biotin; Lane 7: Lane 4 incubated with streptavidin; Lane 8: Lane 5 incubated with streptavidin; Lane 9: Lane 6 incubated with streptavidin. Yields of Gly-tRNA$^{Gly/GCC}$ were determined by quantifying bands using Bio-Rad ImageLab software and found to be consistently around 20% (intensity of top migrated bands/[top + bottom migrated bands]). We attribute the top, double banding to multimers of streptavidin that were not fully denatured (*47*). (**C**) Mom can install 6-*N*cmdA in the absence of GlyRS. Incubating Mom with 20% Gly-tRNA$^{Gly/GCC}$ and the biotinylated dsDNA substrate led to conversion of dA to 6-*N*cmdA (2% conversion). We believe that this level of reaction is unlikely to be caused by a GlyRS contamination and suspect that the reduced conversion rate compared to the reaction with GlyRS, Gly-tRNA$^{Gly/GCC}$ and ATP (which is generally ≥ 20%) is likely a result of incomplete tRNA acylation. Together with prior data on other GNAT acyltransferases acting on tRNA, our findings favor the model that Mom harvests activated glycine from the charged Gly-tRNA$^{Gly}$, and GlyRS is otherwise a spectator during momylation.

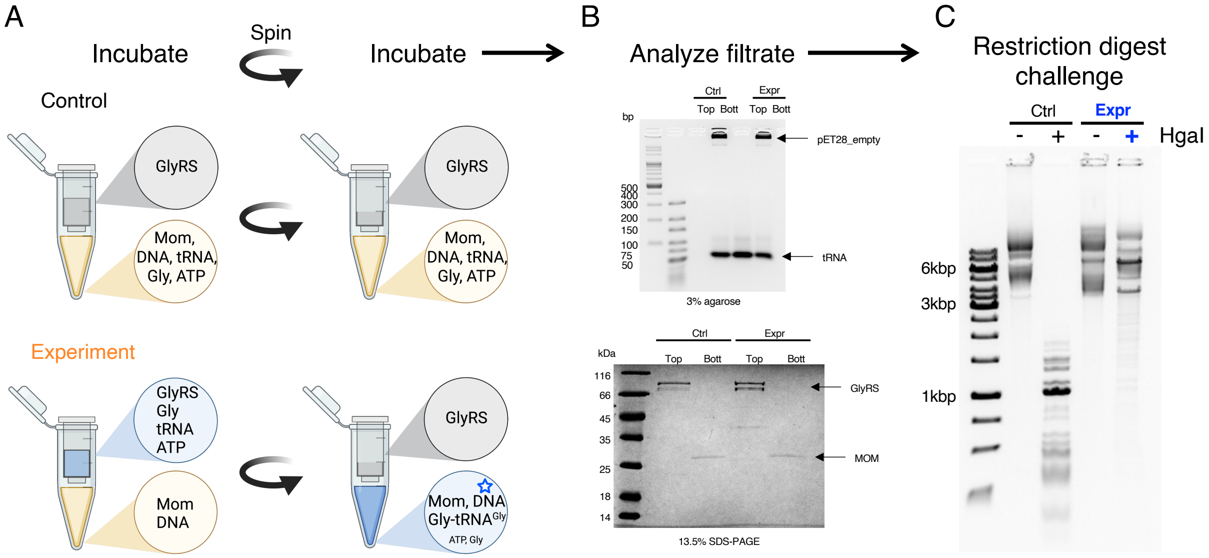

**Fig. S10. Charged tRNA$^{Gly}$ is sufficient for momylation reaction: the filtration assay.**
(**A**) tRNA charging by GlyRS and DNA modification by Mom were separated in space by filtration. The top reservoir of the 100 kDa molecular weight cut-off (MWCO) centrifugal filter unit (Amicon Ultra-0.5, Merck) included tRNA$^{Gly}$ and the components necessary for its aminoacylation (GlyRS, ATP, and glycine). The bottom reservoir included a DNA substrate (pET28a empty plasmid) and the Mom enzyme. After incubation, centrifugation allowed Gly-tRNA$^{Gly}$ (24.5 kDa) but not GlyRS (225.3 kDa) to pass through filter and immediately mix with Mom and DNA in the bottom reservoir. To exclude that GlyRS could pass through the membrane, a control reaction was set up that had only GlyRS in the top reservoir and all other components of the momylation reaction in the bottom reservoir. (**B**) Electrophoretic analysis of the filtrate after incubation and mixing with the bottom solution. Resolution of the nucleic acids on a 3% agarose gel (top gel) demonstrates that tRNA can pass through the filter. Conversely, SDS-PAGE analysis (bottom gel) shows that GlyRS is retained in the top reservoir after spinning. (**C**) Restriction digest challenge of substrate plasmids. Near complete protection is observed in the experiment but not in a negative control, suggesting that charged tRNA$^{Gly}$ alone is the co-substrate of Mom and is sufficient for supporting momlyation.

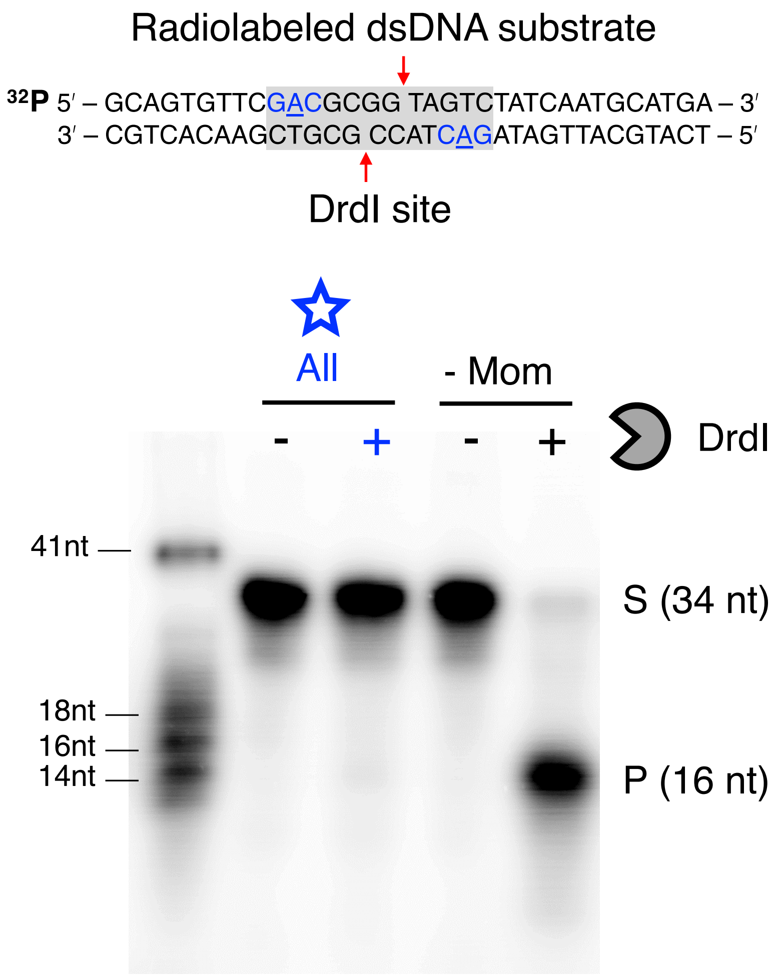

**Fig. S11.** ***In vitro*** **activity of Mom on a 34bp dsDNA substrate.**

A 34-mer oligo with a single recognition site for DrdI was labelled at the 5′ end with $^{32}P$ and annealed to an unlabeled complementary strand. After the overnight incubation with Mom, GlyRS, $tRNA^{Gly}$, ATP, and glycine, DNA was purified by phenol-chloroform extraction and ethanol precipitation. Purified DNA was challenged with AasI (DrdI) FastDigest (Thermo), digestion products were resolved by 20% Urea-PAGE, and visualized by autoradiography. Momylation renders 34bp dsDNA completely resistant to DrdI cleavage.

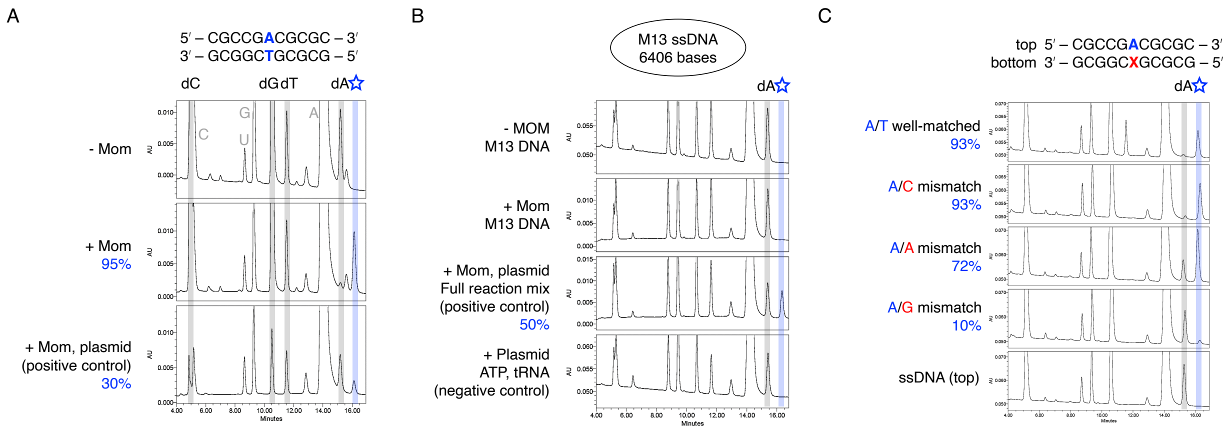

**Fig. S12. *In vitro* Mom activity on short dsDNA, single-stranded DNA, and mismatched DNA: UHPLC detection.**

(**A**) Mom hypermodifies 11 bp DNA containing a single adenine in the SASNY context with a very high conversion rate (95%). Due its small length, the substrate DNA was not purified prior the analysis, and therefore, the chromatogram contains traces of ribonucleotides originating from tRNA and ATP. As a positive control, pET28a plasmid was used as a substrate and was also not subjected to the purification. An unknown peak between dA and 6-*N*cmdA is likely an artifact caused by rA overload of the column. (**B**) Mom activity on single-stranded DNA (ssDNA) from phage M13 could not be detected above background. As in a panel A, substrate DNA was not purified prior analysis. Reactions with the pET28a plasmid with or without Mom added were used as a positive and negative controls, respectively. (**C**) Mom activity on well-matched (WM) or mismatched (MM) 11bp DNA. Mom exhibits activity at A/C MM (93% conversion) and to a lesser extent at an A/A MM (72% conversion). Activity on an A/G MM is diminished compared to the perfectly paired control (10%). Mom is completely inactive on 11 nt ssDNA, which is consistent with the data shown in panel B. Taken together, we conclude that Mom is active on dsDNA (even on very short oligomers) and inactive on ssDNA. We attribute any residual activity on M13 ssDNA as a likely result of Mom binding to partially double-stranded regions.

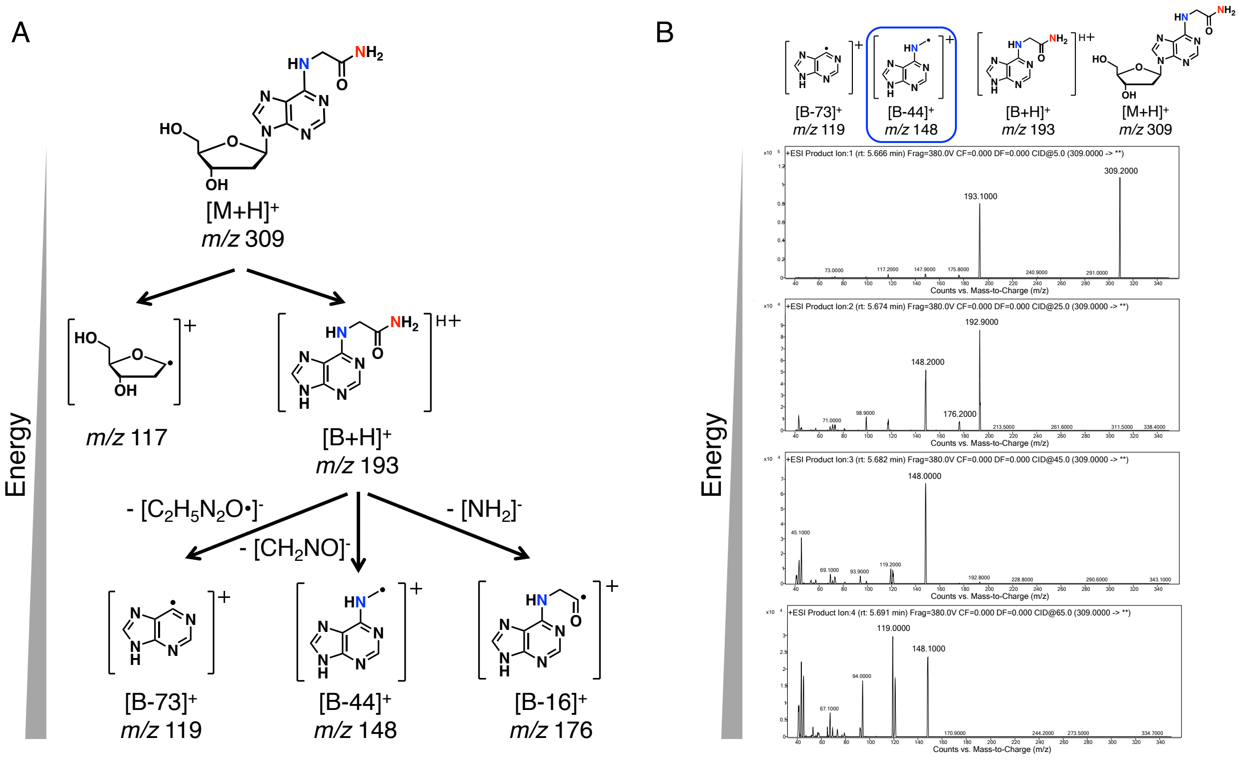

**Fig. S13. LC-MS/MS can be used to probe rearrangement step(s) required to form 6-*N*cmdA.** (**A**) MS/MS fragmentation pattern of 6-*N*cmdA observed by Karambelkar *et al*. The structure of 6-*N*cmdA was first reported by Swinton *et al.* in 1983 which is further supported by the MS/MS results of Karambelkar *et al.* (*3*, *38*). The daughter ion, $[B\text{-}44]^+$ (*m/z* 148), is produced by fragmentation between the exocylic C-C bond and is particularly diagnostic here. In this work, we also refer to this daughter ion as $[6mA^\bullet]^+$. (**B**) The fragmentation pattern of 6-*N*cmdA observed in this work. We see a fragmentation pattern for 6-*N*cmdA (nucleoside originating from the biotinylated DNA substrate) wholly consistent with the published fragmentation pattern. The mass spectra of all CID energies (5, 25, 45, and 65 V) are shown here. In Fig. 2, spectra from the 5 V fragmentation energy are shown for 6-*N*cmdA, and spectra from the 45 V fragmentation energy are shown for $[6mdA^\bullet]^+$.

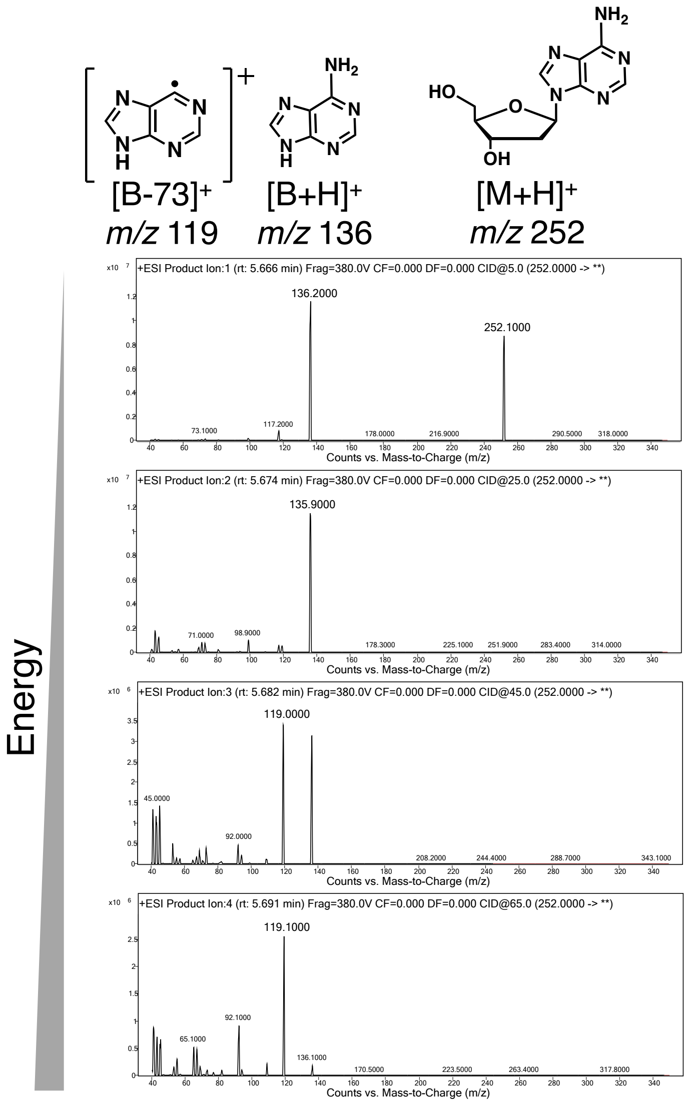

**Fig. S14. dA and 6-*N*cmdA can be separated at baseline resolution.**

The fragmentation pattern of dA shows that 6-*N*cmdA (or any daughter ions) cannot be detected by MS-MS, indicating that dA and 6-*N*cmdA can be fully separated with our UHPLC method. With our MS/MS methodology, we see the parent ion dA (*m/z* = 252) fragment into A (*m/z* = 136) and the purine ring (*m/z* = 119) (*49*, *50*).

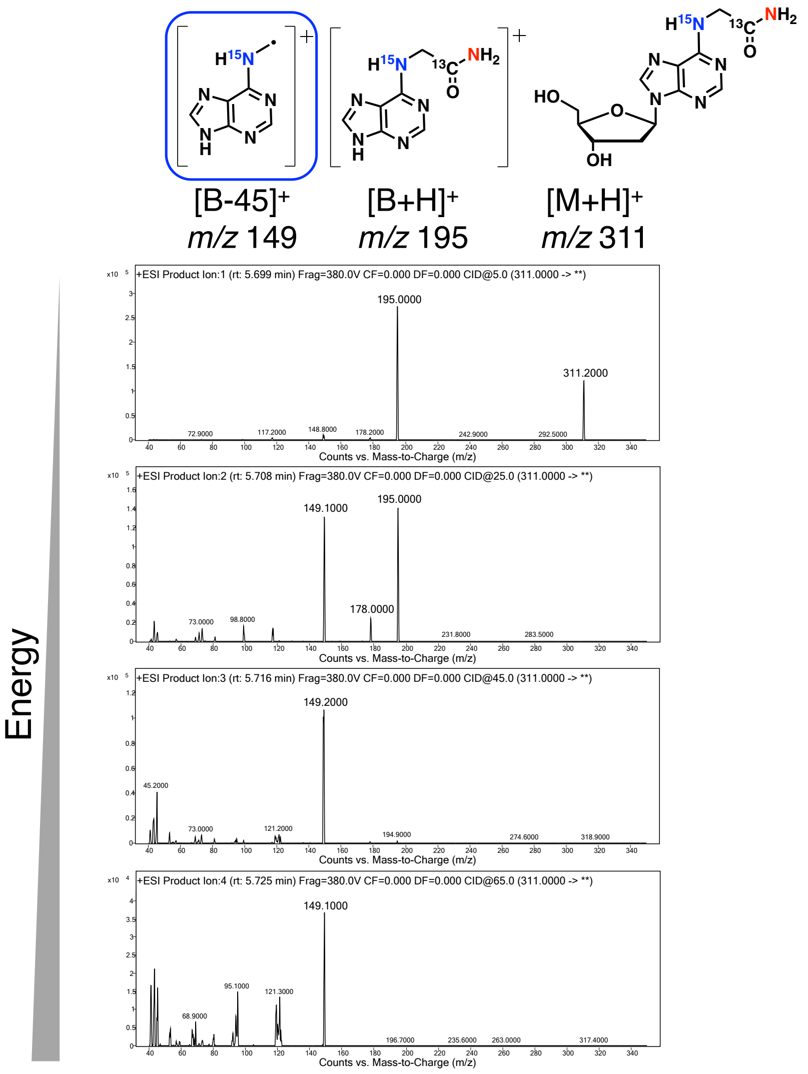

**Fig. S15. Fragmentation of 6-*N*cmdA labeled with $^{15}$N,$^{13}$C-glycine ($H_3{}^{15}NCH_2{}^{13}CO_2H$) demonstrates that the glycyl group 'flips' 180° during momylation, exchanging the nitrogen from the primary amine originating from glycine and the nitrogen originating from the *N*6 position of dA.**

Establishing that Gly-tRNA$^{Gly}$ is the co-substrate used by Mom to hypermodify dA allowed us next to investigate the discrepancy between Mom definitively transferring a glycyl group from a charged tRNA$^{Gly}$ donor and the structural isomer of the glycyl group that forms to yield the mature modification. To do so, we included a stable isotope of glycine, labeled at the α-amine with $^{15}$N (in blue) and at the carboxyl group with $^{13}$C, in a coupled Mom activity assay and used MS/MS to probe the positions of atoms within the modified nucleoside, 6-*N*cmdA, relative to the substrate, dA. All other reaction components were unlabeled. From the fragmentation pattern for 6-*N*cmdA (nucleoside originating from the biotinylated DNA substrate), we assign the major peaks seen to the parent nucleoside 6-*N*cmdA (*m/z* = 311, Δ *m/z* = +2 relative to unlabeled), the nucleobase 6-*N*cmA (*m/z* = 195, Δ *m/z* = +2 relative to unlabeled), and [6m-$^{15}$N-A$^{\bullet}$]$^{+}$ (*m/z* = 149, Δ *m/z* = +1 relative to unlabeled). We assign a smaller peak at seen in the 25 V spectrum at *m/z* 178 ($\Delta$ *m/z* = +2 relative to unlabeled) to the daughter ion fragmented between the C-N exocyclic amide bond. The key observation supporting a "flip" of the glycl group comes from the daughter ion, [6m-$^{15}$N-A$^{•}$]$^{+}$, where $^{15}$N-Labeled $\alpha$-amine nitrogen (in blue) has exchanged with the *N*6 nitrogen from dA (in red), accounting for an increase in mass of 1 Dalton. See main text and complementary labeling experiments with $^{15}N_5$-dA (below) for discussion of a possible mechanism that proceeds through a five-membered ring intermediate.

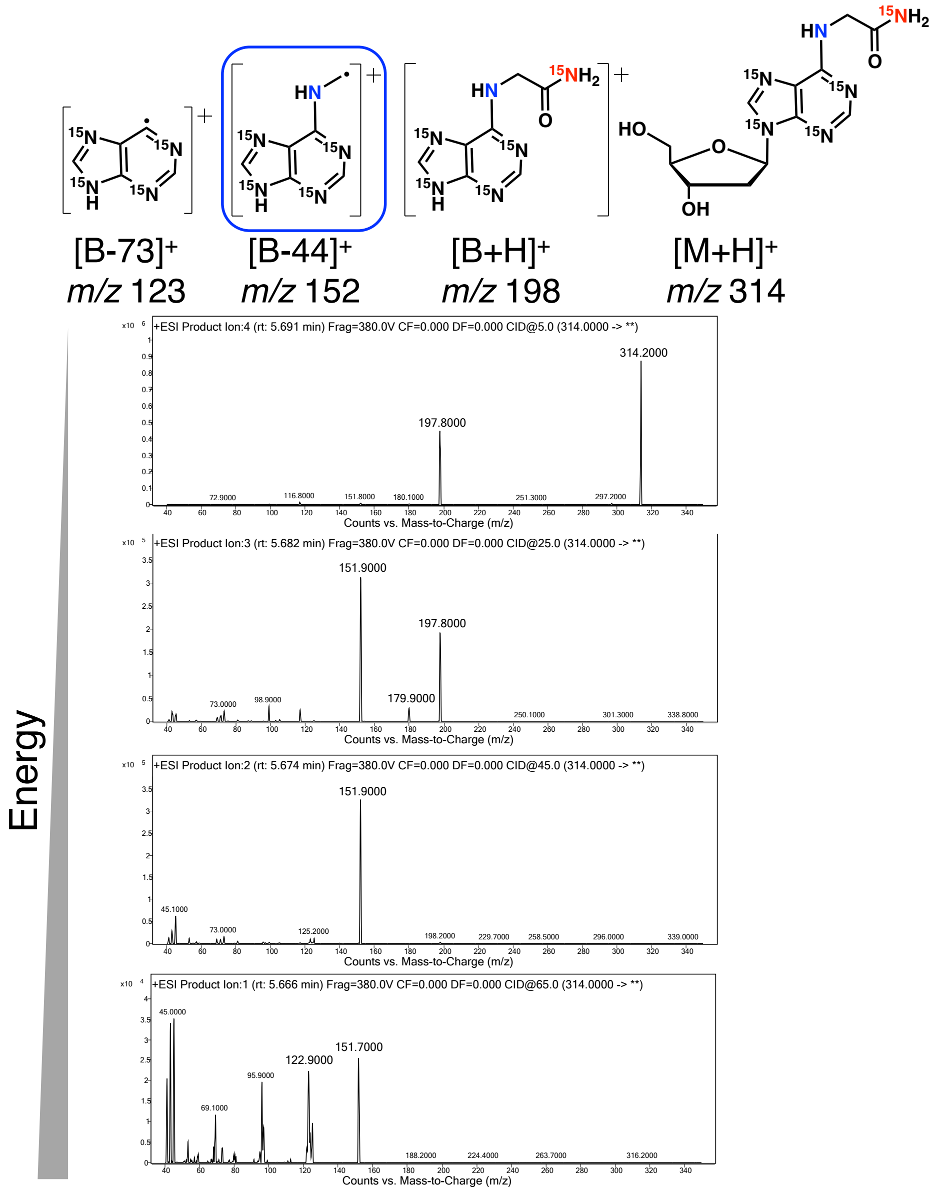

**Fig. S16. Fragmentation of 6-*N*cmdA produced by labeling with $^{15}$N-dA demonstrates that rearrangement of the glycyl group occurs after transfer to dA during momylation.**

After establishing that the glycyl group undergoes a rearrangement to a carbamoylmethyl group during momylation, we sought to gain insight on when this rearrangement occurs in the reaction sequence of momylation. We again utilized stable isotopes in a coupled, Mom activity assay, which here included a labeled dsDNA substrate generated by PCR with $^{15}N_5$-dATP ($^{15}$N at all 5 nitrogens, Δ *m/z* = +5), dC, dG, and dT. All other reaction components were unlabeled. Negative controls excluded Mom from the reaction (not shown). We assign the major peaks seen in the fragmentation pattern to the parent nucleoside 6-$^{15}$*N*cm-$^{15}N_4$-dA (*m/z* = 314, Δ *m/z* = +5 relative to unlabeled), the nucleobase 6-$^{15}$*N*cm-$^{15}N_4$-A (*m/z* = 198, Δ *m/z* = +5 relative to unlabeled), and [6m-$^{15}N_4$-A$^{\bullet}$]$^{+}$ (*m/z* = 152, Δ *m/z* = +4 relative to unlabeled). We assign a smaller peak at seen in the 25

V spectrum at *m/z* 180 (Δ *m/z* = +4 relative to unlabeled) to the daughter ion fragmented between the C-N exocyclic amide bond. For 6-*N*cmdA installed by Mom on the labeled $^{15}$N-dA substrate, we see the unlabeled nitrogen (in blue) originates from the α-amine of the glycyl group at the *N*6 position, as evidenced by both the daughter ions at *m/z* = 152 ([6m-$^{15}N_4$-A$^{\bullet}$]$^+$) and *m/z* 180 where Δ *m/z* = +4. The labeled nitrogen originating from dA at the *N*6 position is shown in red. Taken together, the mechanistic data for the momylation reaction are consistent with glycyl transfer from the tRNA Gly-tRNA$^{Gly}$ ester to the *N*6 atom the adenine target in the first step, and an intramolecular rearrangement reaction in the second step (also see Fig. 2). In the first step, a direct transfer of the glycyl group to *N*6 would require that the aromatic and hence relatively non-nucleophilic amine of the adenine to act as a nucleophile in the acylation reaction, with an alcohol (from tRNA) as the leaving group. It is possible that the enzyme active site can catalyze this otherwise difficult reaction, though, we are not aware of any precedent for GNAT family enzymes catalyzing acyl transfer to aromatic amines. Alternatively, there is also the possibility that the more nucleophilic nitrogen at *N*1 of adenine (*51*, *52*) carries out the initial nucleophilic attack, followed by a transfer of the acyl group from *N*1 to *N*6. Such transfer for alkyl groups has been observed on adenine (*53*) and is known as a Dimroth rearrangement (*54*). To our knowledge, Dimroth rearrangements do not have precedence in a biological context, and it is unclear whether an acyl group instead of an alkyl group could be transferred. Our data leave open whether the substrate base adenine carries out the nucleophilic attack directly, or whether there are intermediate transesterification steps on Mom, as has been reported for some GNATS (*55*). The latter would relax the requirement for spatial proximity between the tRNA acceptor site and the binding site for the substrate adenine. In the second step of the momylation reaction, the *N*6-acylation product rearranges to the final carbamoylmethyl modification. This step is strongly supported by the key observation from isotope labeling that the *N*6 of adenine becomes the distal amide nitrogen in the carbamoylmethyl group. We also note that the final orientation of carbamoylmethyl group remains to be determined. We expect that the exocyclic atoms point away from the Watson-Crick edge so as not to disrupt base pairing (*56*).

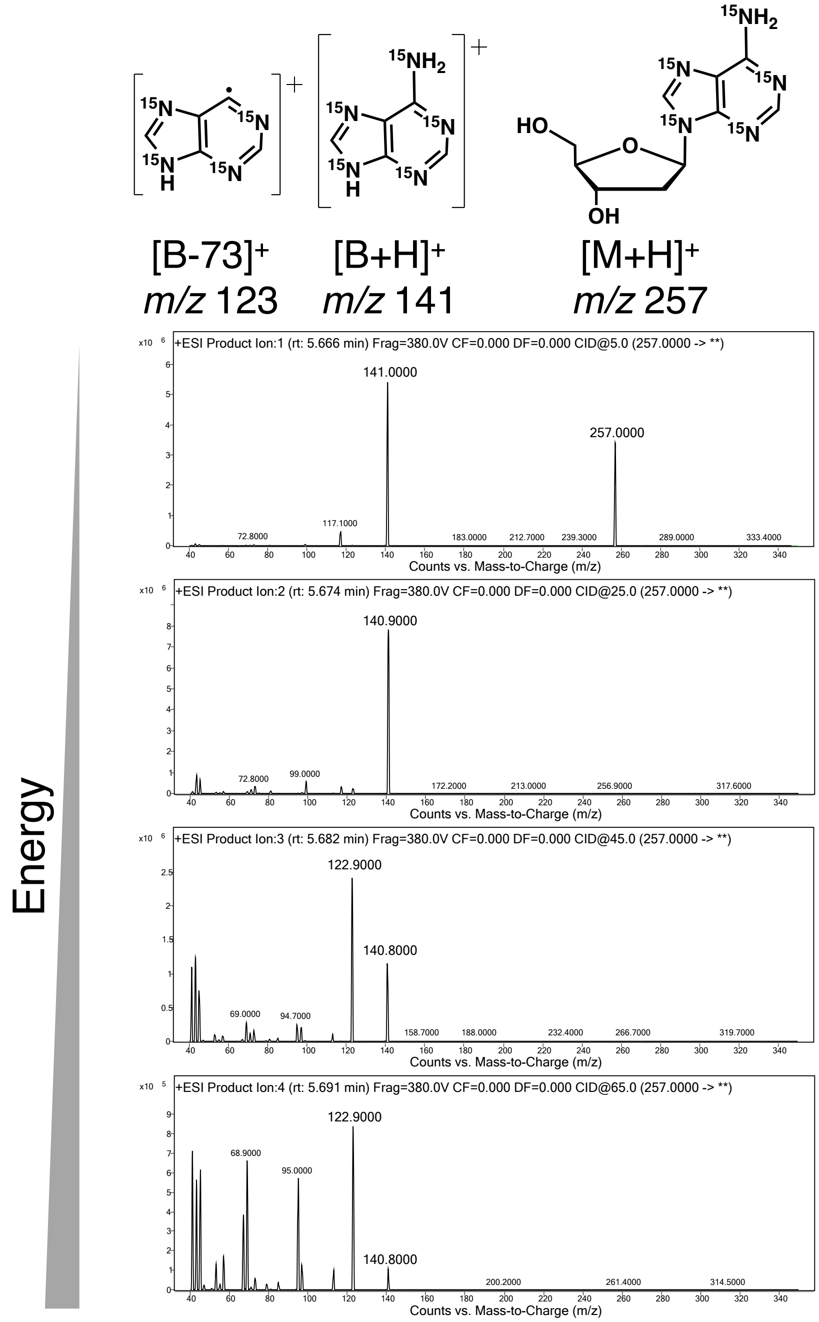

**Fig. S17. DNA can be labeled isotopically by PCR using $^{15}N_5$-dATP.**

A 3 kbp PCR product was amplified from a pUC19 template with $^{15}N_5$-dATP ($^{15}N$ at all 5 nitrogens, $\Delta$ *m/z* = +5), dC, dG, and dT. The labeled nucleoside (*m/z* = 257) and daughter ions can be seen.

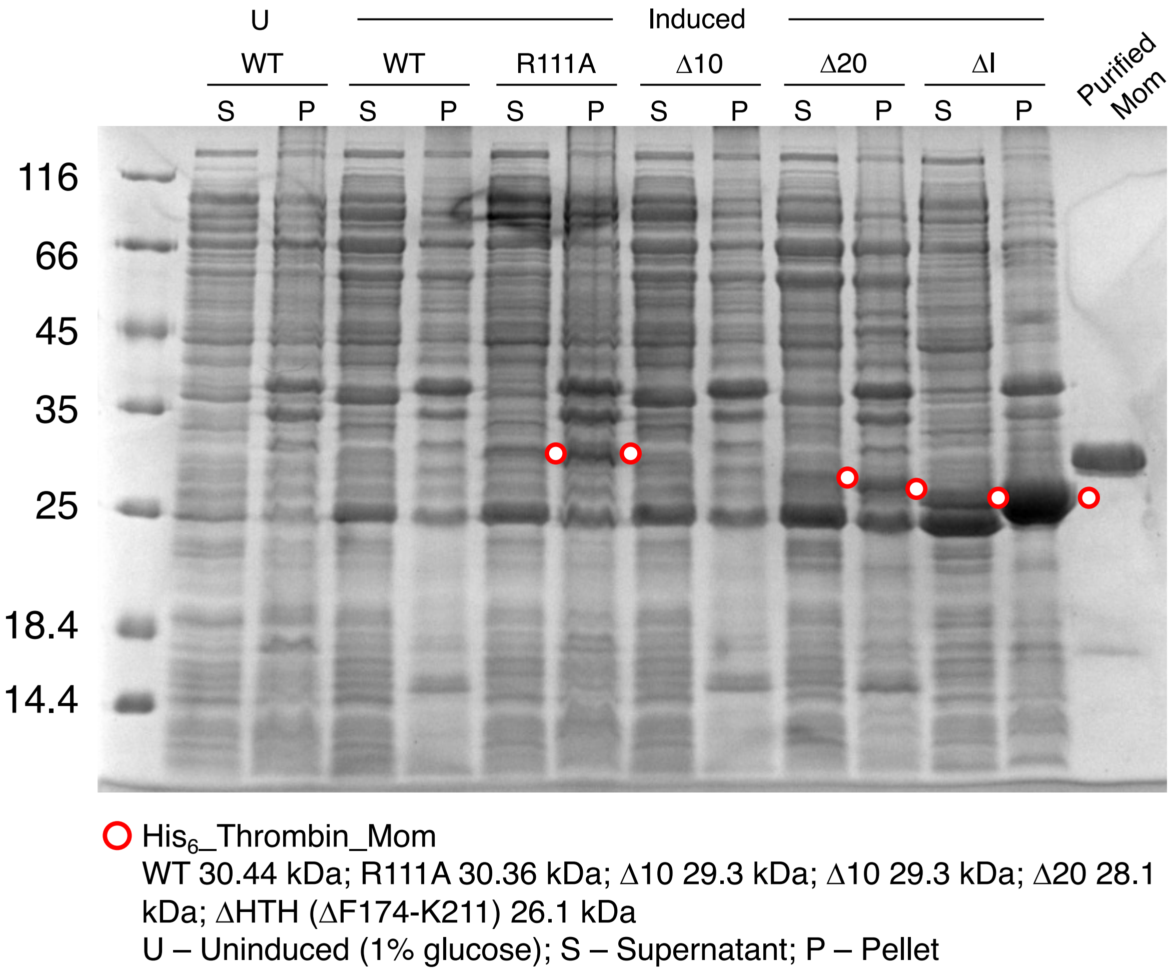

**Fig. S18. SDS-PAGE analysis of the expression of Mom variants.**

Expression of Mom variants with an N6-His-Thrombin_site was induced by IPTG overnight. Sample loading was normalized by the volume of culture in order to reflect the influence of Mom variants on culture growth. The gel was stained by Coomassie blue. While the expression of WT and Δ10 Mom is undetectable by Coomassie staining, the catalytically inactive (R111A, ΔI) and impaired (Δ20) forms is visible, especially in case of ΔI. The levels of Mom variants expression appear anticorrelated with their activity (see Fig. S19).

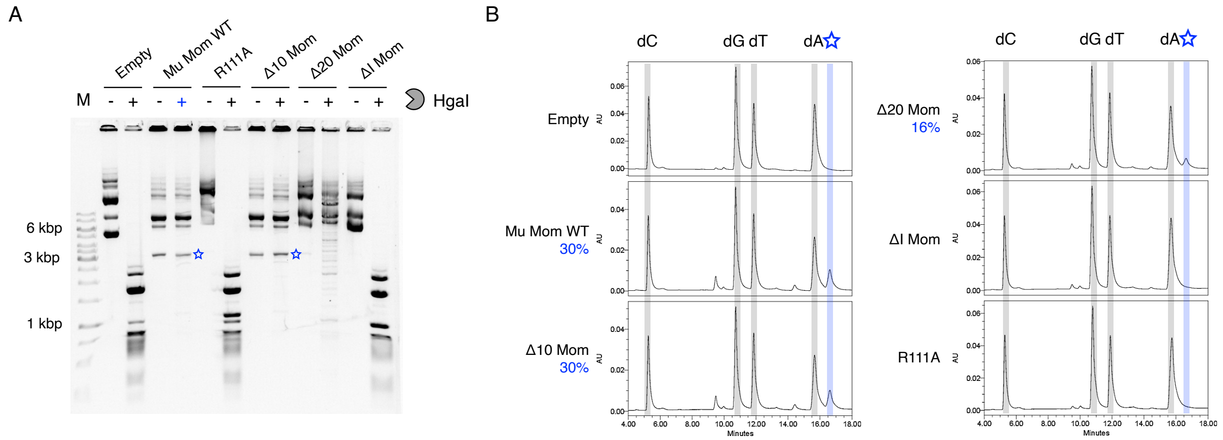

**Fig. S19.** ***In vivo*** **activity of Mom variants.**

(**A**) Restriction digest challenge of plasmids isolated from cultures expressing WT Mom and variants with N6-His-Thrombin tags. While WT and Δ10 Mom fully protect DNA from HgaI, the activity of Δ20 Mom is compromised, as reflected by partial plasmid digestion. In case of R111A and ΔI variants, the plasmids are completely digested, pointing to the lack of any activity. "Empty" lanes from samples expressing a pET28a plasmid that did not contain a Mom ORF were used as a negative control. (**B**) UHPLC analysis of the above plasmids. In agreement with panel A, the conversion rate for WT and Δ10 Mom is similar (~30%) while the conversion rate for Δ20 Mom is reduced (16%). R111A and ΔI Mom variants do not have detectable activity.

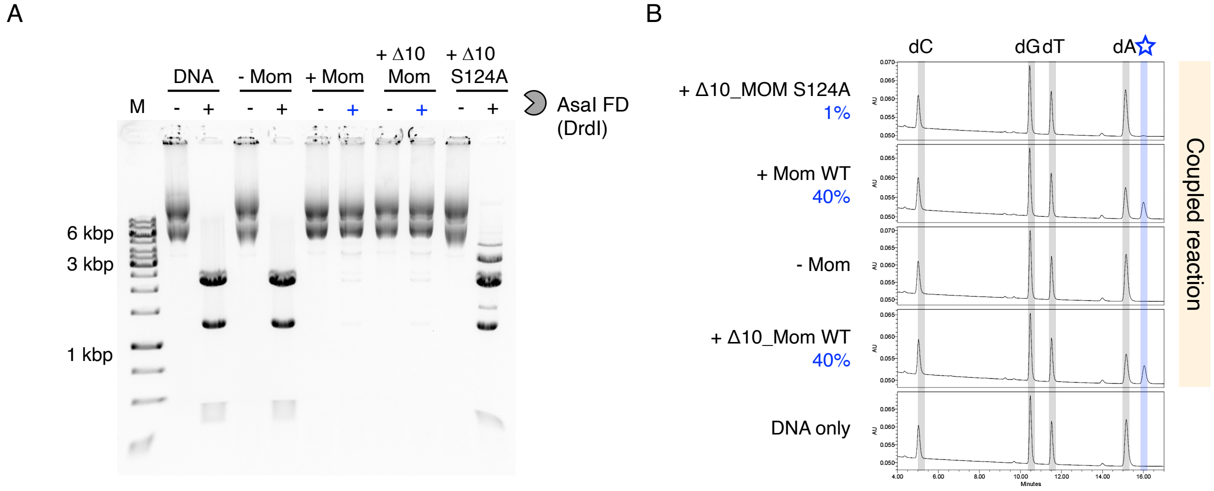

**Fig. S20. *In vitro* activity of Mom variants.**

(**A**) Restriction digest challenge of plasmid substrates following a coupled Mom activity assay. pET28a empty vectors were used as substrates in activity assays *in vitro*, which included purified Mom enzyme and components to generate Gly-tRNA$^{Gly}$ *in situ* – tRNA$^{Gly}$, glycine, ATP, and purified glycyl-tRNA synthetase (GlyRS) from *E. coli*. Reactions including WT Mom and Δ10 Mom were fully protected from restriction by AasI (DrdI) FastDigest (Thermo). In contrast, Δ10 S124A Mom was digested to near completion (compare to the digestion pattern in lanes under the DNA category). (**B**) UHPLC analysis of nucleosides derived from plasmid substrates. Consistent with our gel-based assay, by UHPLC we see high conversion of dA to 6-*N*cmdA in reactions including WT Mom (40%) and Δ10 Mom (40%), whereas a much lower rate of conversion is seen for reactions including Δ10 S124A Mom (1%).

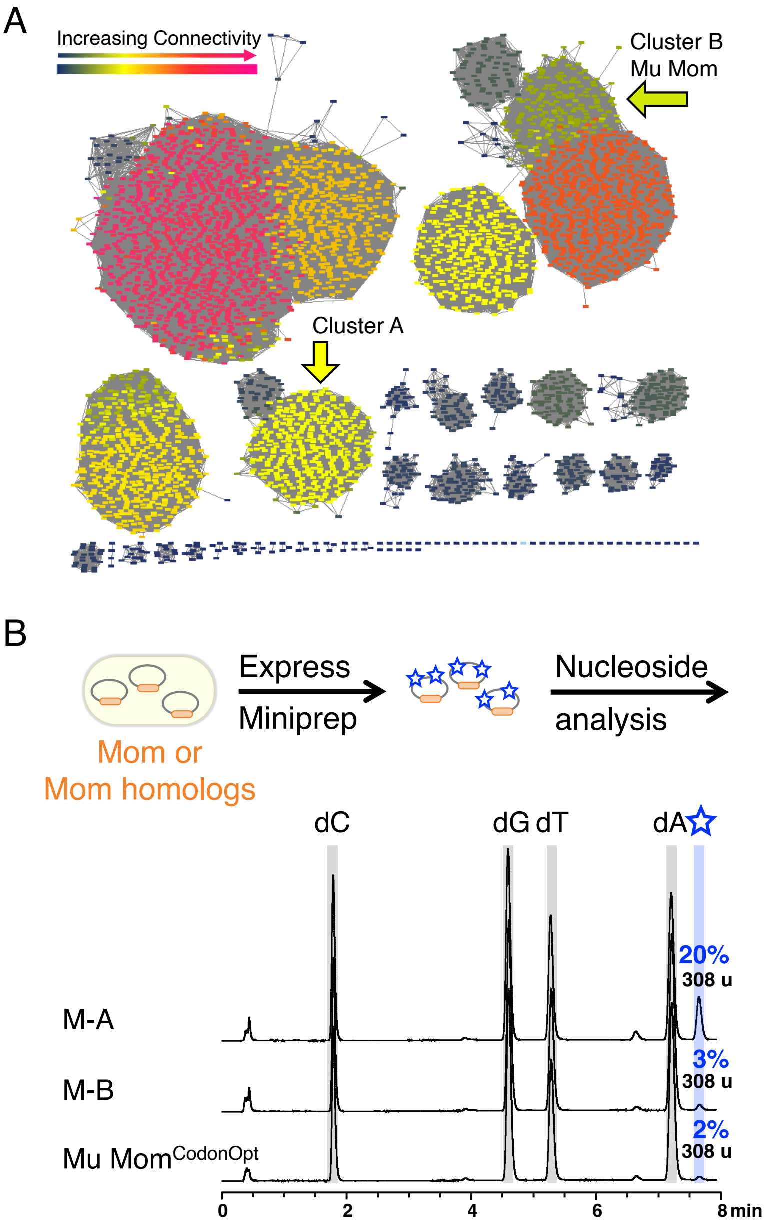

**Fig. S21. Recombinant expression of two predicted Mu Mom homologs leads to appearance of a novel nucleoside with the same retention time and mass as 6-*N*cmdA.**

(**A**) Sequence similarity network of Mu Mom and predicted homologs. After identifying the co-substrate of Mom and reconstituting 6-*N*cmdA *in vitro*, we next sought to recover viral sequences encoding homologs of Mom from metagenome databases in order to document the occurrence of Mom genes as well as any co-association with other genes potentially involved in the biosynthesis of as yet uncharacterized hypermodifications (*4*, *5*). See Methods as well. We sorted recovered sequences using a sequence similarity network (SSN). The network shown is a 95% representative node network (3852 nodes) containing 9723 sequences with 598234 edges representing an E-value cutoff of $1\times10^{-35}$ (*36*). Sequences from major clusters were selected at random for screening and ordered from GenScript with sequences codon optimized for expression in *E. coli*. One predicted homolog screened from Cluster A, Mom homolog A, comes from the cluster indicated. This sequence was found in viral samples collected from the Pacific Ocean. A second predicted homolog from Cluster B, Mom Homolog B, comes from the same cluster as Mu Mom. Homolog B was found in human waste samples. The database entry identifiers are given in table S2. (**B**) Two functional homologs of Mu Mom discovered. Predicted homologs of Mu Mom were screened using our *in vivo* activity assay, and the activity of selected homologs are shown. UHPLC-MS data for Homolog A comes from a 50 mL culture (shaken at 225 rpm in a 250 mL flask) where expression was induced with 100 µM IPTG. Cells were incubated for 18 hours at 18 °C. UHPLC-MS data for Homolog B and Mu $\mathrm{Mom}^{\mathrm{CodonOpt}}$ comes from a 5 mL culture (also shaken at 225 rpm in a 29 mL glass tube) using the same expression conditions. Recombinant expression of both Homolog A and Homolog B results in a novel peak with the same retention time (~7.8 min) and mass (308 u) as 6-*N*cmdA. We conclude that the novel peak is 6-*N*cmdA. The reduced extent of hypermodification seen for Mu $\mathrm{Mom}^{\mathrm{CodonOpt}}$ may be attributed to different growth conditions. The reduced extent of hypermodification seen for Homolog B remains to be determined. As with Mu $\mathrm{Mom}^{\mathrm{CodonOpt}}$, the growth conditions may not have been optimal, or there may be different sequences specificities to explore. The active Mom homologs found suggest that coopting activated glycine from the host translation machinery is used as an effective strategy for genome defense beyond phage Mu. As translation is universal and tRNAs are well conserved, the phage can expect to find an abundant source of tRNA-activated glycine in the broad range of bacteria that phage can infect. Potential damage to the host translation system from an imbalance of charged tRNAs is not a concern for Mu, since Mom is expressed late in the phage infection cycle just prior to lysis (*3*, *38*, *57–60*). The infection strategies of other phages may be similar.

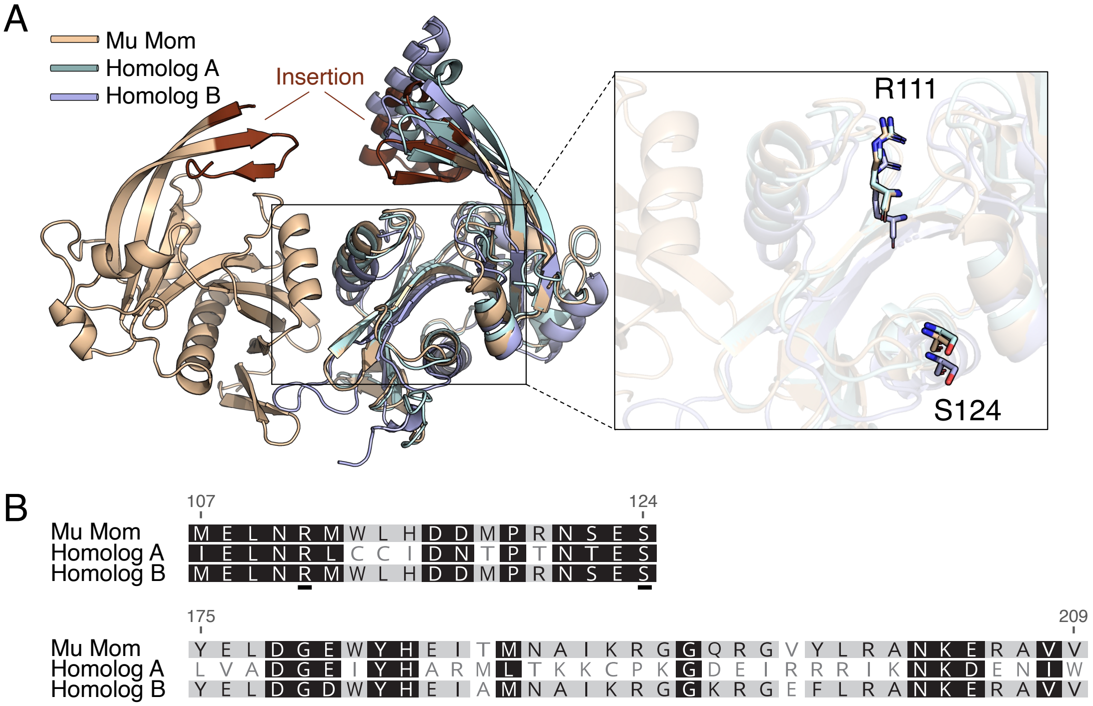

**Fig. S22. Structure and sequence comparison of Mu Mom with active homologs provides insights to regions important for function.**

(**A**) Superposition of Mu Mom and active homologs. The structures of M-A and M-B were predicted using ColabFold (*61*) and superimposed with Chain A of Mom (light brown) using PyMOL (*62*). Mom Homolog A (teal) aligned with Mu Mom at an rmsd value of 4.567, and Mom Homolog B (lavender) aligned with Mu Mom at an rmsd value of 0.672. Predicted structures of Homolog A and Homolog B suggest that the Insertion sequence, shown in brown for Mu Mom, is also present in these active homologs. The inset shows the region including R111 and S124 (alanine is in position 124 for Mu Mom, see methods for details about crystallization of Mom Δ10 S124A). These residues were shown in this work and Karambelkar *et al.* to be critical for activity (*3*). Y159, H58, and D149 were also found by Karambelkar *et al.* to be important for activity (residues not shown here). The corresponding arginine and serine residues for Homologs A and B are also shown. The activity data from this work, structural alignments of Mu Mom with other GNATs, and the alignments shown above all point to this region containing the active site of Mom. It remains to be explored how these residues are specifically involved in transaminoacylation and/or the rearrangement (*39*, *41*, *55*). (**B**) Sequences alignments of Mu Mom and active homologs. Residues in proximity of the putative active side, 107-124 (Mu Mom numbering), are well-conserved among Mu Mom, Homolog A, and Homolog B. The insertion region, residues 175-209, in contrast are not as well-conserved, but are predicted to adopt a similar structure. A hallmark of the GNAT superfamily is strong structural conservation, but poor sequence conservation. It may be the case that for Mom-like GNATs, this region is variable and confers co-substrate and/or substrate specificities.

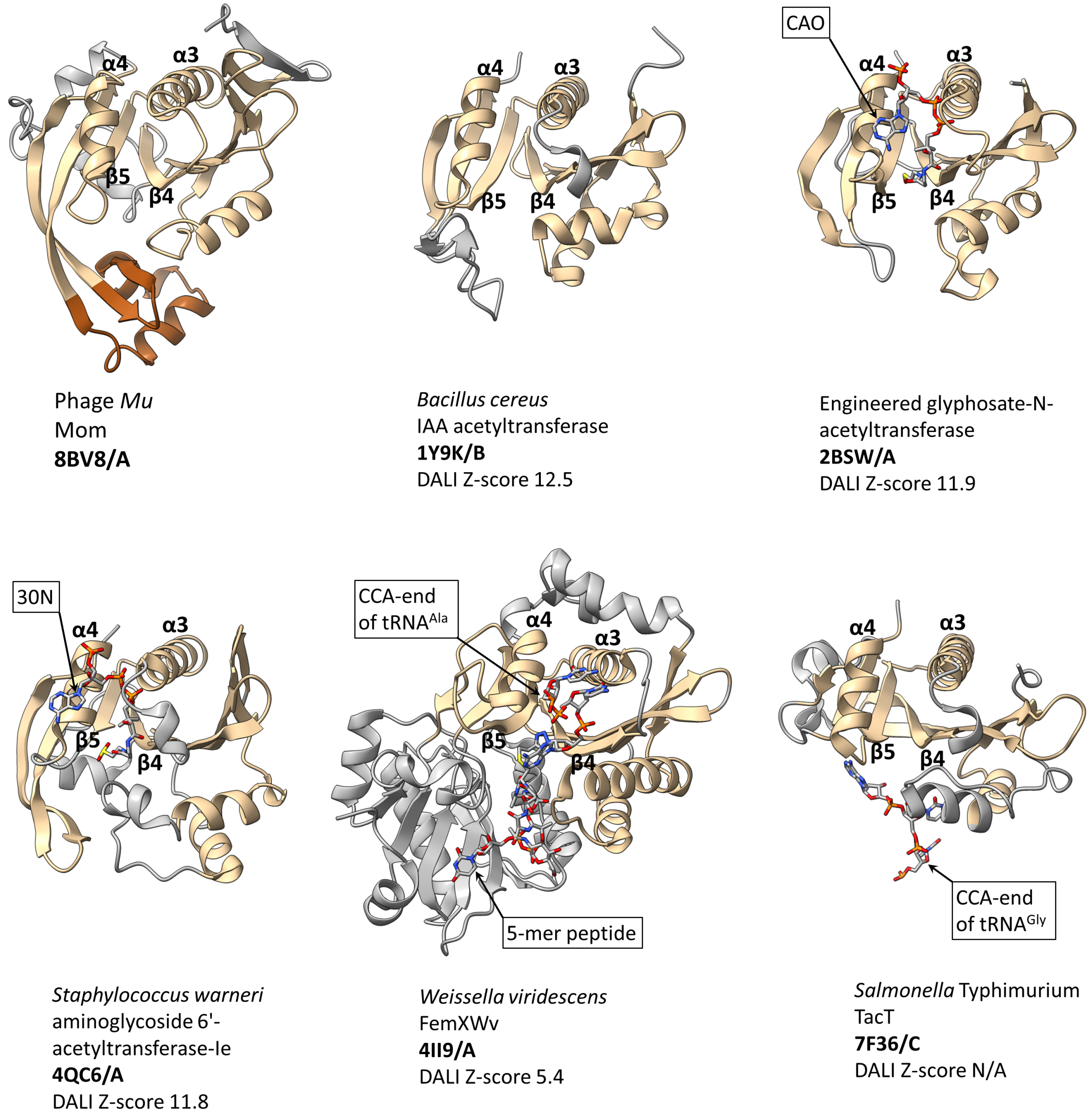

**Fig. S23. Structural comparison of Mom with members of the GNAT family.**

1Y9K, 2BSW, and 4QC6 are the top three hits from the structural similarity search performed against the chain A of Mom (displayed) using the DALI server (*63*). Structures of FemXWv (4II9) and TacT (7F36) are added due to their relevance to our study, despite a low (FemXWv) or unavailable (TacT) Z-score. All six presented structures feature a conserved GNAT core colored in pale brown. The characteristic splay between β4 and β5 strands which framed by α3 and α4 helices is highlighted. Less conserved or non-GNAT regions colored in grey. The Mom-specific insertion is colored in brown. Ligands are represented as sticks and colored by element. See Methods for a list of PDB ID and chains of GNATs with bound CoA and acyl-CoA that were used for superposition in ChimeraX (*24*). CAO — oxidized coenzyme A. 30N — (3R,5S,9R)-1-[(2R,3S,4R,5R)-5-(6-amino-9H-purin-9-yl)-4-hydroxy-3-(phosphonooxy)tetrahydrofuran-2-yl]-3,5,9-trihydroxy-8,8-dimethyl-10,14-dioxo-2,4,6-trioxa-11,15-diaza-3,5-diphosphaheptadecane-17-sulfinic acid 3,5-dioxide.

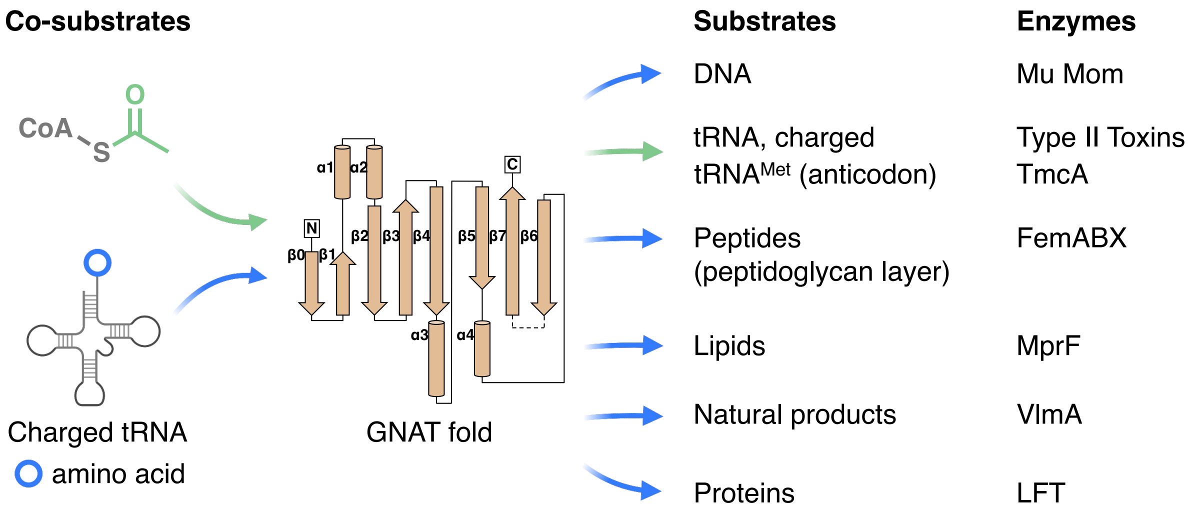

**Fig. S24. tRNA as a co-substrate or substrate of GNAT enzymes.**

Noncanonical roles for aminoacylated tRNA outside of ribosome-dependent protein synthesis are found in all domains of life and continue to emerge in diverse cellular processes (*44*, *45*, *64–67*). Analogous to its canonical role in ribosome-dependent protein synthesis, the aminoacyl group at the 3′ end of a charged tRNA is utilized as an amino acid donor to modify diverse target substrates. Known for its remarkable diversity of co-substrates, substrates, and functions, the GNAT superfamily uses tRNA for several noncanonical purposes (*39–41*, *68–72*). In the case of N-linked aminoacylation by GNAT enzymes that use charged tRNA as the co-substrate, a new amide bond is produced Highlighted here are selected examples from microbes where tRNA has been found either as the co-substrate used to aminoacylate target substrates (orange arrows) or as the substrate that is the target of modification (black arrow). Many instances of charged tRNA acting as an aminoacyl donor have been reported. At the time of this publication, tRNA is known to be the substrate for GNAT enzymes in a few systems. The α-amine at the charged 3′ end can be acetylated, as for Type II Toxins (ex. AtaT, TacT, etc.), or the wobble base in the anticodon region can be acetylated by TmcA as part of maturation for $tRNA^{Met}$. These examples are not comprehensive, but instead emphasize the broad range of cellular processes where GNATs and tRNA are involved. The factors that govern co-substrate and substrate specificity for this superfamily are not fully understood. For example, the co-substrate and substrate of Mom could not be predicted *de novo* from the structure alone or through alignments of Mom with the most closely related structural homologs found by the DALI server. Continued advances in computational approaches may give insights on what features have been repurposed or acquired over protein evolution (insertions, dimerization modes, etc.) and are driving function. Finally, analogous to other instances where the GNAT has been repurposed, we speculate that perhaps the category of GNAT enzymes defined by Mu Mom represents an evolutionary steppingstone where the GNAT fold has evolved to accommodate both a co-substrate and target that are nucleic acids (*72*).

**Table S1. Modification motifs in phage Mu identified by SMRT sequencing**

| Motif | Position | Type[a] | % of Motifs Detected | # of Motifs Detected | # of Motifs in Genome | Mean QV | Mean Coverage | Mean IPD Ratio | Objective Score |
|---|---|---|---|---|---|---|---|---|---|
| Phage Mu, *mom*$^{+}$ | | | | | | | | | |
| SAS | 2 | m6A[b] | 70.10 | 3,339 | 4760 | 355.4 | 424.1 | 3.9 | 862,376.90 |
| GATC | 2 | m6A[c] | 94.00 | 143 | 152 | 401.9 | 422.4 | 3.8 | 54,405.80 |
| HSGAAB | 4 | m6A | 63.70 | 197 | 309 | 243.8 | 423.4 | 3.4 | 32,026.30 |
| Phage Mu, *mom*$^{-}$ | | | | | | | | | |
| GATC | 2 | m6A | 96.00 | 144 | 150 | 560 | 460.7 | 5.1 | 77,726.30 |
| GCA(N)$_6$GTT | 3 | m6A | 100.00 | 6 | 6 | 427.2 | 504.3 | 3.7 | 2,563.00 |
| AAC(N)$_6$GTGC | 2 | m6A | 100.00 | 6 | 6 | 415.7 | 466.2 | 3.9 | 2,494.00 |

a. The modification type detected was reported to be m6A in all sequence contexts. The analysis methods used were trained on the m6A modification, so modifications with a similar deviation of IPD patterns will be assigned as m6A. b. For the *mom*$^{+}$ strain, the modification for the detected SAS motif is in fact 6-*N*cmdA as shown experimentally. c. The GATC motif is detected because host *E. coli* strains are all *dam*$^{+}$.

### Table S2. Primers, Plasmids, Co-substrates, and Substrates

| **Primers** | **Name** | **Sequence** | | | | | | |
|---|---|---|---|---|---|---|---|---|
| Primers (Cloning) | YJLo260 | AATTAAGAAGGAGATATACAATGCCTGCGAGCATCCCA | | | | | | |
| | YJLo261 | TGGTGGTGGTGCTCGAGTGCTCACTTAGGGTATGGCTGAACC | | | | | | |
| | YJLo263 | TGGTGGTGGTGCTCGAGTGCCTTAGGGTATGGCTGAACCTTGAATAG | | | | | | |
| Primers (Variants) | Δ10 truncation F | ctggtgccgcgcggcagccaTGTTGGTAAAGAGAAAAAAAGCCGCATCC | | | | | | |
| | Δ10 truncation R | GGATGCGGCTTTTTTTCTCTTTACCAACAtggctgccgcgcggcaccag | | | | | | |
| | Δ20 truncation F | ctggtgccgcgcggcagccaTACCAAACCGTGTGTGATTGAATATGAAG | | | | | | |
| | Δ20 truncation R | CTTCATATTCAATCACACACGGTTTGGTAtggctgccgcgcggcaccag | | | | | | |
| | ΔI truncation F | CGGCAGCCATGAAAGCACCggcagcTTCAACCAGTATCGTTATATCCGCTTCC | | | | | | |
| | ΔI truncation R | GGAAGCGGATATAACGATACTGGTTGAAgctgccGGTGCTTTCATGGCTGCCG | | | | | | |
| | R111A F | GTTATATGGAACTGAATgcTATGTGGCTGCATGATGA | | | | | | |
| | R111A R | TCATCATGCAGCCACATAgcATTCAGTTCCATATAAC | | | | | | |
| | S124A F | GCCTCGTAATAGCGAAgcCCGTGCAATTAGCTATG | | | | | | |
| | S124A R | CATAGCTAATTGCACGGgcTTCGCTATTACGAGGC | | | | | | |
| Primers (PCR) | RS002 | TGGAGCGGGAAACGAGAC | | | | | | |
| | RS003 | CCATTCAGGCTGCGCAAC | | | | | | |
| | RS005 | CGACTCACTATAGCGGGAATAG | | | | | | |
| | RS006 | TATTACTGCAGCAATTCACTG | | | | | | |
| | RS047 | TGGAGCGGGCAGCGGGAA | | | | | | |
| | RS049 | TGGAGCGGGCGAAGGGAA | | | | | | |
| **Plasmids** | **Vector Name** | **Resistance** | **Gene** | **Tag** | **Cleavage Sites** | **N or S?** | **Database** | **Identifier** |
| | pET 28a WT Mom | Kan | *mom* | N-$His_6$ | Thrombin | S | Uniprot | P06018 |
| | pET 28a R111A Mom | Kan | *mom* | N-$His_6$ | Thrombin | S | Uniprot | P06018 |
| | pET 28a Δ10 Mom | Kan | *mom* | N-$His_6$ | Thrombin | S | Uniprot | P06018 |
| | pET 28a Δ20 Mom | Kan | *mom* | N-$His_6$ | Thrombin | S | Uniprot | P06018 |
| | pET 28a ΔI Mom | Kan | *mom* | N-$His_6$ | Thrombin | S | Uniprot | P06018 |
| | pET 28a Δ10 R111A Mom | Kan | *mom* | N-$His_6$ | SUMO | S | Uniprot | P06018 |
| | pET 28a Δ10 S124A Mom | Kan | *mom* | N-$His_6$ | SUMO | S | Uniprot | P06018 |
| | pRY Mu Mom | Kan | *mom* | $His_6$-C | NA | N | Uniprot | P06018 |
| | pET 28a WT Mom | Kan | *mom* | N-$His_6$ | NA | S | Uniprot | P06018 |
| | pET 28a Mom Homolog A | Kan | *momologA* | N-$His_6$ | NA | S | IMG/VR | See Caption[a] |
| | pET 28a Mom Homolog B | Kan | *momologB* | N-$His_6$ | NA | S | IMG/VR | See Caption[b] |
| | pET 21a GlyRS | Amp | *glyQ, glyS* | $His_6$-C | NA | N | Uniprot; Addgene | P00960, P00961; 124110 |
| | pUC19 $tRNA^{Gly/GCC}$ | Amp | *glyV* | NA | NA | N | GtRNAdb | tRNA-Gly-GCC-1-1 |
| | pUC19 $tRNA^{Gly/CCC}$ | Amp | *glyU* | NA | NA | S | GtRNAdb | tRNA-Gly-CCC-1-1 |
| | pUC19 $tRNA^{Gly/TCC}$ | Amp | *glyT* | NA | NA | S | GtRNAdb | tRNA-Gly-TCC-1-1 |
| **Co-substrates** | **Name** | **Sequence** | | | | | | |
| | $tRNA^{Gly/GCC}$ | GCGGGAAUAGCUCAGUUGGUAGAGCACGACCUUGCCAAGGUCGGGGUCGCGAGUUCGAGUCUCGUU<br>UCCCGCUCCA | | | | | | |
| | $tRNA^{Gly/CCC}$ | GCGGGCGUAGUUCAAUGGUAGAACGAGAGCUUCCCAAGCUCUAUACGAGGGUUCGAUUCCCUUCGC<br>CCGCUCCA | | | | | | |
| | $tRNA^{Gly/TCC}$ | GCGGGCAUCGUAUAAUGGCUAUUACCUCAGCCUUCCAAGCUGAUGAUGCGGGUUCGAUUCCCGCUG<br>CCCGCUCCA | | | | | | |
| **Substrates** | **Name** | **Sequence (for oligomers)** | | | | | | |
| | 11-mer, top | CGCCGACGCGC | | | | | | |
| | 11-mer, bottom | GCGCGTCGGCG | | | | | | |
| | 11-mer, A/C MM bottom | GCGCGCCGGCG | | | | | | |
| | 11-mer, A/G MM bottom | GCGCGGCGGCG | | | | | | |
| | 11-mer, A/A MM bottom | GCGCGACGGCG | | | | | | |
| | 34-mer, top | GCAGTGTTCGACGCGGTAGTCTATCAATGCATGA | | | | | | |
| | 34-mer, bottom | TCATGCATTGATAGACTACCGCGTCGAACACTGC | | | | | | |
| | $WM_{Mu}$ top (RS025) | ACCGTTACCGCACTGGCTGCCGGTGAAGCAGGAA | | | | | | |
| | $WM_{Mu}$ bottom (RS026) | TTCCTGCTTCACCGGCAGCCAGTGCGGTAACGGT | | | | | | |
| | M13 ssDNA | from supplier | | | | | | |
| | Biotinylated dsDNA | derived from λ DNA | | | | | | |
| | 3 kbp PCR product | amplified from pUC19 plasmids | | | | | | |
| | pET28a empty plasmid | from supplier | | | | | | |

Primers, plasmids, co-substrates and substrates were stored in TE buffer, elution buffer (from miniprep kits), nuclease-free water (from the RNA Clean Up Kit), and IDT duplex buffer, respectively. All expression plasmids were under the control of a T7 promoter. N = native, S = synthetic. Mom Homolog A: IMGVR_UViG_3300002484_000500|JGI25129J35166_1001831|JGI25129J35166_10018319. Mom Homolog B: IMGVR_UViG_2872672955_000004|2872672955|2872672955|4871805-4913505_rc.

**Table S3. Data collection and refinement statistics**

| **Data collection statistics** | |
|---|---|
| Beamline | PETRA III P11 |
| Space group | P 21 21 21 |
| Cell dimensions<br>a, b, c (Å)<br>α, β, γ (°) | <br>57.18 66.48 138.39<br>90 90 90 |
| Wavelength (Å) | 1.0332 |
| Resolution range (Å)<br>Highest shell | 48 - 2.03<br>2.10 - 2.03 |
| Total reflections* | 465630 (47677) |
| Unique reflections* | 34815 (3412) |
| Completeness (%)* | 99.7 (99.5) |
| Multiplicity * | 13.4 (14.0) |
| Mean $I/\sigma I$ * | 14.6 (0.85) |
| R(merge) (%)* | 18.6 (315) |
| R(meas) (%)* | 19.4 (327) |
| R(pim) (%)* | 5.2 (87) |
| $CC_{1/2}$ (%)* | 99.9 (74.8) |
| Solvent content (%) | 58 |
| Wilson B-factor (Å$^2$) | 48 |
| **Refinement statistics** | |
| Protein atoms excluding H | 3549 |
| Protein residues | 424 |
| Solvent molecules | 190 |
| $R_{cryst}$ (%) | 21.2 |
| $R_{free}$ (%) $ | 24.0 |
| RMSD bond lengths (Å) | 0.002 |
| RMSD angles (°) | 0.44 |
| Ramachandran favored (%) | 98.3 |
| Ramachandran allowed (%) | 1.7 |
| Ramachandran outliers (%) | 0.00 |
| Rotamer outliers (%) | 1.6 |
| Clashscore | 2.3 |
| **PDB code** | **8BV8** |

* Highest-resolution shell are shown in parentheses

$ 5% of reflections (1741) were set aside randomly

### References

1. Y.-J. Lee, P. R. Weigele, Detection of Modified Bases in Bacteriophage Genomic DNA. *Methods Mol Biology Clifton N J*. 2198, 53–66 (2020).

2. D. Li, C.-M. Liu, R. Luo, K. Sadakane, T.-W. Lam, MEGAHIT: an ultra-fast single-node solution for large and complex metagenomics assembly via succinct de Bruijn graph. *Bioinformatics*. 31, 1674–1676 (2015).

3. S. Karambelkar, S. Udupa, V. N. Gowthami, S. G. Ramachandra, G. Swapna, V. Nagaraja, Emergence of a novel immune-evasion strategy from an ancestral protein fold in bacteriophage Mu. *Nucleic Acids Res*. 48, 5294–5305 (2020).

4. Y.-J. Lee, N. Dai, S. E. Walsh, S. Müller, M. E. Fraser, K. M. Kauffman, C. Guan, I. R. Corrêa, P. R. Weigele, Identification and biosynthesis of thymidine hypermodifications in the genomic DNA of widespread bacterial viruses. *P Natl Acad Sci Usa*. 115, E3116–E3125 (2018).

5. Y.-J. Lee, N. Dai, S. I. Müller, C. Guan, M. J. Parker, M. E. Fraser, S. E. Walsh, J. Sridar, A. Mulholland, K. Nayak, Z. Sun, Y.-C. Lin, D. G. Comb, K. Marks, R. Gonzalez, D. P. Dowling, V. Bandarian, L. Saleh, I. R. Corrêa, P. R. Weigele, Pathways of thymidine hypermodification. *Nucleic Acids Res*. 50, 3001–3017 (2021).

6. M. J. Cavaluzzi, P. N. Borer, Revised UV extinction coefficients for nucleoside-5′-monophosphates and unpaired DNA and RNA. *Nucleic Acids Res*. 32, e13–e13 (2004).

7. D. B. Dunn, R. H. Hall, Handbook of Biochemistry and Molecular Biology, 269–358 (2010).

8. C. J. L. Francois, Y. H. Jang, T. Cagin, W. A. Goddard, L. C. Sowers, Conformation and Proton Configuration of Pyrimidine Deoxynucleoside Oxidation Damage Products in Water. *Chem Res Toxicol*. 13, 462–470 (2000).

9. P. P. Chan, T. M. Lowe, GtRNAdb: a database of transfer RNA genes detected in genomic sequence. *Nucleic Acids Res*. 37, D93–D97 (2009).

10. P. P. Chan, T. M. Lowe, GtRNAdb 2.0: an expanded database of transfer RNA genes identified in complete and draft genomes. *Nucleic Acids Res*. 44, D184–D189 (2016).

11. Y. Shimizu, T. Ueda, Cell-Free Protein Production, Methods and Protocols. *Methods Mol Biology*. 607, 11–21 (2009).

12. C. Tuckey, H. Asahara, Y. Zhou, S. Chong, *Curr Protoc Mol Biology*, in press, doi:10.1002/0471142727.mb1631s108.

13. H. Asahara, O. C. Uhlenbeck, The tRNA Specificity of Thermus thermophilus EF-Tu. *Proc National Acad Sci*. 99, 3499–3504 (2002).

14. A. L. Edwards, A. D. Garst, R. T. Batey, Determining structures of RNA aptamers and riboswitches by X-ray crystallography. *Methods Mol Biology Clifton N J*. 535, 135–63 (2009).

15. R. Villet, M. Fonvielle, P. Busca, M. Chemama, A. P. Maillard, J.-E. Hugonnet, L. Dubost, A. Marie, N. Josseaume, S. Mesnage, C. Mayer, J.-M. Valéry, M. Ethève-Quelquejeu, M. Arthur, Idiosyncratic features in tRNAs participating in bacterial cell wall synthesis. *Nucleic Acids Res*. 35, 6870–6883 (2007).

16. E. P. Quinlivan, J. F. Gregory, DNA digestion to deoxyribonucleoside: A simplified one-step procedure. *Anal Biochem*. 373, 383–385 (2008).

17. S. E. Walker, K. Fredrick, Preparation and evaluation of acylated tRNAs. *Methods*. 44, 81–86 (2008).

18. H. Gamper, Y.-M. Hou, A Label-Free Assay for Aminoacylation of tRNA. *Genes-basel*. 11, 1173 (2020).

19. M. Krug, M. S. Weiss, U. Heinemann, U. Mueller, XDSAPP : a graphical user interface for the convenient processing of diffraction data using XDS. *J Appl Crystallogr*. 45, 568–572 (2012).

20. A. J. McCoy, R. W. Grosse-Kunstleve, P. D. Adams, M. D. Winn, L. C. Storoni, R. J. Read, Phaser crystallographic software. *J Appl Crystallogr*. 40, 658–674 (2007).

21. J. Jumper, R. Evans, A. Pritzel, T. Green, M. Figurnov, O. Ronneberger, K. Tunyasuvunakool, R. Bates, A. Žídek, A. Potapenko, A. Bridgland, C. Meyer, S. A. A. Kohl, A. J. Ballard, A. Cowie, B. Romera-Paredes, S. Nikolov, R. Jain, J. Adler, T. Back, S. Petersen, D. Reiman, E. Clancy, M. Zielinski, M. Steinegger, M. Pacholska, T. Berghammer, S. Bodenstein, D. Silver, O. Vinyals, A. W. Senior, K. Kavukcuoglu, P. Kohli, D. Hassabis, Highly accurate protein structure prediction with AlphaFold. *Nature*. 596, 583–589 (2021).

22. D. Liebschner, P. V. Afonine, M. L. Baker, G. Bunkóczi, V. B. Chen, T. I. Croll, B. Hintze, L.-W. Hung, S. Jain, A. J. McCoy, N. W. Moriarty, R. D. Oeffner, B. K. Poon, M. G. Prisant, R. J. Read, J. S. Richardson, D. C. Richardson, M. D. Sammito, O. V. Sobolev, D. H. Stockwell, T. C. Terwilliger, A. G. Urzhumtsev, L. L. Videau, C. J. Williams, P. D. Adams, Macromolecular structure determination using X-rays, neutrons and electrons: recent developments in Phenix. *Acta Crystallogr Sect D*. 75, 861–877 (2019).

23. P. Emsley, B. Lohkamp, W. G. Scott, K. Cowtan, Features and development of Coot. *Acta Crystallogr Sect D*. 66, 486–501 (2010).

24. E. F. Pettersen, T. D. Goddard, C. C. Huang, E. C. Meng, G. S. Couch, T. I. Croll, J. H. Morris, T. E. Ferrin, UCSF ChimeraX : Structure visualization for researchers, educators, and developers. *Protein Sci*. 30, 70–82 (2020).

25. R. A. Laskowski, J. Jabłońska, L. Pravda, R. S. Vařeková, J. M. Thornton, PDBsum: Structural summaries of PDB entries. *Protein Sci Publ Protein Soc*. 27, 129–134 (2018).

26. J.-F. Gibrat, T. Madej, S. H. Bryant, Surprising similarities in structure comparison. *Curr Opin Struc Biol*. 6, 377–385 (1996).

27. M. J. Parker, P. R. Weigele, L. Saleh, Insights into the Biochemistry, Evolution, and Biotechnological Applications of the Ten-Eleven Translocation (TET) Enzymes. *Biochemistry-us*. 58, 450–467 (2019).

28. E. J. Burke, S. S. Rodda, S. R. Lund, Z. Sun, M. R. Zeroka, K. H. O'Toole, M. J. Parker, D. S. Doshi, C. Guan, Y.-J. Lee, N. Dai, D. M. Hough, D. A. Shnider, I. R. Corrêa, P. R. Weigele, L. Saleh, Phage-encoded ten-eleven translocation dioxygenase (TET) is active in C5-cytosine hypermodification in DNA. *Proc National Acad Sci*. 118, e2026742118 (2021).

29. S. R. Eddy, Accelerated Profile HMM Searches. *Plos Comput Biol*. 7, e1002195 (2011).

30. D. Paez-Espino, S. Roux, I.-M. A. Chen, K. Palaniappan, A. Ratner, K. Chu, M. Huntemann, T. B. K. Reddy, J. C. Pons, M. Llabrés, E. A. Eloe-Fadrosh, N. N. Ivanova, N. C. Kyrpides, IMG/VR v.2.0: an integrated data management and analysis system for cultivated and environmental viral genomes. *Nucleic Acids Res*. 47, D678–D686 (2019).

31. A. C. Gregory, A. A. Zayed, N. Conceição-Neto, B. Temperton, B. Bolduc, A. Alberti, M. Ardyna, K. Arkhipova, M. Carmichael, C. Cruaud, C. Dimier, G. Domínguez-Huerta, J. Ferland, S. Kandels, Y. Liu, C. Marec, S. Pesant, M. Picheral, S. Pisarev, J. Poulain, J.-É. Tremblay, D. Vik, T. O. Coordinators, S. G. Acinas, M. Babin, P. Bork, E. Boss, C. Bowler, G. Cochrane, C. de Vargas, M. Follows, G. Gorsky, N. Grimsley, L. Guidi, P. Hingamp, D. Iudicone, O. Jaillon, S. Kandels-Lewis, L. Karp-Boss, E. Karsenti, F. Not, H. Ogata, S. Pesant, N. Poulton, J. Raes, C. Sardet, S. Speich, L. Stemmann, M. B. Sullivan, S. Sunagawa, P. Wincker, M. Babin, C. Bowler, A. I. Culley, C. de Vargas, B. E. Dutilh, D. Iudicone, L. Karp-Boss, S. Roux, S. Sunagawa, P. Wincker, M. B. Sullivan, Marine DNA Viral Macro- and Microdiversity from Pole to Pole. *Cell*. 177, 1109-1123.e14 (2019).

32. T. Seemann, Prokka: rapid prokaryotic genome annotation. *Bioinformatics*. 30, 2068–2069 (2014).

33. A. P. Arkin, R. W. Cottingham, C. S. Henry, N. L. Harris, R. L. Stevens, S. Maslov, P. Dehal, D. Ware, F. Perez, S. Canon, M. W. Sneddon, M. L. Henderson, W. J. Riehl, D. Murphy-Olson, S. Y. Chan, R. T. Kamimura, S. Kumari, M. M. Drake, T. S. Brettin, E. M. Glass, D. Chivian, D. Gunter, D. J. Weston, B. H. Allen, J. Baumohl, A. A. Best, B. Bowen, S. E. Brenner, C. C. Bun, J.-M. Chandonia, J.-M. Chia, R. Colasanti, N. Conrad, J. J. Davis, B. H. Davison, M. DeJongh, S. Devoid, E. Dietrich, I. Dubchak, J. N. Edirisinghe, G. Fang, J. P. Faria, P. M. Frybarger, W. Gerlach, M. Gerstein, A. Greiner, J. Gurtowski, H. L. Haun, F. He, R. Jain, M. P. Joachimiak, K. P. Keegan, S. Kondo, V. Kumar, M. L. Land, F. Meyer, M. Mills, P. S. Novichkov, T. Oh, G. J. Olsen, R. Olson, B. Parrello, S. Pasternak, E. Pearson, S. S. Poon, G. A. Price, S. Ramakrishnan, P. Ranjan, P. C. Ronald, M. C. Schatz, S. M. D. Seaver, M. Shukla, R. A. Sutormin, M. H. Syed, J. Thomason, N. L. Tintle, D. Wang, F. Xia, H. Yoo, S. Yoo, D. Yu, KBase: The United States Department of Energy Systems Biology Knowledgebase. *Nat Biotechnol*. 36, 566–569 (2018).

34. S. El-Gebali, J. Mistry, A. Bateman, S. R. Eddy, A. Luciani, S. C. Potter, M. Qureshi, L. J. Richardson, G. A. Salazar, A. Smart, E. L. L. Sonnhammer, L. Hirsh, L. Paladin, D. Piovesan, S. C. E. Tosatto, R. D. Finn, The Pfam protein families database in 2019. *Nucleic Acids Res*. 47, D427–D432 (2019).

35. J. A. Gerlt, J. T. Bouvier, D. B. Davidson, H. J. Imker, B. Sadkhin, D. R. Slater, K. L. Whalen, Enzyme Function Initiative-Enzyme Similarity Tool (EFI-EST): A web tool for generating protein sequence similarity networks. *Biochimica Et Biophysica Acta Bba - Proteins Proteom*. 1854, 1019–1037 (2015).

36. K. H. O'Toole, B. Imperiali, K. N. Allen, Glycoconjugate pathway connections revealed by sequence similarity network analysis of the monotopic phosphoglycosyl transferases. *Proc National Acad Sci*. 118, e2018289118 (2021).

37. P. Shannon, A. Markiel, O. Ozier, N. S. Baliga, J. T. Wang, D. Ramage, N. Amin, B. Schwikowski, T. Ideker, Cytoscape: A Software Environment for Integrated Models of Biomolecular Interaction Networks. *Genome Res*. 13, 2498–2504 (2003).

38. D. Swinton, S. Hattman, P. F. Crain, C.-S. Cheng, D. L. Smith, J. A. McCloskey, Purification and characterization of the unusual deoxynucleoside, a-N-(9-fi-D-2'-deoxyribofuranosylpurin-6-yl)glycinamide, specified. *Proceedings of the National Academy of Science* (1983).

39. M. W. Vetting, L. P. S. de Carvalho, M. Yu, S. S. Hegde, S. Magnet, S. L. Roderick, J. S. Blanchard, Structure and functions of the GNAT superfamily of acetyltransferases. *Arch Biochem Biophys*. 433, 212–226 (2005).

40. L. Favrot, J. S. Blanchard, O. Vergnolle, Bacterial GCN5-Related N-Acetyltransferases: From Resistance to Regulation. *Biochemistry-us*. 55, 989–1002 (2016).

41. R. M. Burckhardt, J. C. Escalante-Semerena, Small-Molecule Acetylation by GCN5-Related N -Acetyltransferases in Bacteria. *Microbiol Mol Biol R*. 84 (2020), doi:10.1128/mmbr.00090-19.

42. R. Milo, P. Jorgensen, U. Moran, G. Weber, M. Springer, BioNumbers—the database of key numbers in molecular and cell biology. *Nucleic Acids Res*. 38, D750–D753 (2010).

43. L. E. Leiva, A. Pincheira, S. Elgamal, S. D. Kienast, V. Bravo, J. Leufken, D. Gutiérrez, S. A. Leidel, M. Ibba, A. Katz, Modulation of Escherichia coli Translation by the Specific Inactivation of tRNAGly Under Oxidative Stress. *Frontiers Genetics*. 11, 856 (2020).

44. T. S. STEWART, R. J. ROBERTS, J. L. STROMINGER, Novel Species of tRNA. *Nature*. 230, 36–38 (1971).

45. R. J. ROBERTS, Structures of Two Glycyl-tRNAs from Staphylococcus epidermidis. *Nat New Biology*. 237, 44–45 (1972).

46. I. Avcilar-Kucukgoze, H. Gamper, Y.-M. Hou, A. Kashina, Purification and Use of tRNA for Enzymatic Post-translational Addition of Amino Acids to Proteins. *Star Protoc*. 1, 100207 (2020).

47. G. P. Kurzban, E. A. Bayer, M. Wilchek, P. M. Horowitz, The quaternary structure of streptavidin in urea. *J Biol Chem*. 266, 14470–14477 (1991).

48. K. H. Kaminska, J. M. Bujnicki, Bacteriophage Mu Mom protein responsible for DNA modification is a new member of the acyltransferase superfamily. *Cell Cycle*. 7, 120–121 (2008).

49. R. P. Newton, E. E. Kingston, A. Overton, Identification of novel nucleotides found in the red seaweed Porphyra umbilicalis. *Rapid Commun. Mass Spectrom.* 9, 305–311 (1995).

50. D. Strzelecka, S. Chmielinski, S. Bednarek, J. Jemielity, J. Kowalska, Analysis of mononucleotides by tandem mass spectrometry: investigation of fragmentation pathways for phosphate- and ribose-modified nucleotide analogues. *Sci Rep-uk*. 7, 8931 (2017).

51. D. Fu, J. A. Calvo, L. D. Samson, Balancing repair and tolerance of DNA damage caused by alkylating agents. *Nat Rev Cancer*. 12, 104–120 (2012).

52. G.-L. Xu, M. Bochtler, Reversal of nucleobase methylation by dioxygenases. *Nat Chem Biol*. 16, 1160–1169 (2020).

53. J. B. Macon, R. Wolfenden, 1-Methyladenosine. Dimroth rearrangement and reversible reduction. *Biochemistry-us*. 7, 3453–3458 (1968).

54. A. R. Katritzky, C. A. Ramsden, J. A. Joule, V. V. Zhdankin, Handbook of Heterocyclic Chemistry (Third Edition). *Part 3 React Heterocycles*, 473–604 (2010).

55. J. T. Baumgartner, T. S. H. Mohammad, M. P. Czub, K. A. Majorek, X. Arolli, C. Variot, M. Anonick, W. Minor, M. A. Ballicora, D. P. Becker, M. L. Kuhn, Gcn5-Related N-Acetyltransferases (GNATs) With a Catalytic Serine Residue Can Play Ping-Pong Too. *Frontiers Mol Biosci*. 8, 646046 (2021).

56. M. Bochtler, H. Fernandes, DNA adenine methylation in eukaryotes: Enzymatic mark or a form of DNA damage? *Bioessays*. 43, 2000243 (2021).

57. B. Allet, A. I. Bukhari, Analysis of bacteriophage Mu and λ-Mu hybrid DNAs by specific endonucleases. *J Mol Biol*. 92, 529–540 (1975).

58. A. Toussaint, The DNA modification function of temperate phage Mu-1. *Virology*. 70, 17–27 (1976).

59. H. Khatoon, A. I. Bukhari, Bacteriophage Mu-induced modification of DNA is dependent upon a host function. *J Bacteriol*. 136, 423–428 (1978).

60. S. Hattman, Unusual Modification of Bacteriophage Mu DNA. *J Virol*. 32, 468–475 (1979).

61. M. Mirdita, K. Schütze, Y. Moriwaki, L. Heo, S. Ovchinnikov, M. Steinegger, ColabFold: making protein folding accessible to all. *Nat Methods*. 19, 679–682 (2022).

62. S. LLC, W. DeLano, *PyMOL* (2020; http://www.pymol.org/pymol).

63. L. Holm, Dali server: structural unification of protein families. *Nucleic Acids Res*. 50, W210–W215 (2022).

64. C. S. Francklyn, A. Minajigi, tRNA as an active chemical scaffold for diverse chemical transformations. *Febs Lett*. 584, 366–375 (2010).

65. E. C. Ulrich, W. A. van der Donk, Cameo appearances of aminoacyl-tRNA in natural product biosynthesis. *Curr Opin Chem Biol*. 35, 29–36 (2016).

66. Z. Su, B. Wilson, P. Kumar, A. Dutta, Noncanonical Roles of tRNAs: tRNA Fragments and Beyond. *Annu Rev Genet*. 54, 1–23 (2020).

67. C. Maruyama, Y. Hamano, tRNA-dependent amide bond–forming enzymes in peptide natural product biosynthesis. *Curr Opin Chem Biol*. 59, 164–171 (2020).

68. S. Chimnaronk, T. Suzuki, T. Manita, Y. Ikeuchi, M. Yao, T. Suzuki, I. Tanaka, RNA helicase module in an acetyltransferase that modifies a specific tRNA anticodon. *Embo J*. 28, 1362–1373 (2009).

69. K. L. Hentchel, J. C. Escalante-Semerena, Acylation of Biomolecules in Prokaryotes: a Widespread Strategy for the Control of Biological Function and Metabolic Stress. *Microbiol Mol Biol R*. 79, 321–346 (2015).

70. Y. Yashiro, Y. Sakaguchi, T. Suzuki, K. Tomita, Mechanism of aminoacyl-tRNA acetylation by an aminoacyl-tRNA acetyltransferase AtaT from enterohemorrhagic E. coli. *Nat Commun*. 11, 5438 (2020).

71. Y. Yashiro, C. Zhang, Y. Sakaguchi, T. Suzuki, K. Tomita, Molecular basis of glycyl-tRNAGly acetylation by TacT from Salmonella Typhimurium. *Cell Reports*. 37, 110130 (2021).

72. M. A. Skiba, C. L. Tran, Q. Dan, A. P. Sikkema, Z. Klaver, W. H. Gerwick, D. H. Sherman, J. L. Smith, Repurposing the GNAT Fold in the Initiation of Polyketide Biosynthesis. *Structure*. 28, 63-74.e4 (2020).
